# Neural correlates of transitions between internal and external cognitive domains

**DOI:** 10.1101/2025.10.27.683183

**Authors:** Risa Dwi Ratnasari, Paul Zen Cheng, Tzu-Yu Hsu

## Abstract

Human cognition involves continuous transitions between processing information guided by the external world and engaging with internally-generated thoughts, yet research has predominantly investigated switching based on external rules. We therefore examined neural mechanisms underlying external and internal domain switching using independent groups of EEG (N=46) and fMRI (N=60) in a colour judgement task requiring evaluations based on either colour similarity (external domain) or colour preference (internal domain) through switching on a trial-by-trial basis. ERPs revealed amplitude differences between the two judgements as well as switching and repeated trials. EEG analysis revealed a sustained neural differentiation between switch and repeated trials, specifically during similarity judgements, with effects lasting from early time points through the entire trial duration. fMRI showed similarity judgements engaged visual and motor regions during switching, while preference judgements showed minimal subcortical activation for repeated trials. These findings demonstrate that external and internal cognitive domain switching effects are asymmetrical. These findings provide the first temporal and spatial neural characterisation of domain switching between processing information guided by the external world and internally-generated thoughts, indicating that shifting between internal and external domains involves mechanisms distinct from traditional external rule-switching paradigms.

## Introduction

Human cognition involves continuous transitions between processing information directly mediated by the external world and engaging with internally-generated mental content. When reading a novel, for instance, our brain is constantly switching information from both the physical text of the book (i.e. external domain) and the ongoing narrative in our mind (i.e. internal domain). Although we experience these transitions as subjectively fluid, limited work has been done to investigate the nature of the processes involved and whether transitions in each direction are mechanistically symmetrical. Similarly, the neural correlates of switches between external and internal domains remain rarely characterised.

Extensive research has focused on task switching in paradigms requiring participants to alternate between distinct rules (Chiu & Yantis, 2009; Kim et al., 2012; Ravizza & Carter, 2008; Vandierendonck et al., 2010), such as shape versus letter identification. These studies consistently demonstrate “switch costs”, with increasing reaction times (RTs) or error responses when transitioning between different task rules compared to repeating the same rule on a trial-by-trial basis (Koch et al., 2018). Such cost reflects cognitive demands associated with reconfiguring task sets while inhibiting competing alternatives (Dove et al., 2000). This phenomenon has been replicated across diverse stimuli and research domains (Samson et al., 2022), establishing a well-characterised neural circuit involving several brain regions within the frontoparietal network (Chiu & Yantis, 2009; Cools et al., 2004; Esterman et al., 2009; Periáñez et al., 2024; Worringer et al., 2019).

As well as work investigating rule-switching, a growing number of studies have begun to explore switch costs between internal and external domains under memory-based (i.e. internal domain) and sensory-based (i.e. external domain) tasks (Dark, 1990; Gresch et al., 2024; Hautekiet et al., 2023; Verschooren, Schindler, et al., 2019; Verschooren et al., 2020, 2025). These demonstrate that domain switches induce RTs costs similar to those related to rule switching. Importantly, some behavioural studies also demonstrated asymmetrical switch costs, with longer RTs being observed when switching from the external to the internal task but not in reverse (Dark, 1990; Verschooren et al., 2025; Verschooren, Liefooghe, et al., 2019, 2019), although some studies have not seen this asymmetry (Calzolari et al., 2022; Gresch et al., 2024; Hautekiet et al., 2023). Calzolari et al. (2022) further extended the internal and external domain-switching phenomenon beyond memory modality to self-referential processing. They compared switches between internally-oriented tasks (i.e. personality or bodily sensation assessments) and externally-oriented tasks (i.e. letter-based judgements) and found that switch cost occurs not only between tasks within each domain but also between domains. These findings suggest that internal-external domain switching may be associated with different cognitive operations, yet its underlying neural mechanisms remain largely uncharacterized.

Despite its importance to ongoing cognitive processes, the neural basis of domain switching has received limited attention. Gilbert et al. (2005) provided early evidence that transitions between stimulus-oriented thought (external domain) and self-generated information independent from surrounding sensory stimuli (internal domain) engage the right anterior mid-dorsolateral prefrontal cortex, right rostrolateral prefrontal cortex, and bilateral superior parietal cortex with a small number of participants. Complementing this work, Kam et al. (2018) demonstrated that patients with lateral prefrontal cortex lesions exhibit impaired coordination between externally and internally directed attention tasks. While these studies offer valuable insights, neural networks underlying internal-external domain switching remain not fully explored. The present study addresses these limitations by investigating the neural mechanisms that govern transitions between self-initiated internal cognition and visually-guided external processing.

To investigate the temporal and spatial neural mechanisms underlying domain switching, we adapted a colour judgement paradigm originally developed by ( Johnson et al., 2005) that operationalises the external-internal distinction through objective versus subjective evaluation criteria. Participants performed colour similarity judgements (external domain: “Which colour patch more closely matches the target?”) and colour preference judgements (internal domain: “Which pair of colours do you prefer?”) to probe external and internal domains, respectively (Figure 1A, 1E). These two judgements were varied on a trial-by-trial basis with an equivalent number of repeating and switch trials. With both judgement types involving similar visual stimuli and motor responses, this design allows the examination of switching between the different cognitive evaluation modes of interest while controlling for basic perceptual and motor demands. We conducted complementary EEG and fMRI experiments using two independent participant groups (except five participants participating in both experiments) to characterise both the temporal dynamics and spatial networks involved in domain switching.

**Figure 1.**
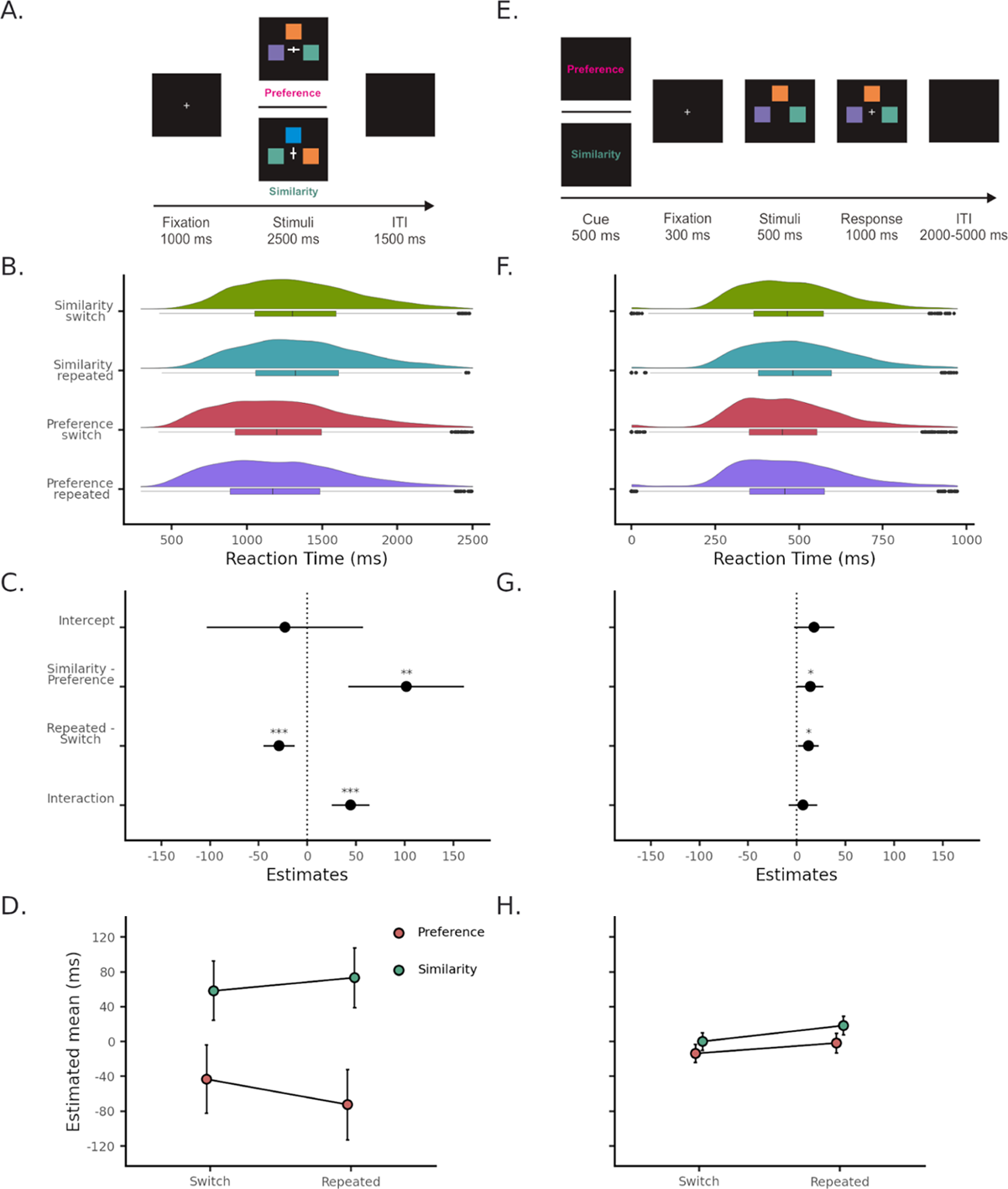
The design of the colour judgement task in EEG and fMRI experiments, and the LME model estimated behavioural results. (A) Experimental procedure for the EEG experiment. Participants performed colour judgements based on cue type indicated by fixation cross orientation: vertical elongation signalled the similarity judgement, while horizontal elongation signalled the preference judgement. Meanwhile, participants viewed three colour patches and made their judgement via button press within 2500 ms. (B) Behavioral performance in the EEG experiment. Density and box plots show RT distributions for each judgement type (Preference and Similarity) and condition (Switch and Repeated). (C) EEG LME model intercept and fixed effects coefficient of interest. Each estimated fixed term is presented, with error bars as confidence intervals. Significant level is indicated with *p < 0.05, **p < 0.01, ***p< 0.001. The model shows a strong interaction between Judgement (preference vs. similarity) and Condition (switch vs. repeated), demonstrating differential switching costs across domains. (D) EEG LME model’s Judgement to Condition type estimated mean RTs with estimated standard error of mean. (E) Experimental procedure for the fMRI experiment. At the beginning of each trial, explicit text cues (“Similarity” or “Preference”) indicate the required judgement types for the upcoming trial, followed by the same colour judgement task as in (A). A notable difference between the fMRI and EEG paradigms is that the judgement type cue and preparation duration to make responses are different within the trial. (F)Behavioral performance in the fMRI experiment. Density and box plots show RT distributions for each judgement type and condition. (G) fMRI’s LME model intercept and fixed effects coefficient of interest. Each estimated fixed term is presented with error bars as confidence intervals. Significant level is indicated with *p < 0.05, **p < 0.01, ***p< 0.001. The model shows a negligible interaction effect and weak main effects of Judgement and Condition. (H) fMRI LME model’s Judgement to Condition type estimated mean RTs with estimated standard error of mean.

## Results

### Behavioral Performance for External and Internal Domain Switching

We used linear mixed-effects (LME) models to examine whether *Judgement* (colour preference vs. colour similarity judgement) and *Condition* (switch vs. repeated trial) influenced RTs. All models included random intercepts for participants, with additional fixed and random effects for block and trial sequence determined through model comparison procedures (see Methods and Supplementary 1-2). The model structures reported here represent optimal configurations identified via likelihood ratio tests during model selection, conducted separately for EEG and fMRI datasets. Figure 1B and 1F display RTs as a function of judgement type and condition for both experiments (Supplementary 1.4 & 2.4 for quantiles). Detailed demographic information is provided in Table 1.

**Table 1.** Demographic information in EEG and fMRI experiments.

| Variable |  | EEG |  |  |  | fMRI |  |  |  |
| --- | --- | --- | --- | --- | --- | --- | --- | --- | --- |
|  |  | *n* | % | *M* | *SD* | *n* | % | *M* | *SD* |
| Age |  |  |  | 27.6 | 4.9 |  |  | 27.8 | 4.9 |
| Sex | Female | 25 | 53.2 |  |  | 32 | 53.3 |  |  |
|  | Male | 22 | 46.8 |  |  | 28 | 46.7 |  |  |
| Total |  | 47 |  |  |  | 60 |  |  |  |

#### EEG Behavioural Results

The model is structured as follows: *RTs ~ Judgement × Condition + Blocks + (Judgement + Condition | Participants)* (see Supplementary Table 1.2 for interaction model effect table). This model revealed fixed effects of *Judgement*, *Condition* and their interaction. Colour similarity judgements were estimated to be 101.51 milliseconds (ms) slower than colour preference judgements (*SE* = 29.46, *t*(47.66) = 3.44, *p* < 0.01, CI = [42.25, 160.76]; Figure 1C). Condition types have estimated differences where repeated trials are estimated to be 29.16 ms faster than the switch trials (*SE* = 8.12, *t*(95.24) = −3.59, *p* < 0.001, CI = [−45.29, −13.04]; Figure 1C). An interaction effect was also found, indicating switch cost difference between judgements (*β =* 44.28, *SE* = 9.85, *t*(16.40) = 4.95, *p* < 0.001, CI = [24.98, 63.60]; Figure 1C and 1D). Following the interaction result, we conducted post hoc pairwise comparison on the interaction model to test the switch cost across Judgement types. We found switch cost effect with preference judgement where preference switch trials estimated 29.14 ms slower than preference repeated trials (*SE* = 8.12, *z* = −3.59, *p* < 0.001), and negligible varied switch cost effect with similarity judgement where similarity’s switch trial estimated 15.12 ms faster than repeated trials (SE = 8.59, z = −1.75, p = 0.078). This suggests that switching costs are only observed in preference judgement but not similarity judgement.

#### fMRI Behavioural Results

To minimize motor response-related confounds in BOLD signal estimation, we deliberately implemented a delayed response array in the fMRI study and explicitly instructed participants not to respond quickly. Therefore, we did not expect to observe switch costs. Indeed, no interaction between *Judgement* and *Condition* was obtained (*β =* 6.28, t(63.92) = 0.86, *p* = 0.39, CI = [−8.25, 20.80]; Figure 1G). We observe a weak difference between colour judgement types: the estimated coefficient in the similarity judgement were 13.81 ms slower than the preference judgement (SE = 6.80, t(87.95) = 2.03, *p* < 0.05, CI = [0.30, 27.33]; Figure 1H), suggesting that external domain processing requires more time than internal domain processing. Interestingly, we found a weak difference between condition types that contradicted our EEG LME model—the repeated condition is estimated 11.95 ms slower than the switch condition (SE = 5.35, t(63.44) = 2.23, *p* < 0.05, CI = [1.26, 22.65]; Figure 1H).

The behavioural results demonstrate that the similarity judgement consistently requires longer processing times than the preference judgement. Importantly, the behavioural results in the EEG experiment revealed asymmetric switching costs when the cue and stimuli were presented simultaneously, with clear costs for the switch trials within the preference judgement but not within the similarity judgement. As expected, no switch cost was observed in the fMRI behavioural results due to the experimental design customised for fMRI experiment.

### Distinct Brain Activity Pattern for External and Internal Domain Processing

To examine whether internal and external domain processing engage distinct neural activity patterns, we conducted comprehensive analyses comparing the preference judgement (internal domain) and the similarity judgement (external domain). Our analytical approach included univariate event-related potentials (ERPs), time-resolved multivariate pattern analysis (MVPA), temporal generalisation analysis, and univariate fMRI.

#### EEG Reveals Early and Sustained Domain Differences

Univariate ERP analysis revealed an early post-stimulus component from 155 to 290 ms with significantly higher amplitudes for similarity judgements compared to preference judgements (cluster p < 0.05). This effect was broadly distributed across channels, suggesting widespread cortical differences between external and internal domain processing (Figure 2A).

**Figure 2.**
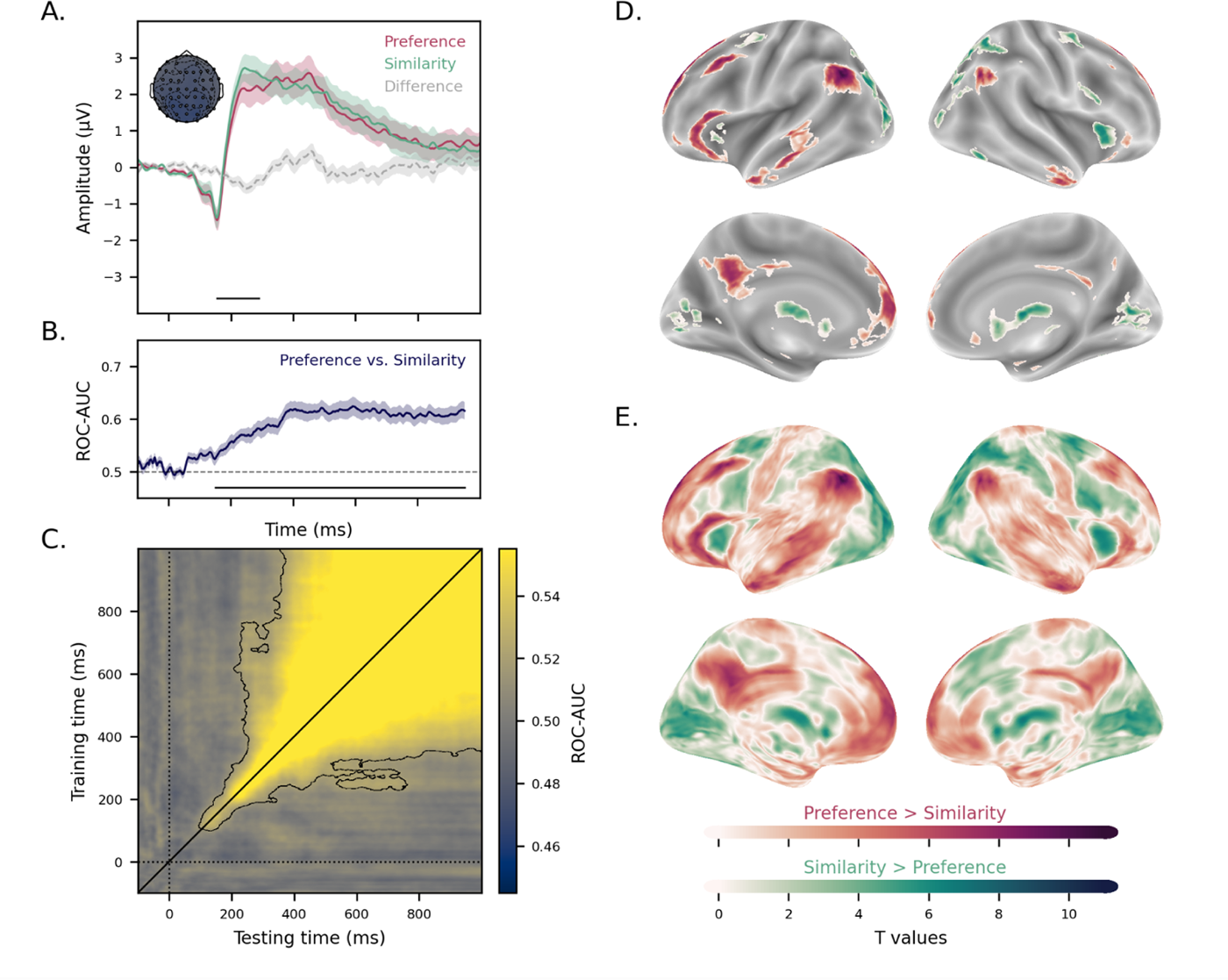
Distinct brain activity patterns for external and internal domain processing. (A) Grand averaged ERPs for colour preference (red) and colour similarity (green) judgements. The horizontal bar indicates the significant time window (155-290 ms) where the similarity judgement showed greater amplitude than the preference judgement amplitude. This significant cluster component has widespread scalp distributions displayed in topographical maps in the left-hand corner. Significant channels in this component are highlighted in bold circles, while non-significant electrodes are displayed as empty circles in the topography. (B) Time-resolved multivariate decoding classification between preference and similarity judgements. The classifier performed significantly above chance in discriminating capability to distinguish domains in post-stimulus intervals between 151 to 951 ms (cluster p < 0.001). The horizontal bar denotes significant time periods. This finding indicates persistent domain-specific neural representations. (C) Temporal generalisation analysis showing how domain-specific neural representations evolve over time. The matrix displays decoding performance when training classifiers at one time point and testing at another time point. Strong diagonal performance (100-400 ms) indicates temporally-specific representations, while the sustained square pattern (>400 ms) reflects temporally-generalised representations that persist across time. (D & E) Comparison of BOLD signal between preference and similarity judgements. Internal domain (preference > similarity) indicated by warm colour palette (red) and external domain (similarity> preference) indicated by cold colour palette (green). (D) Greater activations in self-referential regions (i.e., precuneus and frontal regions) with the internal domain and greater activation with the external domain in perceptual-related brain regions (i.e., occipital and caudate regions). The T-value maps are thresholded with P_fwe_ < 0.05 across two contrasts. (E) Unthresholded versions of the internal and external domain contrasts map.

To determine the temporal dynamics of domain-specific neural activity, we employed a time-resolved cross-validation decoding approach. We trained a linear discriminant analysis (LDA) classifier to discriminate between Judgement types at each time point, using the Area Under the Curve of the Receiver Operating Characteristic (ROC-AUC) as the performance metric. Results revealed successful decoding from 151 to 951 ms post-stimulus onset (cluster p < 0.001; Figure 2B), indicating that domain-specific neural activity emerges shortly after stimulus onset and persists throughout most of the trial period.

To examine the stability of these neural representations over time, we conducted temporal generalisation analysis. The LDA classifier was trained at each time point and tested against all other time points. This cross-temporal decoding revealed a complex pattern: transient neural activity along the diagonal from 100 to 400 ms post-stimulus, followed by a square-shaped sustained activity pattern extending beyond 400 ms (cluster p < 0.001; Figure 2C). This temporal structure suggests that domain discrimination may involve two distinct processing phases—an initial transient phase reflecting rapid stimulus categorisation (100-400 ms), followed by a sustained representational phase supporting ongoing judgement processes (> 400 ms).

#### fMRI Reveals Distinct Networks for Internal and External Domains

The fMRI results were analysed using a General Linear Model (GLM) approach. We conducted permutation tests to compare beta maps between colour preference judgements and colour similarity judgements (see Methods and Supplementary 12 for detailed first-level HRF modelling and preprocessing procedures). Statistical significance was determined using a threshold of p < 0.05 with family-wise error (FWE) corrected voxel-wise and within model contrast-wise. To provide a comprehensive view of regional activation differences, we present both thresholded statistical maps and unthresholded statistical maps in Figure 2E, with the latter highlighting the full spatial extent of domain-specific activation patterns (Chen et al., 2022).

We found differences in BOLD activity from internal to external domains (i.e., preference > similarity) throughout the frontal, parietal, and temporal lobes and cerebellum (Figure 2D, red regions; Supplementary Table 3.1; Supplementary 5 for fMRI results from cue onset). Notably, many of the highlighted regions are constituent parts of the default-mode network (DMN), including the posterior parietal cortex (PCC), dorsal prefrontal cortex (DPFC) and anteromedian prefrontal cortex (AMPFC) (Alves et al., 2019). This network has been associated with self-referential, subjective evaluation, and internally-oriented cognitive operations (Levorsen et al., 2025; Nakao et al., 2012).

The contrast of the external with internal domain (similarity > preference) showed increased activation in regions associated with perceptual processing and externally oriented processes (Figure 2D, green regions), including the bilateral middle occipital gyrus, bilateral caudate nucleus, bilateral calcarine fissure and surrounding cortex, right fusiform gyrus, bilateral superior parietal gyrus, right insula. Also, other central regions include the bilateral precentral gyrus and superior frontal gyrus, and the right inferior temporal gyrus in the temporal region. Last, the hindbrain regions, vermis and bilateral cerebellum (Figure 2D, green regions; Supplementary Table 3.2). This pattern may reflect enhanced sensory processing mechanisms during objective perceptual comparisons.

Although we deliberately implemented a delayed response array designed in the fMRI experiment, we still conducted additional control analysis using GLM with the original parameters above and added RTs duration (RTDUR) as a covariate (Mumford et al., 2024; Taylor et al., 2014). Overall, similarity judgement contrast with RTDUR has retained it but preference judgement contrasted with RTDUR has decreased task-related pattern (See supplementary Figure 4.1 and Table 4.1 for figures and the ROI table supplementary Figure 4.2-3 and Table 4.2-3).

These converging EEG and fMRI findings support our hypothesis that external and internal domains engage distinct neural networks. The EEG results reveal early differentiation (within 150-200 ms) that persists throughout processing, while fMRI demonstrates anatomically segregated networks, with similarity judgements activating perceptual regions and preference judgements engaging self-referential areas. Together, these findings establish that external-internal distinctions are fundamental to neural organisation, manifesting across both temporal and spatial dimensions of brain activity.

### Domain-Independent Task Switching Effects Show Minimal Activity Differentiation

To investigate whether switch versus repeated trials engage common mechanisms regardless of judgement type, we analysed activity patterns associated with task switching by collapsing across similarity and preference judgements. We applied the same analytical approaches as described previously—univariate ERPs, MVPA, temporal generalisation analysis, and univariate fMRI—but focused specifically on the condition effect (switch vs. repeated trials) independent of judgement type.

#### Sustained But Weak Signatures of Task Switching Independent of Domain Type

We found a significant ERP component starting at 324 to 964 ms (cluster *p* < 0.001) with higher amplitudes for switch condition compared to repeated condition. This effect was distributed across central to posterior electrodes (Figure 3A), indicating that neural activity around the posterior — central region contributes to the switching performance during the colour judgement task.

**Figure 3.**
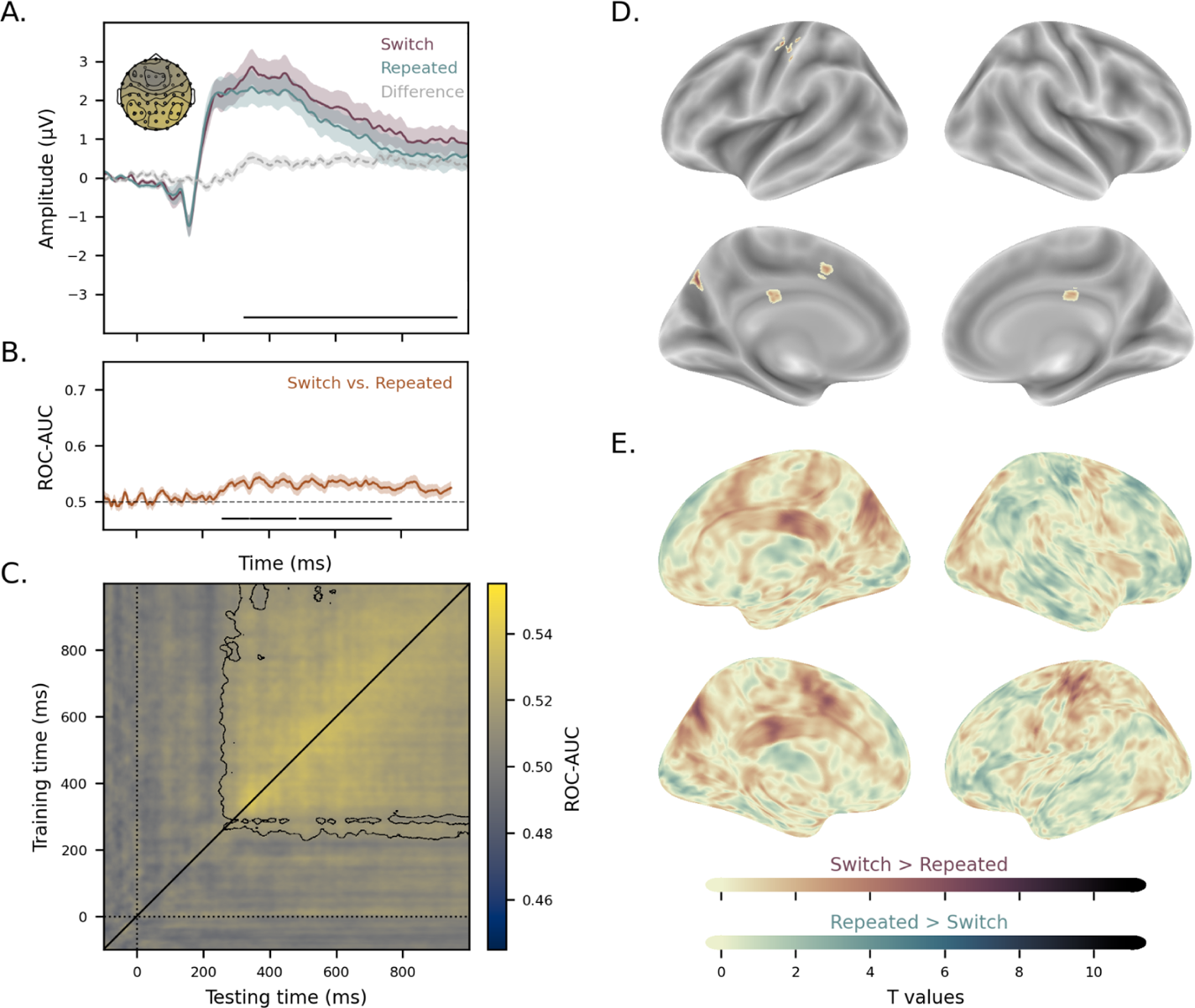
Brain activity patterns for switch and repeated trials. (A) Grand average event-related potentials (ERPs) for switch (mauve) and repeated (turquoise) conditions. The horizontal bar indicates the significant time window where switch judgements showed greater amplitude than repeated conditions. Topographic maps display the distribution of neural activity from central to posterior across the significant cluster. Electrodes contributing to the significant cluster are shown in bold circles, while non-significant electrodes are indicated with empty circles. (B) Time-resolved multivariate decoding accuracy (ROC-AUC) for discriminating between switch and repeated conditions. The horizontal bar denotes significant time periods. (C) Temporal generalisation analysis showing how condition-specific neural representations evolve over time. (D & E) Comparison of BOLD activations between the switch and repeated trials conditions. Switch > repeated trials contrast indicated by shade of reddish brown colour palette (mauve) and repeated > switch trials contrast indicated by shades of ocean blue colour palette (turquoise). (D) Greater activations in the left precentral and precuneus cortex in the switch > repeated trials condition, which indicates higher visual and motor demand in the switch condition. The T-value maps are thresholded with P_fwe_ < 0.05 across two contrasts. (E) Unthresholded versions of the contrasts map to help visualise the extended network.

The time-resolved multivariate decoding was performed using LDA to train the classifier to distinguish different conditions between switch and repeated trials. The result showed three significant clusters starting from 260 ms - 338 ms (cluster *p* < 0.05), 345 ms - 480 ms (cluster *p* <0.05), and 493 ms - 769 ms (cluster *p* < 0.01).

A cross-validation analysis of the Condition effect was then conducted. As expected, the classifier detected sustained generalizability of the neural representation of the switching effect. A square-shaped pattern spanning from 250 to the end of the temporal generalisation window at 951 ms can be seen (cluster *p* < 0.001; Figure 3C). This pattern generalisation indicates that the neural representations of switch and repeated trial type were maintained throughout stimulus presentation.

#### fMRI Reveals Minimal Neural Activation Differences Between Switch and Repeated Trials

Switch versus repeated conditions were then compared in the fMRI data. Switch trials showed higher activation than repeated trials in left pre-postcentral and occipital-precuneus regions (Figure 3D, mauve regions; Supplementary Table 3.2), while no significant differences were observed for the reverse contrast. Similar results were also found from cue onset (Supplementary 7 for fMRI results from cue onset), suggesting the onset time does not affect the results. From the unthresholded brain maps presented in Figure 3E, compared switch to repeated repeated patterns extend voxel pattern from pre-postcentral regions, occipital-precuneus and middle cingulate & paracingulate to supplementary motor area. The switch trial effect did not survive when controlling for RTDUR (see Supplementary 6.1-6.2 for corrected and unthresholded maps and table).

The minimal differences between switch and repeated trials in both ERP and fMRI analyses provide limited evidence for general task switching mechanisms. These effects were weak compared to the robust domain-specific differences between similarity and preference judgements.

### Brain Activity Patterns for Switching vs. Repeated Trials Under External Domain Processing

The weak differences between switch and repeated trials in the previous analysis may reflect different switching mechanisms depending on whether the switch is from internal to external domains or external to internal. To test this possibility, we conducted separate analyses of the condition effect within each judgement type (see Methods). We hypothesised that switch-related neural patterns would be modulated by the specific judgement context. Using the same analytical approach as before—ERP and decoding analyses for EEG data, and univariate analysis for fMRI data—we first examined switch versus repeated trials specifically within the similarity judgement.

#### EEG Reveals Robust Switch-Repeat Distinctions During the Similarity Judgement

We first examined whether switch and repeated trials during similarity judgements produced distinguishable evoked potentials. ERP analysis revealed that switch trials elicited greater amplitude responses than repeated trials from 255 to 1000 ms post-stimulus (cluster *p* < 0.001, Figure 4A), with this effect broadly distributed across all scalp channels.

**Figure 4.**
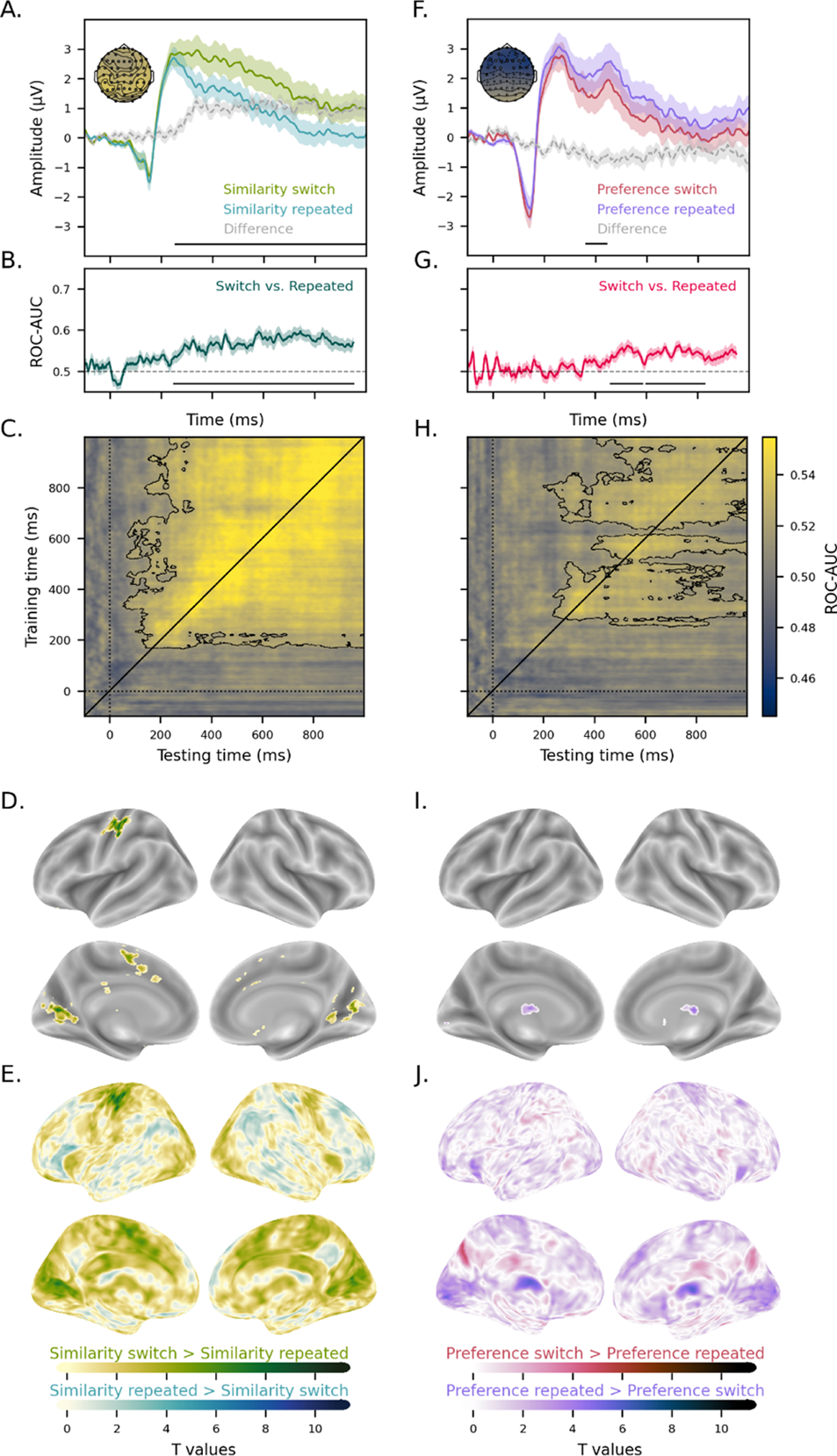
Distinct Switching Effects Between External and Internal Domains. (A-E) Neural patterns for switching vs. repeated trials under external domain processing. (A) Grand average ERPs comparing similarity switch (lime green) and similarity repeated (cyan) trials. Similarity switch trials showed significantly greater amplitude than similarity repeated trials during the 255-1000 ms time window (indicated by horizontal bar). The topographical map (top left) displays the widespread scalp distribution of this effect, with significant electrodes indicated by bolded circles. (B) Time-resolved multivariate decoding results for discriminating between switch and repeated trials within similarity judgements. Decoding accuracy reached above-chance levels from 250 ms post-stimulus and persisted until 951 ms (horizontal bar indicates significant periods, cluster p < 0.001), demonstrating sustained condition-specific neural representations throughout this extended time window. (C) Temporal generalisation analysis reveals that condition-specific neural representations evolve over time during the similarity judgement. The square pattern emerging after 156 ms reflects temporally generalised representations that persist across time, indicating stable and sustained neural signatures of switching effects within the similarity judgement condition. (D & E) Comparison of BOLD activations in the external domain’s task switching. Similarity-switch> similarity-repeated contrast indicated by a shades of yellowish green colour palette (lime green) and similarity-repeated > similarity-switch contrast indicated by a shades of greenish-blue palette (cyan). (D) Multiple regional activations are associated with visual processing and motor control in the similarity-switch > similarity-repeated contrast, including the postcentral gyrus, supplementary motor area, calcarine, caudate, putamen, lingual and cingulate cortex. The T-value maps are thresholded with P_fwe_ < 0.05 across two contrasts. (E) Unthreshold versions of the contrasts map to help visualise the extended network. The T-value maps are thresholded with P_unthreshold_ < 0.01. (F-J) Neural patterns for switching vs. repeated trials under internal domain processing (F) Grand average ERPs comparing switch (red) and repeated (purple) conditions during the preference judgement. Repeated trials showed significantly greater amplitude than switch trials during a brief time window (364-444 ms, indicated by horizontal bar). The topographic map displays the frontal-central distribution of this effect, with electrodes contributing to the significant cluster highlighted in bold circles and non-significant electrodes shown as empty circles. (G) Time-resolved multivariate decoding accuracy (ROC-AUC) for discriminating between switch and repeated conditions during preference judgements. Above-chance decoding emerged in two distinct clusters: an early cluster from 460-583 ms post-stimulus (p < 0.01) and a later cluster from 598-826 ms (p < 0.001), indicated by horizontal bars. This pattern suggests late-onset and relatively weak condition-specific neural representations compared to similarity judgements. (H) Temporal generalisation analysis showing the evolution of condition-specific neural representations during preference judgements. The weak square pattern emerging after 260 ms reflects limited temporally-generalised representations, indicating minimal switching effects within the preference judgement condition compared to the robust effects observed during similarity judgements. (I & J) Comparison of BOLD activations in the internal domain’s task switching. Preference switch > preference repeated indicated by a shades of brown colour palette (red) and preference repeated > preference switch indicated by a shades of purple palette (purple). (I) Weak regional activation in subcortical regions in the preference repeated > preference switch condition, including right cerebellum, right caudate and bilateral thalamus. The T-value maps are thresholded with P_fwe_ < 0.05 across two contrasts. (J) Unthresholded versions of the contrasts map to help visualise the extended network.

Following the ERP, time-resolved multivariate decoding successfully discriminated between switch and repeated trials from 250 to 951 ms (cluster *p* <0.001; Figure 4B). This indicates that neural representations distinguishing switch from repeated trials emerge early after stimulus onset and are maintained throughout similarity judgement processing. The temporal generalisation analysis further supported these findings, showing a square-shaped sustained activity pattern of above-chance decoding beginning at 156 ms (cluster *p* < 0.001; Figure 4C).

#### Switch Trials in the Similarity Judgement Activate Visual and Motor Networks

Higher activation in regions associated with visual processing and motor control was found in switch compared to repeated trials during similarity judgements (Figure 4D, lime green; Supplementary Table 3.4; Supplementary 9 for fMRI results from cue onset). Conversely, no difference was found in the repeated versus switch trial contrast. Notably, traditional switching-related brain regions did not show significant differences in either contrast. We also present unthresholded brain maps in Figure 4E, and they present repeated to switch effects with a minimal effect in the left triangular part of inferior frontal gyrus. In the control analysis with RTDUR, the motor process in the left pre-postcentral gyrus survived voxel-wise FWE correction, suggesting external domain switching may require higher motor and sensory demand (see supplementary Figure 8.3 and Table 8.3).

### Brain Activity Patterns for Switching vs. Repeated Trials *Under Internal Domain Processing*

Similar analyses were conducted comparing switch and repeated trials under the preference judgement only to examine whether switching from external to the internal domains involves different neural mechanisms.

#### EEG Reveals Weak and Brief Switching-Repeated Distinctions During the Preference Judgement

We performed univariate ERP analysis on preference switch and repeated trials and found a late onset and brief duration of differentiation, with significant differences emerging only between 364-444 ms post-stimulus. Notably, repeated trials produced higher amplitudes than switch trials (cluster p < 0.05; Figure 4F). Topographical results revealed that switching-related neural activity during preference judgements was primarily localised to frontal and central electrodes.

Multivariate analysis of the segmented signal revealed two significant clusters of discrimination: an early cluster from 460-583 ms (cluster p < 0.01) and a later cluster from 598-826 ms (cluster p < 0.001; Figure 4G). The overall decoding performance indicated detectable switching signatures within preference judgements.

Temporal generalisation analysis showed significant sustained decoding performance for switch versus repeated trials within the preference condition, though the decoding was weak and emerged two clusters: 265-583 ms and 608-899 ms (cluster p <0.01; Figure 4H). This pattern suggests that neural representations underlying switching within preference judgements are relatively similar to each other, indicating that preference judgements may rely on consistent neural mechanisms that are less strongly modulated by switching demands.

#### fMRI Shows Minimal Subcortical Activation for Repeated vs. Switch in the Preference Judgement

Higher activation in subcortical regions was found with the repeated trials compared to switch trials during preference judgements. Although these finding clusters were small, repeated trials found higher activation in the right cerebellum, right caudate nucleus, and left thalamus region (Figure 4I, purple regions; Supplementary Table 3.3; unthresholded maps in Figure 4J; Supplementary 9 for fMRI results from cue onset). No effect survived for the switch compared to repeated trials during preference judgements, nor voxel-wise FWE correction when controlling for RTDUR (see Supplementary 8.1). To note that the unthresholded maps (Figure 4J, red regions) have presented a similar pattern as in occipital-precuneus and middle cingulate and paracingulate to supplementary motor area, which was found in Condition contrast between switch to repeated trials (Figure 3D), and this effect can also be observed in the unthresholded maps contrast controlled with RTDUR as well (see Supplementary 8.2). Taking it together suggests that reaction time variance may account for the observed activation patterns.

EEG analysis revealed robust and sustained neural differentiation between switch and repeated trials, specifically during similarity judgements, with effects lasting from early time points through the entire trial duration. In contrast, fMRI showed modest activation differences: similarity judgements engaged visual-motor regions during switching, while preference judgements showed minimal subcortical activation for repeated trials. These findings demonstrate that task switching effects are most pronounced within the external domain (similarity judgements), with limited neural signatures during internal domain (preference) processing.

## Discussion

We examined the neural mechanisms underlying transitions between internal and external cognitive domains using complementary EEG and fMRI approaches. Our results across both neuroimaging modalities revealed a consistent asymmetrical transition pattern: shifting from internal processing (preference judgements) to external processing (similarity judgements) produced robust neural activation and enhanced decoding accuracy, while the reverse transition from external to internal processing showed minimal differential effects. This asymmetry implicates distinct neural substrates underlying different directions of cognitive domain transition.

### Domain-Specific Neural Networks Underlying Internal and External Cognitive Processes

Similarity and preference judgements differentiated at 155-290 ms post-stimulus, with similarity judgements eliciting higher amplitudes (Figure 2A-B). These findings align with Verschooren et al. (2025), who reported larger P1, N1, and P2 amplitudes for external relative to internal processing, indicating that domain-specific neural mechanisms diverge during perceptual and early post-perceptual stages. Temporal generalization analysis revealed that neural patterns evolved from transient, time-specific representations during early processing to sustained, broadly distributed patterns during later processing (Figure 2C), suggesting multiple processing stages are engaged across cognitive domains.

Complementary fMRI results demonstrated distinct spatial networks for each domain (Figure 2D-E). Similarity judgements elicited higher activity than preference judgements in bilateral occipital cortex, right inferior parietal cortex, and right frontal cortex—regions associated with sensory processing and external cognition. Conversely, preference judgements induced higher activity in default mode network regions associated with internal processes including mind-wandering (Christoff et al., 2016; Mason et al., 2007; Smallwood & Schooler, 2015), autobiographical memory (Spreng et al., 2009), and self-referential processing (Andrews-Hanna et al., 2010, 2014; Johnson et al., 2005; Northoff & Bermpohl, 2004; Schacter et al., 2012). The converging EEG and fMRI evidence indicates that external and internal processing engage dissociable neural networks that diverge within the first 300 ms and are maintained through distinct distributed cortical systems.

### Does the frontoparietal network serve as a superordinate controller in domain switching?

A superordinate control mechanism has been proposed as one of the mechanisms to regulate transitions between external and internal domains (Gresch et al., 2024; Nobre & Gresch, 2025). The frontoparietal network consistently implicated in attentional shifts and task switching (Chiu & Yantis, 2009; Chuikova et al., 2025; Gilbert et al., 2005; Monsell, 2003; Nobre & Gresch, 2025) and show enhanced activity when contrasting switching versus repeated conditions regardless of transition direction. However, we did not observe robust frontoparietal network engagement. When collapsing across both transition directions, we found higher activities in precuneus, occipital cortex, and pre- and post-central gyrus (Figure 3D), with only weak frontoparietal activation in unthresholded statistical maps that did not survive correction (Figure 3E). Temporal generalization analysis revealed sustained but weak patterns during late time windows (Figure 3C), further suggesting that broadly distributed control networks were not prominently engaged during domain switching.

In the direction-specific analysis, i.e. internal-to-external and external-to-internal transitions, we still did not observe frontoparietal network activity in either direction. Internal-to-external transitions (similarity switch vs. repeated) recruited primary sensory cortex and subcortical regions including visual cortex, motor cortex, thalamus, caudate, and cerebellum (Figure 4D-E), while external-to-internal transitions (preference switch vs. repeated) showed minimal differential activation (Figure 4I-J). Complementary EEG evidence demonstrated higher temporal generalization decoding accuracy for similarity switches (Figure 4C) than preference switches (Figure 4H). These findings suggest that domain switching does not operate through a frontoparietal superordinate control mechanism, but rather through direction-dependent mechanisms.

One alternative interpretation is that external and internal processing compete through their respective specialized networks rather than requiring superordinate control. The default mode network is widely shown to be intrinsically anticorrelated with task-positive networks during rest and task performance (Fox et al., 2005; Fransson, 2005; Murphy & Fox, 2017), suggesting competitive rather than cooperative dynamics between internal and external processing systems. Our findings are consistent with this framework: rather than recruiting a general switching network, domain transitions engage the specific networks associated with the target domain. Internal-to-external transitions activate sensory and subcortical systems that support external processing, while external-to-internal transitions show minimal additional activation, potentially reflecting fall back to a baseline state. The DMN has been long associated with a baseline state during the rest (Poerio & Karapanagiotidis, 2025; Qin et al., 2010). When external task demands gradually diminish during external-to-internal transitions, the DMN activity increases may simply reflect it falling back to baseline. Therefore, no extra cost is needed.

The subcortical regions we observed, particularly thalamus and caudate, showed anatomical connectivity patterns that could support large-scale network coordination and state transitions (Basile et al., 2021; Lam et al., 2025; Lehéricy et al., 2004; Lyu et al., 2025; Shine et al., 2023; Sydnor et al., 2025; Whyte et al., 2024). Clinical evidence demonstrates that modulating subcortical activity influences the balance between internal and external processing: deep brain stimulation can restore sensory responsiveness in disorders of consciousness (Chudy et al., 2023) and reduce excessive internal focus in major depressive disorder (K. A. Johnson et al., 2024). These anatomical and clinical findings support the possibility that domain transitions occur through competitive network dynamics rather than superordinate control.

However, several limitations should be acknowledged. We did not manipulate attention explicitly in our study; rather, we designed our paradigm to simulate naturalistic transitions between internal and external processing. We cannot exclude the possibility that both superordinate control and competitive network mechanisms exist, with the brain flexibly adopting one strategy depending on task demands or context. Our experimental design may have favored anticorrelated dynamics over top-down control, and different task structures—such as those with explicit preparatory cues or higher cognitive demands—might reveal superordinate control mechanisms. Future research systematically manipulating control demands and transition predictability could clarify when and how different coordination mechanisms are engaged during domain switching.

### Limitations

Future research should explore domain switching using no-report paradigms to isolate neural correlates without response-related confounds. Approaches such as pupillometry, analysis of spontaneous eye movements, or continuous neural decoding methods could capture domain transitions without explicit behavioral reports. Additionally, examining clinical populations with impaired cognitive flexibility could provide insights into the functional significance of asymmetric switching mechanisms. Individuals with major depressive disorder characterized by rumination, for example, may exhibit altered subcortical coordination during internal-to-external transitions, while those with difficulty sustaining goal-directed behavior may show disrupted external domain maintenance. Characterizing these deficits could inform targeted interventions—such as subcortical neuromodulation or cognitive training—designed to facilitate adaptive transitions between internal and external processing in clinical contexts.

## Conclusion

Our study provides multimodal evidence for asymmetric neural mechanisms underlying transitions between external and internal cognitive domains. Using complementary EEG and fMRI approaches, we demonstrate that internal-to-external transitions uniquely engage subcortical networks and primary sensory cortex, while external-to-internal transitions show minimal differential activation. This asymmetry challenges prevailing a superordinate control mechanism and suggests that subcortical regions coordinate cortical network reconfiguration in a direction-dependent manner. The sustained neural activity observed during both preference switch and repeated conditions may reflect continuous engagement of self-referential processing networks, suggesting that internal domain processing resists habituation responses that characterize repeated external judgements. These findings provide insight into the distinct neural dynamics governing transition between internal and external cognitive domains.

## Methods

### Participants

#### EEG demographics

Forty-seven healthy adults were recruited for the EEG experiment (25 females; mean age = 27.5 ± 4.8 years). All participants had normal or corrected-to-normal vision and no self-reported psychiatric diagnoses, neurological diseases, or colour blindness. One participant was excluded due to technical issues during data collection, leaving 46 participants for behavioural analysis. An additional 6 participants were excluded from EEG statistical analyses due to insufficient trials remaining after artefact rejection preprocessing, resulting in a final number of 40 participants for EEG analyses. Detailed demographic information is in Table 1.

#### fMRI demographics

Sixty healthy adults were recruited for the fMRI experiment (32 females; mean age = 27.8 ± 4.9 years). All participants met standard fMRI inclusion criteria, and each confirmed verbally and via screening questionnaire during recruitment. Exclusion criteria included history of psychiatric or neurological disorders, previous head injury, current recreational drug use, metal implants, pregnancy, claustrophobia, or other MRI contraindications. All participants are right-handed, have normal or corrected-to-normal vision and have no self-reported colour blindness. For univariate analysis, three participants were rejected due to one or more runs’ motions frame-wise displacement above 5 mm (Jenkinson et al., 2002a), and two additional participants were rejected from any model related to Conditions due to high miss and incorrect trials in a run. Detailed demographic information is in Table 1.

#### Ethics and compensation

The experimental protocol was approved by the Taipei Medical University Institutional Review Board (IRB N202003129). Written informed consent was obtained from all participants prior to participation, and all participants received financial compensation for their time. For EEG, participants received 1000 NTD, equivalent to 33 USD. For fMRI, participants received 2400 NTD equivalent of 80 USD.

### General experimental task and procedure

We adapted a colour judgement paradigm from Hsu et al. (2021), which required participants to make judgements based on either external sensory information (colour similarity) or internal subjective preferences. While the original paradigm used a block design, we modified it to an interleaved design to investigate the neural dynamics of switching between external and internal domain processes.

On each trial, participants viewed a display containing three colour squares: one target square at the top and two choice squares positioned below and opposite each other (Figure 1A and 1D). Each square subtended 1.5° × 1.5° of visual angle. Participants performed one of two judgement types on each trial: colour similarity or colour preference judgements.

During colour similarity judgements (external domain), participants identified which of the two lower squares was most similar in hue to the target square. This task relies on objective perceptual comparison of external visual information. For colour preference judgements (internal domain), participants selected which of the two colour options they personally preferred, despite identical visual presentation and positioning. This task engages subjective, internally-driven decision-making processes.

Colour arrangements were designed in CIELAB colour space to control task difficulty and ensure comparable cognitive demands across conditions. For colour preference judgements, both colour options were positioned at equal 60° intervals from the target colour, ensuring no objective “correct” answer. For colour similarity judgements, the correct option was positioned 60° from the target colour, while the incorrect option was positioned 120° away, creating a discernible perceptual difference that could be objectively evaluated.

Trial conditions were defined based on consecutive transitions between judgement types. Repeat trials consisted of two consecutive trials requiring the same judgement type (similarity-similarity or preference-preference). Switch trials consisted of two consecutive trials requiring different judgement types (similarity-preference or preference-similarity). The number of switch and repeat trials was almost equally distributed across both EEG and fMRI experiments to enable balanced comparisons of neural activity between judgements and conditions.

#### EEG task design

The task flow EEG experiment is as follows: at the beginning of each trial, a fixation cross is presented for 1000 ms. Then, stimuli and judgement response cues were simultaneously presented for 2500 ms. Trials concluded with a blank screen serving as an intertrial interval for 1500 ms. The judgement type for each trial was indicated by the configuration of the fixation cross during the response phrase: for the preference judgement, the horizontal line was longer than the vertical line; for the similarity judgement, the vertical line was longer than the horizontal line (Figure 1A).

In the EEG experiment, the experiment comprised 500 trials organised into 10 blocks. The first 100 trials followed a block design with 50 trials of similarity and 50 trials of preference judgements to familiarise participants with both judgement types. The subsequent 400 trials employed an interleaved design incorporating both repeated and switch trials. To ensure balanced presentation of conditions while maintaining unpredictability, we implemented pseudo-randomisation that limited consecutive presentations of any single condition to a maximum of three trials.

#### fMRI task design

The task flow of the fMRI experiment is as follows: at first, the condition cue is presented on the screen for 500 ms. The condition cue is an explicit text cue (i.e., preference or similarity). Second, a 300-ms fixation cross. Third, the colour patch stimuli were presented for 500 ms. Third, a response cue is presented as a fixation cross at the centre while the colour stimuli are still presented for 1000 ms. During this time, participants are instructed to make a corresponding side response to the condition cue within this time window. Lastly, a blank screen appears, indicating that the intertrial interval and duration of this period are randomised between 2000 ms and 5000 ms (Figure 1D). To note, the task duration is prolonged compared to the EEG experiment, which gives ample time to help induce and detect HRF for fMRI.

The fMRI experiment is structured as follows: each participant performed a total of three blocks of the colour judgement task, and each block consisted of sixty-eight trials. The two types of judgements are counterbalanced, comprising thirty-four of each of the colour similarity and colour preference judgements. The order of judgement types within each block is pseudo-random across blocks. Across subtypes of Judgement and Condition trials, are not pseudo counterbalanced, which includes 18 preference switch trials, 16 preference repeated trials, 17 similarity switch trials and 16 similarity repeated trials.

#### Scan-day procedure

In the EEG experiment, stimuli were presented on a 54 cm monitor with participants positioned 46 cm from the screen using a chin rest to maintain a consistent viewing distance. In the fMRI experiment, stimuli were projected on a screen with 4:3 aspect ratio of 1024×768 resolution pixel layout inside the MRI machine. Both tasks were implemented using MATLAB R2019b/R2021a with Psychtoolbox (Brainard, 1997).

### Behavioural data

#### EEG behaviour data characteristic

The EEG experiment’s behavioural data included 46 participants with 400 trials from each participant, arranged in a single-trial format and categorised by Judgement (similarity and preference judgements) and Condition (switch and repeat trials). Reaction times (RTs) were de-centred values by subtracting participants’ mean RTs. Trials with missing or incorrect responses, as well as the first trial of each block, were excluded from analysis. Outliers were further excluded if RTs exceeded ±3 standard deviations from the participant’s mean, a procedure commonly applied to reduce bias in cognitive studies (Berger & Kiefer, 2021). After preprocessing, the dataset consisted of 16,518 observations.

Prior to model fitting, the univariate distributions of the overall RTs and the RTs of categorised conditions were visually inspected to check for potential violations of normality or linearity and to identify spurious effects on the dependent variable (Fife, 2020). In addition, potential confounding variables including block sequence, trial sequence were also inspected. During inspection, a systematic variability, suspected learning effect, in RTs across blocks were observed; therefore, block-to-block variance was considered added to the LME model.

#### fMRI behavioural data characteristic

The fMRI experiment’s behavioural data included 60 participants with 204 trials from each participant, and underwent the same preprocessing procedure as the EEG experiment. After rejection, the dataset comprised 10514 observations.

During the confounding inspection process, a clear learning effect was observed in trial-by-trial variability which characterised slower reaction in the beginning and progressively faster as more trials are performed. In addition, similar systematic variability across blocks were also observed. Thus trial-to-trial and block-to-block variance were considered added to the LME model.

### Behavioural analysis

#### EEG LME models

We conducted reaction time analysis with linear mixed effect (LME) model to account for variability due to trials sequencing and nested variance within participants. The LME modeling was conducted using the Pymer4 package in Python (Jolly, 2018).

Model selection process uses criteria mainly based on results of the likelihood ratio tests (LRTs) with lower score indicating better fitness of the model, with other fitness criteria such as Akaike Information Criterion (AIC), Bayesian Information Criterion (BIC) and log likelihood (LL) as supplement. With above criteria, we used the ground up approach by starting the LME model with each confounds and participants variables as random intercept, and gradually adding each interested categorised variable (Judgement and Condition) into the model to demonstrate each variable finestness as each variable progressively modeled. Details of the model comparison progression can be in Supplementary Table 1.1.

The final model is as followed:

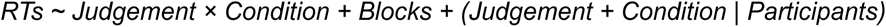

This model explained the variance of Reaction Times (RTs) with AIC is 237309.43 and BIC is 237448.25, and the log-likelihood (LL) score is −118636.71. LME were estimated via Restricted-Maximum-Likelihood-Estimation (Jolly, 2018), resulting fixed, random interaction effect, and confidence Intervals of random effect were estimated via bootstrapped method implemented with *boot* function in Pymer4 package (see Supplementary Table 1.2 for interaction model effect table). Post Hoc analysis was conducted on the final model using *emmeans* function to compute estimated marginal means (EMMs) for each interaction condition (see Figure 1D). In addition, pairwise comparisons with Tukey correction were conducted to estimate Judgement types differences in switch cost (Condition). Lastly, to cross-validate the robustness of the results, a type-III ANOVA was conducted, using the *anova* function, on the linear function of the predictor in the final model. The type III ANOVA uses Satterthwaite’s approximation of degrees of freedom for estimation (Kuznetsova et al., 2017). The ANOVA table is reported in Supplementary Table.1.2.

#### fMRI LME models

With the same model comparison criteria, the fMRI resulting best fitted RT model is as follows:

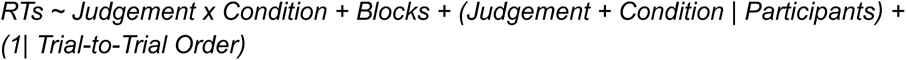

*This model explained the variance of RT with AIC is 134422.3 and BIC is 123516.7, and LL score is −67198.2* (Supplementary Table 2.1 for fMRI model comparison). The same model estimation method and robustness comparison were applied (see Supplementary Table 1.2 for the interaction Model details and Supplementary Table 1.3 for the ANOVA table). Since we found no interaction effect in the fMRI behavior model, further post hoc pairwise contrasts were not conducted. EMMs of each interaction condition was estimated for visualisation (see Figure 1H).

### EEG analysis

#### EEG acquisition and preprocessing

Continuous EEG data were recorded at a sampling rate of 1000 Hz using a standard 64 Ag/AgCl electrode cap (easycap; Brain Products GmbH). The electrode montage included 57 active scalp electrodes, with additional electrodes for measuring eye movements (4), mastoids (2), reference (1), and ground (1). Vertical EOG was recorded with electrodes above the left eye (VEOG-U) and below the right eye (VEOG-L), while horizontal EOG was recorded at the outer corners of both eyes (HEOG-L and HEOG-R). Mastoid electrodes (M1 and M2) were placed bilaterally, with online reference at FCz and ground at AFz. Electrode impedance was maintained below 10 kΩ throughout the experiment.

Offline preprocessing was performed using the MNE-Python (Gramfort et al., 2014). The continuous data underwent band-pass filtering (0.05-30 Hz) with Finite Impulse Response (FIR) filters (Widmann et al., 2015). Eye blink artefacts were corrected using Independent Component Analysis (ICA) with the fastICA algorithm (Hyvarinen, 1999). EEG event markers were aligned with behavioural data to identify missing trials, incorrect responses and the first trial of each block. Prior to epoching, signals were re-referenced to the average of the left and right mastoids.

Data were epoched from −100 ms to 1000 ms relative to stimulus onset, with the pre-stimulus interval (−100 to 0 ms) used for baseline correction. A two-stage artefact rejection procedure was implemented. First, amplitude-based rejection criteria of ±100 µV (EEG) and ±250 µV (EOG) were applied to remove residual artefacts. Participants with >50% rejected trials were excluded from further analyses. Second, outlier detection was performed on the baseline interval using z-scoring, with epochs exceeding ±5 standard deviations discarded (Luck, 2014; Wellcome Centre for Human Neuroimaging, 2021).

#### ERPs analysis

ERP analysis focused on the 0-1000 ms post-stimulus window. Prior to averaging, missing trials, incorrect responses, and the first trials of each block were excluded. To avoid bias from unequal trial numbers, epochs were equalised across conditions. Condition-specific waveforms were generated by averaging across trials within participants, followed by grand averaging across participants for visualisation.

We computed four key contrasts: (1) main effect of judgement type (preference vs. similarity), (2) main effect of condition type (switch vs. repeated), (3) within-judgement contrasts (switch vs. repeated for each judgement type separately). Statistical analysis employed a nonparametric spatiotemporal cluster-based permutation test (Groppe et al., 2011) applied across all electrodes and time points. Two-tailed tests were conducted with 1,024 permutations and a significance threshold of p < 0.05 (corrected). Spatial adjacency was defined using Delaunay triangulation, combined with temporal neighbours to form a joint channel × time adjacency matrix. A step-down correction procedure (α = 0.05) controlled family-wise error rate by sequentially testing clusters in order of significance (Maris & Oostenveld, 2007).

#### Time-resolved decoding and temporal generalisation analysis on EEG data

Time-resolved decoding followed established procedures (Carlson et al., 2019; Hubbard et al., 2019). Data were segmented into 50 ms windows with 1 ms sliding steps to capture stable neural patterns. For each participant, data were split into training (75%) and test (25%) sets for binary classification comparisons.

At each time point, data were z-score standardised using StandardScaler from the scikit-learn library (Pedregosa et al., 2011) to normalize data across trials to unit variance. The decoding pipeline employed supervised machine learning by applying Linear Discriminant Analysis (LDA) with 5-fold cross-validation. Classification performance was evaluated using Area Under the Curve (AUC), with scores averaged across folds to obtain grand AUC values.

We conducted complementary temporal generalisation analysis to evaluate neural representation stability across time (King & Dehaene, 2014; Li et al., 2024). This approach trains classifiers at each time point and tests them across all the other time points, generating a temporal generalisation matrix. Cross-validation was configured identically to temporal decoding, with performance evaluated using ROC-AUC. The resulting matrix captures both diagonal decoding accuracy (time-specific representations) and off-diagonal generalisation patterns reflecting neural code stability across time.

For group-level analysis, decoding results were pooled across participants and baseline-corrected by subtracting the chance level of 0.50 for binary classification (Fawcett, 2006). Statistical significance was assessed using cluster-based permutation one-sample t-tests with a cluster-forming threshold of p < 0.05. Condition labels were randomly permuted to generate null distributions compared against observed results. For temporal decoding, clusters were defined as sequential time points exceeding a threshold. For temporal generalisation, clusters represented continuous regions in the training × testing time matrix, assessing both diagonal and off-diagonal effects. Significant clusters indicated time periods (temporal decoding) or matrix regions (temporal generalisation) with reliable above-chance classification performance.

#### EEG analysis software

EEG preprocessing and ERP analysis were conducted in Python (v 3.11) with the MNE-Python package(v 1.8.0). EEG data were organised by implementing the Brain Imaging Data Structure (BIDS;v 0.15.0). The statistical analysis was performed with a combination of functions from the MNE-Python and SciPy (v 1.15.3). For decoding analysis, the sliding time window of the algorithm was performed by Sliding Estimator from MNE-Python in conjunction with the scikit-learn package (v 1.3.0) (Pedregosa et al., 2011). The decoding data was stored in the pickle format (v 4.0). The visualisation was done by using Matplotlib (v 3.10), seaborn(v 0.13.2) and some functions provided by MNE-Python.

#### fMRI analysis

##### MRI acquisition

Neuroimaging data were acquired on a 3 Tesla Siemens Magnetom Prisma scanner (syngo MR D13D software version) using a body coil for transmission and a 64-channel receive head coil conducted at the Imaging Centre for Integrated Body, Mind, and Culture Research at National Taiwan University. A magnetisation-prepared rapid acquisition-gradient echo (MPRAGE) sequence was used to obtain whole-brain T1-weighted structural images (208 sagittal slices, TR/TE/flip angle = 2000 ms/2.43 ms/ 9°, matrix = 288 × 288 mm^2^; voxel resolution = 0.88 × 0.88 × 0.88 mm^3^). Functional image obtained from an axial view aligned with the anterior-posterior commissure (AC-PC line) (37 axial slices, TR/TE/flip angle = 2000 ms/30 ms/ 90°, matrix = 64 × 64 mm^2^; voxel resolution = 3.5 × 3.5 × 3.5 mm^3^). Participants’ heads were secured with sponge pads and instructed to minimise their movements during scanning.

#### fMRI preprocessing

The initial preprocessing was conducted with fMRIprep (Esteban et al., 2019). The full preprocessed workflow generated by fMRIprep can be found in Supplementary 12. In short, the T1-weighted structural image was corrected for intensity non-uniformity, skull-stripped, and spatially normalised to the ICBM 152 Nonlinear Asymmetrical template version 2009c (MNI152Lin2009cAsym) space using the Advanced Normalisation Tools (ANTs) (Fonov et al., 2009; Tustison et al., 2021). Functional data were slice-timing corrected using 3dTshift from AFNI and motion-corrected with six degrees of freedom using MCFLIRT (Cox, 1996; Jenkinson et al., 2002b). The processed BOLD data were then co-registered to the corresponding T1-weighted structural image using boundary-based registration with six degrees of freedom (Greve & Fischl, 2009). Nuisance regressor derivatives such as FD, DVARS, and CompCor were computed. Lastly, the BOLD time series were spatially normalised to standard space using ANTs. In addition to fMRIprep preprocessing, spatial smoothing with a 5 mm FWHM Gaussian kernel with SUSUAN is applied (Jenkinson et al., 2012; Smith & Brady, 1997).

#### First-level process

First-level GLM-based analysis was carried out using FEAT (FMRI Expert Analysis Tool), part of FSL (FMRIB’s Software Library, https://www.fmrib.ox.ac.uk/fsl). Voxel time-series statistical analysis was carried out using FILM with local autocorrelation correction (Woolrich et al., 2001). Each explanatory variable (EV) was input as a custom time series with stimuli-onset and 1.5-second trial duration. The EVs’ waveform is then convoluted with the double gamma hemodynamic response function (HRF). Each model’s design matrix consists of nuisance regressors generated by fMRIprep, which includes anatomical global signals in cerebrospinal fluid, white matter, twenty estimated head-motion parameters, temporal CompCor, and cosine-basis regressors as a temporal high-pass filter (Behzadi et al., 2007; Satterthwaite et al., 2013).

A total of four first-level GLM-based analyses were modelled in the primary analysis to explore the depth of domain switching, which include the Judgement model (preference and similarity judgement), the Condition model (switch and repeated trials), and the judgement to Condition interaction model (similarity-repeated, similarity-switch, preference-repeated, and preference-switch conditions) and domain switching model (similarity-switch subtract similarity-repeated, preference-switch subtracted preference-repeated). The generated contrast of parameter estimate (COPE) from each model was merged and averaged across three runs within each participant.

#### RT duration modelling

We hypothesise that cognitive process differences between each condition may contribute to RT variances in our task design. At the same time, using fMRI analysis to identify regional differences in conditions has a known complex and confounded relationship between RTs and BOLD response (Yarkoni et al., 2009). To ensure the RTs’ variance is accounted for, we constructed three additional reaction time models to complement our primary analysis. To ensure our reaction time model accounted for Type-I errors as the number of EVs increases, we modelled each original EV with its trial response time (Grinband et al., 2008; Mumford et al., 2024). Trial response time is defined here by the boxcar durations of each trial, which are used to estimate the neural activity durations that match RTs durations (Grinband et al., 2008). Thus, our three RT first-level GLM-based models will include the original EVs, matching trial RT EVs, and nuisance regressors.

#### Group analysis

Group models were estimated using permutation-based methods from Permutation analysis Linear Model (PALM) (Winkler et al., 2014). The contrast difference between conditions was assessed using nonparametric one-tailed paired t-tests. Statistic maps were controlled over voxels and contrast-wise with family-wise error (FWER) correction, and a threshold below 0.05 based on 500 permutations (Alberton et al., 2020; Winkler et al., 2016).

#### Conjunction analysis

The group mean of each of the trial conditions from judgement to Condition interaction model was estimated as a non-parametric one-sample t-test. Statistic maps were controlled for a voxel-wise with FWER, and a threshold below 0.05 based on 500 permutations. A conjunction map was constructed by taking the overlap of the thresholded data (Nichols et al., 2005). A total of two construction maps are generated, including switch conjunction map overlapping switch trials of preference and similarity judgement (Supplementary 10 for Figure and Table) and repeated conjunction overlapping repeated trials type with preference and similarity judgement (Supplementary 11 for Figure and Table).

#### fMRI analysis software

Original DICOM images were converted to NIFTI format with dcm2niix (v1.0.20241211). Image preprocessing and statistical analyses were conducted with a combination of the following packages: fMRIprep (22.1.0), FSL (6.00), FreeSurfer (6.0.0), PALM (alpha115), Python Standard Library (3.11.6) and Python packages including, Nipype (1.8.6), Numpy (1.24.4), Pandas (2.2.3), Nipype (1.8.6) and Nibabel (5.3.2). Visualisation conducted with combination Python’s visualization packages including cmocean (v3.0.3), matplotlib (3.10.3), nilearn (0.11.1). ROI table generated automatically by AtlasReader (0.3.2) (Notter et al., 2019). All fMRI computations were conducted on a Linux distribution Pop!_OS 22.04 LTS with a desktop computer with CPU of AMD Ryzen 9 7950X.

## Acknowledgement

The authors thank all participants for their time and effort. This work was supported by funding from the National Science and Technology Council (NSTC113-2410-H-038-030; MOST111-2410-H-038-009-MY2) and the Taiwan Ministry of Education Higher Education Sprout Project to TYH. In addition, we would like to thank the Imaging Center for Integrated Body, Mind and Culture Research at NTU for fMRI data collection.

We followed CRediT statements to list each author’s contribution.

Risa Dwi Ratnasari: Investigation, Methodology, EEG Data Curation, Software, Visualisation, Writing - Original draft preparation

Paul Zen Cheng: Investigation, Methodology, fMRI Data Curation, Software, Visualisation, Writing - Original draft preparation

Tzu-Yu Hsu: Conceptualisation, Investigation, Methodology, Software, Writing - Reviewing and Editing, Supervision

## Data availability

To be disclosed upon publication.

## Code availability

To be disclosed upon publication.

