## supplementary for "Neural correlates of transitions between internal and external cognitive domains"

Supplementary materials

**Supplementary 1: EEG’s Behavior Linear Mixed Effect Model Information.**	**3**

Supplementary Table 1.1: Model Comparison	3

Supplementary Table 1.2: Interaction Model Effect Table	3

Supplementary Table 1.3: Interaction Model, ANOVA Table.	4

ANOVA type III	4

Supplementary Table 1.4: RT Distribution Quartiles by Judgement & Condition	5

**Supplementary 2: fMRI's Behavior Linear Mixed Effect Model Information.**	**6**

Supplementary Table 2.1: Model Comparison	6

Supplementary Table 2.2: Interaction Model Effect Table	6

Supplementary Table 2.3: Interaction model, ANOVA Table.	7

Supplementary Table 2.4: RT Distribution Quartiles by Judgement and Condition	7

**Supplementary 3: fMRI Main Models' Significant Contrast ROIs**	**8**

Supplementary Table 3.1: Similarity > Preference: Cluster Table.	8

Supplementary Table 3.2: Preference > Similarity: Cluster Table.	9

Supplementary Table 3.2: Condition Model: Cluster Table.	11

Supplementary Table 3.3: Interaction Model: Preference repeated vs Preference Switch Contrast, Cluster Table.	11

Supplementary Table 3.4: Interaction Model: Preference repeated vs Preference Switch Contrast, Cluster Table.	11

**Supplementary 4: fMRI Judgement Model with RTDUR as Covariates**	**13**

Supplementary Figure 4.1: Judgement RT Model: Preference vs Similarity, Pfwe < 0.05.	13

Supplementary Table 4.1: Judgement RT Model: Preference vs Similarity, Pfwe < 0.05, Cluster Table.	14

Supplementary Figure 4.2: Judgement RT Model: Preference vs Similarly, Unthreshold T map.	16

Supplementary Figure 4.3: Judgement RT Model: Preference vs Preference-RTDUR, Pfwe < 0.05.	17

Supplementary Table 4.3: Judgement RT Model: Preference vs Preference-RTDUR, Pfwe < 0.05, Cluster Table.	18

Supplementary Figure 4.4: Judgement RT Model: Preference vs Preference-RTDUR, Unthreshold T map.	22

Supplementary Figure 4.5: Judgement RT Model: Similarity vs Similarity-RTDUR, Pfwe < 0.05.	23

Supplementary Table 4.5: Judgement RT Model: Similarity vs Similarity-RTDUR, Pfwe < 0.05, Cluster Table.	24

Supplementary Figure 4.6: Judgement RT Model: Similarity vs Similarity-RTDUR, Unthreshold T map.	27

**Supplementary 5: fMRI Judgement Model with Cue-onset.**	**28**

Supplementary Figure 5.1: Judgement Cue-onset Model: Preference Cue-onset vs Similarity Cue-onset, Pfwe < 0.05.	28

Supplementary Table 5.1: Judgement Cue-onset Model :Preference Cue-Onset vs Similarity Cue-Onset, Pfwe < 0.05, Cluster Table.	29

Supplementary Figure 5.2: Judgement Cue-onset Model: Preference Cue-Onset vs Similarity Cue-Onset, Unthreshold T map.	32

**Supplementary 6: fMRI Condition Model with RTDUR as Covariates.**	**33**

Supplementary Figure 6.1: Condition RT Model: Switch Trial vs Repeated Trial, Pfdr <0.01.	33

Supplementary Table 6.1: Condition RT Model: Switch Trial vs Repeated Trial, Pfdr <0.01, Cluster Table.	34

Supplementary Figure 6.2: Condition RT Model: Switch Trial vs Repeated Trial, Unthreshold T map.	35

**Supplementary 7: fMRI Condition Model with Cue-onset.**	**36**

Supplementary Figure 7.1: Condition Cue-onset Model: Switch Trial's Cue-onset vs Repeated Trial's Cue-onset, Pfwe < 0.05.	36

Supplementary Table 7.1: Condition Cue-onset Model: Switch Trial's Cue-Onset vs Repeated trial's Cue-Onset, Pfwe < 0.05, Cluster Table.	37

Supplementary Figure 7.2: Condition Cue-onset Model: Switch Trial's Cue-Onset vs Repeated trial's Cue-Onset, Unthreshold T map.	38

**Supplementary 8: fMRI Judgement X Condition Model with RTDUR as Covariates.**	**39**

Supplementary Figure 8.1: Interaction RT Model: Preference Switch Cost Contrast, Pfdr < 0.01	39

Supplementary Table 8.1: Interaction RT Model: Preference Switch Cost Contrast, Pfdr < 0.01, Cluster Table.	40

Supplementary Figure 8.2: Interaction RT Model: Preference Switch Cost Contrast, Unthreshold T map.	41

Supplementary Figure 8.3: Interaction RT Model: Similarity Switch Cost Contrast, Pfwe < 0.05.	42

Supplementary Table 8.3: Interaction RT Model: Similarity Switch Cost Contrast, Pfwe < 0.05., Cluster Table.	43

Supplementary Figure 8.4: Interaction RT Model: Similarity Switch Cost Contrast, Unthreshold T map.	44

**Supplementary 9: fMRI Judgement X Condition Model with Cue onset.**	**45**

Supplementary Figure 9.1: Interaction Cue-onset Model: Preference Switch Cost Contrast with Cue-onset, Pfwe < 0.05.	45

Supplementary Table 9.1: Interaction Cue-onset Model: Preference Switch Cost Contrast with Cue-onset, Pfwe < 0.05, Cluster Table	46

Supplementary Figure 9.2: Interaction Cue-onset Model: Preference Switch Cost Contrast with Cue-onset, Unthreshold T map..	47

Supplementary Figure 9.3: Interaction Cue-onset Model: Similarity Switch Cost Contrast with Cue-onset, Pfwe < 0.05.	48

Supplementary Table 9.3: Interaction Cue-onset Model: Similarity Switch Cost Contrast with Cue-onset, Pfwe < 0.05, Cluster Table.	49

Supplementary Figure 9.4: Interaction Cue-onset Model: Similarity Switch Cost Contrast, Unthreshold T map..	50

**Supplementary 10: Conjunction Analysis for Switch Trials.**	**51**

Supplementary Figure 10.1:Interaction Model: Similarity Switch Condition, one sample two tailed t-test, Pfwe < 0.05	51

Supplementary Table 10.1: Interaction Model: Similarity Switch Condition, one sample two tailed t-test, Pfwe < 0.05, Cluster Table.	52

Supplementary Figure 10.2: Interaction Model: Preference Switch Condition, one sample two tailed t-test, Pfwe < 0.05.	57

Supplementary Table 10.2: Interaction Model: Preference Switch Condition, one sample two tailed t-test, Pfwe < 0.05, Cluster Table.	58

Supplementary Figure 10.3: Switch Conjunction Analysis, Pfwe < 0.05.	61

Supplementary Table 10.3: Switch Conjunction Analysis, Pfwe < 0.05, Cluster Table.	62

**Supplementary 11: Conjunction Analysis for Repeated Trials**	**66**

Supplementary Figure 11.1: Interaction Model: Similarity Repeated condition, one sample two tailed t-test, Pfwe < 0.05.	66

Supplementary Table 11.1: Interaction Model: Similarity Repeated condition, one sample two tailed t-test, Pfwe < 0.05, Cluster Table.	67

Supplementary Figure 11.2: Interaction Model: Preference Repeated condition, one sample two tailed t-test, Pfwe < 0.05.	71

Supplementary Table 11.2: Interaction Model: Preference Repeated condition, one sample two tailed t-test, Pfwe < 0.05, Cluster Table.	72

Supplementary Figure 11.3: Repeated Conjunction Analysis, Pfwe < 0.05.	75

Supplementary Table 11.3: Repeated Conjunction Analysis, Pfwe < 0.05, Cluster Table.	76

**Supplementary 12: fMRIprep output prompt**	**80**

### Supplementary 1: EEG’s Behavior Linear Mixed Effect Model Information.

#### Supplementary Table 1.1: Model Comparison

| Number of participants = 46 | | | | | | | | | | |
| --- | --- | --- | --- | --- | --- | --- | --- | --- | --- | --- |
| Number of participants = 46  Number of observation = 16518 |  |  |  | Random Effects |  | Model fits |  |  | LRT Test against nested | |
| Model specification | Model name | Nest/simpler model | Fixed effects added | participants | block | AIC | BIC | LL | df | Chisq |
| RE only | Null | N/A |  | intercept | intercept | 239254.8 | 239285.7 | −119623.4 |  | n/a |
| FE 1 | Judgement Main effect | Null | Judgement | .. | intercept | 238685.1 | 238723.7 | −119337.6 | 1 | <.001*** |
| FE 1 +RE | Judgement Main effect across participants | Judgement Main effect | Judgement | slope | intercept | 237408.9 | 237462.8 | −118697.4 | 2 | <.001*** |
| FE 2 | Condition main effect | Null | condition | intercept | intercept | 239249.2 | 239287.8 | −119619.6 | 1 | n/a |
| FE 2 + RE | Condition main effect across participants | Condition main effect | condition | slope | intercept | 239246.3 | 239300.2 | −119616.1 | 2 | <0.05 * |
| FE 1 & 2 + RE | Judgement & Condition main effects | N/A | Judgement + condition | slope | intercept | 237395.7 | 237480.5 | −118686.8 |  |  |
| FE 1 X 2 + RE | Judgement & Condition interaction | Judgement & Condition main effects | Judgement + condition | slope | intercept | 237371.1 | 237463.6 | −118673.5 | 1 | <.001*** |
| FE 1 X 2 + FE 3 + RE | Judgement & Condition interaction with Block as fixed effect | Judgement & Condition interaction | Judgement + condition + Block | slope | n/a | 237309.4 | 237448.2 | −118636.7 | 1 | <.001*** |
| FE = Fixed effect; RE = Random effect, X = interaction | | | | | | | | | | |
| Model comparison likelihood ratio test based on Akaike Information criterion (AIC), Bayesian information criterion (BIC), and log likelihood (LL). The second model with participant (R1) random slope has significantly improved model performance. The df is the degree of freedom that shows the number of parameters included in the parameter.; Chisq is the chi-square statistic calculated from the difference in log-likelihoods. ; The likelihood ratio test (LRT) compares the simpler model against the more complex nested model; a significant result indicates that the improvement is meaningful for model performance. | | | | | | | | | | |

#### Supplementary Table 1.2: Interaction Model Effect Table

| Fixed effect | | | | | | | | | | | | | | | | | | | | | |
| --- | --- | --- | --- | --- | --- | --- | --- | --- | --- | --- | --- | --- | --- | --- | --- | --- | --- | --- | --- | --- | --- |
|  | | | 𝞫 | | | SE | | | 95% CI | | | t | | | df | | | p | | |  |
| (Intercept) | | | -23.049 | | | 39.881 | | | -103.243, 57.144 | | | -0.577 | | | 47.836 | | | 0.566 | | |  |
| Judgement | | | 101.505 | | | 29.468 | | | 42.244, 160.765 | | | 3.444 | | | 47.666 | | | <.01 | | | *** |
| Condition | | | -29.164 | | | 8.122 | | | -45.827, -13.041 | | | -3.591 | | | 95.246 | | | <.001 | | | *** |
| Block1-Block1 | | | 12.846 | | | 9.865 | | | -6.489, 32.182 | | | 1.302 | | | 16,389.113 | | | 0.1928 | | | m/s |
| Block2-Block1 | | | 20.041 | | | 9.840 | | | 0.754, 39.329 | | | 2.037 | | | 16,389.385 | | | <0.05 | | | * |
| Block3-Block1 | | | 13.730 | | | 9.851 | | | −5.579, 33.038 | | | 1.394 | | | 16,394.345 | | | 0.1634 | | |  |
| Block4-Block1 | | | −5.806 | | | 9.831 | | | −25.076, 13.465 | | | 0.591 | | | 16,389.480 | | | 0.554 | | |  |
| Block5-Block1 | | | -64.685 | | | 9.873 | | | −84.037, −45.333 | | | −6.552 | | | 16,389.088 | | | <.001 | | |  |
| Block6-Block1 | | | −74.535 | | | 9.831 | | | −93.805, −55.265 | | | −7.582 | | | 16,385.314 | | | <.001 | | |  |
| Block7-Block1 | | | −63.625 | | | 9.840 | | | −82.913, −44.338 | | | −6.466 | | | 16,394.518 | | | <.001 | | |  |
| Judgement X Condition | | | 44.286 | | | 9.852 | | | 24.975, 63.596 | | | 4.495 | | | 16,405.567 | | | <.001 | | | *** |
| Random effect | | | | | | | | | | | | | | | | | | | | | |
|  | | | Variance | | | S.D. | | | S.D. est. 95% Boot CI | | | | | | Correlation | | | | | | |
| Participant (Intercept) | | | 70162.8 | | | 264.88 | | | 209.18, 313.5821 | | | | | |  | | | | | | |
| Judgement \| Participant (slope) | | | 37681.8 | | | 194.12 | | | 155.264, 233.055 | | | | | | -0.54 | | | | | | |
| Condition \| Participant (slope) | | | 984.2 | | | 31.37 | | | 13.593, 43.964 | | | | | | 0.17 -0.24 | | | | | | |
| Result summary of primary linear-mixed-effect model predicting the task response times. The linear mixed-effects model fitted using restricted maximum likelihood (REML), including fixed effects for Judgement type, condition, and their interaction. Reported values include estimates (β), standard errors (SE), 95% confidence intervals (CI), degrees of freedom (df), and significance levels. The model included a random slope for participants. The lower table reports the random effect with parameter score of Variance, S.D (standard deviation), Boot CI (Bootstrap confidence interval) and correlation of the effect of participant slope to the model effect | | | | | | | | | | | | | | | | | | | | | |
| Model fit | | | | | | | | | | | | | | | | | | | | | |
| R^2^ | | | | | | | | Marginal | | | | | | | | Conditional | | | | | |
|  |  |  |  |  |  |  |  | 0.408 | | | | | | | | 0.031 | | | | | |
| Key: p-values of fixed effects, as well as Confidence intervals calculated using Satterthwaite's approximations.  Model equation: RT ~ Judgement X Condition + Block + (1 \| Trial Order) + (Judgement X Condition \| participant)  Note: 𝞫 = Estimated beta value, SE =estimated standard error of mean, CI= confidence interval. Significant's level is indicated by * with following index: *p < 0.05, **p < 0.01, ***p< 0.005,. | | | | | | | | | | | | | | | | | | | | | |

#### Supplementary Table 1.3: Interaction Model, ANOVA Table.

| ANOVA type III | | | | | | |
| --- | --- | --- | --- | --- | --- | --- |
| Model Term | df1* | df2* | F ratio | chi-square | *p* value | sig. |
| Judgement | 1 | inf | 18.125 | 18.125 | <.001 | *** |
| Condition | 1 | inf | 1.080 | 1.080 | 0.298 |  |
| Block | 7 | inf | 33.581 | 235.067 | <.001 | *** |
| Judgement x condition | 1 | inf | 20.207 | 20.207 | <.001 | *** |
| Significant Codes: *** : *p* < 0.001, ** : *p* < 0.01, * : *p* < 0.05. | | | | | | |
| *Satterthwaite's approximation was used to calculate degrees of freedom. | | | | | | |

#### Supplementary Table 1.4: RT Distribution Quartiles by Judgement & Condition

| **Judgement Type** | **Condition** | **Q1 (ms)** | **Median (ms)** | **Q3 (ms)** | **IQR (ms)** |
| --- | --- | --- | --- | --- | --- |
| Preference | Repeated | 885.75 | 1171.39 | 1486.52 | 600.77 |
| Preference | Switch | 919.47 | 1197.12 | 1495.28 | 575.81 |
| Similarity | Repeated | 1056.97 | 1321.56 | 1608.99 | 552.02 |
| Similarity | Switch | 1049.04 | 1300.99 | 1590.94 | 541.90 |

### Supplementary 2: fMRI's Behavior Linear Mixed Effect Model Information.

#### Supplementary Table 2.1: Model Comparison

| Number of observation = 10828 |  |  |  | Random Effects |  |  | Model fits |  |  | LRT Test against nested | |
| --- | --- | --- | --- | --- | --- | --- | --- | --- | --- | --- | --- |
| Model specification | Model name | Nest/simpler model | Fixed effects added | participants | trial-to-trial | Blocks | AIC | BIC | LL | df | Chisq |
| RE only | Null | N/A |  | intercept | intercept | intercept | 138698.42 | 138734.87 | -69344.21 |  | n/a |
| FE 1 | Judgement Main effect | Null | Judgement | .. | .. | .. | 138681.91 | 138725.64 | -69334.95 |  | 13.76*** |
| FE 1 + RE | Judgement Main effect across participants | Judgement Main effect | Judgement | slope | .. | .. | 138611.18 | 138669.5 | -69297.59 | 2 | 74.02*** |
| FE 2 | Condition main effect | Null | condition | intercept | intercept | intercept | 138683.2 | 138726.94 | -69335.6 |  | 12.45*** |
| FE 2 + RE | Condition main effect across participants | Condition main effect | condition | slope | intercept | intercept | 138674.14 | 138732.46 | -69329.07 | 2 | 12.87** |
| FE 1 & 2 + RE | Judgement & Condition main effects | N/A | Judgement + condition | slope | intercept | intercept | 138581.27 | 138668.75 | -69278.63 |  | n/a |
| FE 1 X 2 + RE | Judgement & Condition interaction | Judgement & Condition main effects | Judgement + condition | slope | intercept | intercept | 138577.01 | 138671.78 | -69275.5 | 1 | 0.32 |
| FE 1 X 2 + FE 3 + RE | Judgement & Condition interaction with Block as fixed effect | Judgement & Condition interaction | Judgement + condition + Block | slope | intercept | N/A | 134422.3 | 134516.7 | −67198.2 |  | 13.05*** |
| FE = Fixed effect; RE = Random effect; X = interaction | | | | | | | | | | | |

#### Supplementary Table 2.2: Interaction Model Effect Table

| Fixed Effects | | | | | | | |
| --- | --- | --- | --- | --- | --- | --- | --- |
|  | 𝞫 | SE | 95% CI | t | df | p |  |
| Intercept | 17.603 | 10.325 | -2.996,38.202 | 1.705 | 68.938 | 0.093 |  |
| Judgement | 13.813 | 6.799 | 0.301, 27.325 | 2.032 | 87.949 | 0.045 | * |
| Condition | 11.952 | 5.352 | 1.257, 22.646 | 2.233 | 63.435 | 0.029 | * |
| Block2 - Block1 | -41.142 | 3.343 | -47.696,-34.589 | -12.307 | 10291.307 | 0 | *** |
| Block3 - Block1 | -52.857 | 3.35 | -59.423,-46.29 | -15.779 | 10293.875 | 0 | *** |
| Judgement X Condition | 6.278 | 7.271 | -8.249, 20.804 | 0.863 | 63.923 | 0.391 |  |
| Random Effect | | | | | | | |
|  | | | | Variance | S.D. | S.D. est. 95 % Boot CI | Correlation |
| Trial Order (Intercept) | | | | 94.54 | 9.723 | 4.89, 13.181 |  |
| Participant (Intercept) | | | | 5290.16 | 72.733 | 57.95, 87.072 |  |
| Judgement \| Participant (slope) | | | | 1216.59 | 34.880 | 25.872, 43.244 | -0.27 |
| Condition \| Participant (slope) | | | | 226.98 | 15.066 | 7.237, 22.384 | 0.54 |
| Model fit | | | | | | | |
| R^2^ | | | | Marginal | | Conditional | |
|  | | | | 0.025 | | 0.249 | |
| Key: p-values of fixed effects calculated using Satterthwaite's approximations. Confidence intervals have been calculated using Satterthwaite's method.  Model equation: RT ~ Judgement X Condition + Block + (1 \| Trial Order) + (Judgement X Condition \| participant)  Note: 𝞫 = Estimated beta value, SE =estimated standard error of mean, CI= confidence interval. Significant's level is indicated by * with following index: *p < 0.05, **p < 0.01, ***p< 0.005,. | | | | | | | |

#### Supplementary Table 2.3: Interaction model, ANOVA Table.

| ANOVA type III | | | | | | |
| --- | --- | --- | --- | --- | --- | --- |
| Model Term | df1* | df2* | F ratio | chi-square | *p* value | sig. |
| Judgement | 1 | inf | 8.398 | 8.398 | 0.004 | ** |
| Condition | 1 | inf | 13.293 | 13.293 | <.001 | *** |
| Block | 1 | inf | 137.44 | 274.88 | <.001 | *** |
| Judgement x condition | 1 | inf | 0.745 | 0.745 | 0.388 |  |
| Significant Codes: *** : *p* < 0.001, ** : *p* < 0.01, * : *p* < 0.05. | | | | | | |
| *Satterthwaite's approximation was used to calculate degrees of freedom. | | | | | | |

#### Supplementary Table 2.4: RT Distribution Quartiles by Judgement and Condition

| **Judgement Type** | **Condition** | **Q1 (ms)** | **Median (ms)** | **Q3 (ms)** | **IQR (ms)** |
| --- | --- | --- | --- | --- | --- |
| Preference | Repeated | 352.54 | 457.70 | 576.03 | 223.49 |
| Preference | Switch | 350.12 | 449.20 | 533.30 | 183.18 |
| Similarity | Repeated | 377.46 | 480.90 | 596.40 | 218.94 |
| Similarity | Switch | 363.37 | 465.00 | 573.11 | 209.74 |

### Supplementary 3: fMRI Main Models' Significant Contrast ROIs

#### Supplementary Table 3.1: Similarity > Preference: Cluster Table.

Task: Similarity > Preference

| Anatomical label | Hemisphere | x | y | z | cluster mean T value | mm3 |
| --- | --- | --- | --- | --- | --- | --- |
| 52.11% Caudate  44.57% no label | R  n/a | 9.5 | 9.5 | 5.5 | 6.85 | 3608 |
| 93.57% Occipital Mid  5.00% no label | L  n/a | -30.5 | -86.5 | 7.5 | 6.34 | 3360 |
| 52.74% Parietal Sup  27.40% Parietal Inf  13.70% Occipital Sup  6.16% no label | R  R  R  n/a | 27.5 | -60.5 | 61.5 | 6.14 | 2336 |
| 69.59% Caudate  24.42% no label | L  n/a | -8.5 | 5.5 | 3.5 | 6.93 | 1736 |
| 57.67% Vermis 9  21.47% Vermis 8  6.75% Vermis 10  5.52% Cerebelum 9 | n/a  n/a  n/a  L | -0.5 | -58.5 | -38.5 | 6.45 | 1304 |
| 75.97% Calcarine  20.13% Calcarine | R  L | 17.5 | -70.5 | 13.5 | 6.02 | 1232 |
| 52.29% Insula  37.91% no label  5.88% Frontal Inf Tri | R  n/a  R | 27.5 | 25.5 | 1.5 | 6.23 | 1224 |
| 90.41% Occipital Mid  5.48% Occipital Sup | R  R | 31.5 | -80.5 | 37.5 | 6.17 | 1168 |
| 60.44% Frontal Sup 2  17.58% Precentral  14.29% Frontal Mid 2  7.69% no label | R  R  R  n/a | 27.5 | -4.5 | 53.5 | 6.23 | 728 |
| 44.68% Precentral  29.79% Frontal Sup 2  25.53% Frontal Mid 2 | L  L  L | -28.5 | -2.5 | 55.5 | 6 | 376 |
| 100.00% Calcarine | L | -4.5 | -70.5 | 11.5 | 6.23 | 352 |
| 87.18% Frontal Inf Oper  10.26% Frontal Inf Tri | R  R | 47.5 | 11.5 | 25.5 | 5.98 | 312 |
| 100.00% no label | n/a | 3.5 | -30.5 | -0.5 | 7.02 | 272 |
| 100.00% Temporal Inf | R | 45.5 | -58.5 | -10.5 | 6.34 | 240 |
| 91.67% Occipital Mid  8.33% no label | R  n/a | 33.5 | -84.5 | 5.5 | 6.01 | 192 |
| 63.64% Parietal Sup  36.36% Parietal Inf | R  R | 45.5 | -44.5 | 55.5 | 6.07 | 176 |
| 80.00% Cerebelum 7b  20.00% Cerebelum Crus2 | L  L | -8.5 | -72.5 | -40.5 | 6.1 | 160 |
| 83.33% Precentral  16.67% Frontal Inf Oper | R  R | 53.5 | 11.5 | 37.5 | 5.97 | 144 |
| 61.11% Cerebelum 7b  38.89% Cerebelum 8 | R  R | 9.5 | -74.5 | -42.5 | 6.18 | 144 |
| 100.00% Parietal Sup | L | -20.5 | -70.5 | 45.5 | 5.86 | 136 |
| 100.00% Calcarine | L | -16.5 | -74.5 | 7.5 | 5.92 | 104 |
| 100.00% Occipital Mid | R | 45.5 | -70.5 | 23.5 | 6.21 | 96 |
| 60.00% Fusiform  40.00% Cerebelum 6 | R  R | 33.5 | -56.5 | -18.5 | 5.94 | 80 |
| Cluster defined is 10 voxels | | | | | | |

#### Supplementary Table 3.2: Preference > Similarity: Cluster Table.

**Task: Preference > Similarity**

| Anatomical label | Hemisphere | x | y | z | cluster mean T value | mm3 |
| --- | --- | --- | --- | --- | --- | --- |
| 43.69% Frontal Sup 2  30.44% Frontal Sup Medial  15.44% Frontal Mid 2 | L  L  L | -14.5 | 39.5 | 53.5 | 7.09 | 23840 |
| 53.80% Angular  32.47% no label  9.94% Parietal Inf | L  n/a  L | -50.5 | -62.5 | 35.5 | 7.47 | 9904 |
| 66.71% Frontal Sup 2  29.83% Frontal Sup Medial | R  R | 17.5 | 35.5 | 53.5 | 6.53 | 6920 |
| 38.54% Frontal Inf Tri  26.32% Frontal Inf Orb 2  11.75% OFCpost  11.16% no label  6.11% OFClat | L  L  L  n/a  L | -50.5 | 25.5 | -6.5 | 6.9 | 6808 |
| 60.61% Cerebelum Crus1  38.81% Cerebelum Crus2 | R  R | 37.5 | -78.5 | -34.5 | 6.55 | 5504 |
| 78.17% Angular  16.50% Parietal Inf  5.08% no label | R  R  n/a | 51.5 | -60.5 | 35.5 | 6.83 | 3152 |
| 39.58% Cingulate Post  30.08% Cingulate Mid  22.16% Precuneus  6.07% no label | L  L  L  n/a | -4.5 | -50.5 | 33.5 | 6.49 | 3032 |
| 92.88% Temporal Mid  7.12% no label | L  n/a | -66.5 | -22.5 | -10.5 | 6.37 | 2920 |
| 44.51% Temporal Inf  21.10% no label  17.92% Temporal Mid  16.47% Temporal Pole Mid | L  n/a  L  L | -50.5 | -2.5 | -34.5 | 6.58 | 2768 |
| 49.08% Temporal Inf  45.02% Temporal Mid  5.17% Temporal Pole Mid | R  R  R | 49.5 | -6.5 | -30.5 | 6.39 | 2168 |
| 80.18% Cerebelum Crus2  16.13% Cerebelum Crus1 | L  L | -30.5 | -82.5 | -40.5 | 6.51 | 1736 |
| 54.10% Frontal Mid 2  27.87% no label  18.03% OFClat | L  n/a  L | -42.5 | 49.5 | -14.5 | 6.16 | 488 |
| 100.00% Cingulate Mid | L | -2.5 | -24.5 | 37.5 | 6.21 | 384 |
| 82.61% Cerebelum Crus2  17.39% Cerebelum 7b | R  R | 45.5 | -54.5 | -44.5 | 6.38 | 368 |
| 92.31% Precuneus  7.69% Precuneus | L  R | -0.5 | -64.5 | 37.5 | 5.86 | 208 |
| 68.18% Temporal Pole Mid  31.82% Temporal Pole Sup | R  R | 33.5 | 21.5 | -34.5 | 6.17 | 176 |
| 66.67% Frontal Inf Orb 2  33.33% OFClat | R  R | 47.5 | 31.5 | -14.5 | 5.98 | 168 |
| 54.55% Cingulate Ant  45.45% Frontal Med Orb | L  L | -6.5 | 39.5 | -6.5 | 6.11 | 88 |
| 100.00% Hippocampus | R | 29.5 | -10.5 | -20.5 | 5.89 | 88 |
| 90.00% Frontal Mid 2  10.00% no label | R  n/a | 39.5 | 29.5 | 43.5 | 5.87 | 80 |
| Cluster defined is 10 voxels | | | | | | |

#### Supplementary Table 3.2: Condition Model: Cluster Table.

| Task | Anatomical label | Hemisphere | x | y | z | cluster mean T value | mm3 |
| --- | --- | --- | --- | --- | --- | --- | --- |
| Switch > repeated | 89.47% Precuneus  5.26% Occipital Sup  5.26% Cuneus | L  L  L | -8.5 | -68.5 | 37.5 | 6.05 | 304 |
|  | 61.54% Precentral  38.46% Postcentral | L  L | -38.5 | -20.5 | 51.5 | 6.06 | 104 |
| Cluster defined is 10 voxels | | | | | | | |

#### Supplementary Table 3.3: Interaction Model: Preference repeated vs Preference Switch Contrast, Cluster Table.

| Task | Anatomical label | Hemisphere | x | y | z | cluster mean T value | mm3 |
| --- | --- | --- | --- | --- | --- | --- | --- |
| Preference repeated > Preference switch | 97.47% Cerebelum Crus1 | R | 29.5 | -84.5 | -28.5 | 6.36 | 632 |
|  | 64.29% no label  28.57% Thalamus  7.14% Thalamus | n/a  R  L | -2.5 | -8.5 | 11.5 | 5.88 | 224 |
|  | 95.24% Caudate | R | 7.5 | 11.5 | 3.5 | 6.28 | 168 |
| Cluster defined is 10 voxels | | | | | | | |

#### Supplementary Table 3.4: Interaction Model: Preference repeated vs Preference Switch Contrast, Cluster Table.

| Task | Anatomical label | Hemisphere | x | y | z | cluster mean T value | mm3 |
| --- | --- | --- | --- | --- | --- | --- | --- |
| Similarity switch > Similarity repeated | 49.17% Postcentral  42.56% Precentral  8.26% no label | L  L  n/a | -38.5 | -22.5 | 51.5 | 6.77 | 3872 |
|  | 84.77% Calcarine  12.69% Lingual | L  L | -16.5 | -70.5 | 7.5 | 6.26 | 1576 |
|  | 100.00% Calcarine | R | 11.5 | -66.5 | 13.5 | 6.24 | 888 |
|  | 78.79% Caudate  14.14% no label | R  n/a | 11.5 | 15.5 | -8.5 | 6.29 | 792 |
|  | 85.94% Supp Motor Area  14.06% Cingulate Mid | L  L | -2.5 | -8.5 | 53.5 | 6.13 | 512 |
|  | 85.29% Calcarine  11.76% Lingual | R  R | 21.5 | -52.5 | 3.5 | 5.98 | 272 |
|  | 62.50% Putamen  16.67% Pallidum  16.67% no label | L  L  n/a | -16.5 | 9.5 | -0.5 | 6.07 | 192 |
|  | 56.25% Cingulate Mid  31.25% Cingulate Mid  12.50% Frontal Sup Medial | R  L  L | 1.5 | 19.5 | 37.5 | 5.84 | 128 |
|  | 50.00% Cingulate Mid  50.00% Supp Motor Area | L  L | -10.5 | 5.5 | 43.5 | 6.11 | 112 |
|  | 53.85% no label  30.77% Putamen  15.38% Olfactory | n/a  L  L | -22.5 | 7.5 | -12.5 | 6.08 | 104 |
| Cluster defined is 10 voxels | | | | | | | |

### Supplementary 4: fMRI Judgement Model with RTDUR as Covariates

#### Supplementary Figure 4.1: Judgement RT Model: Preference vs Similarity, *P_fwe_ < 0.05.*

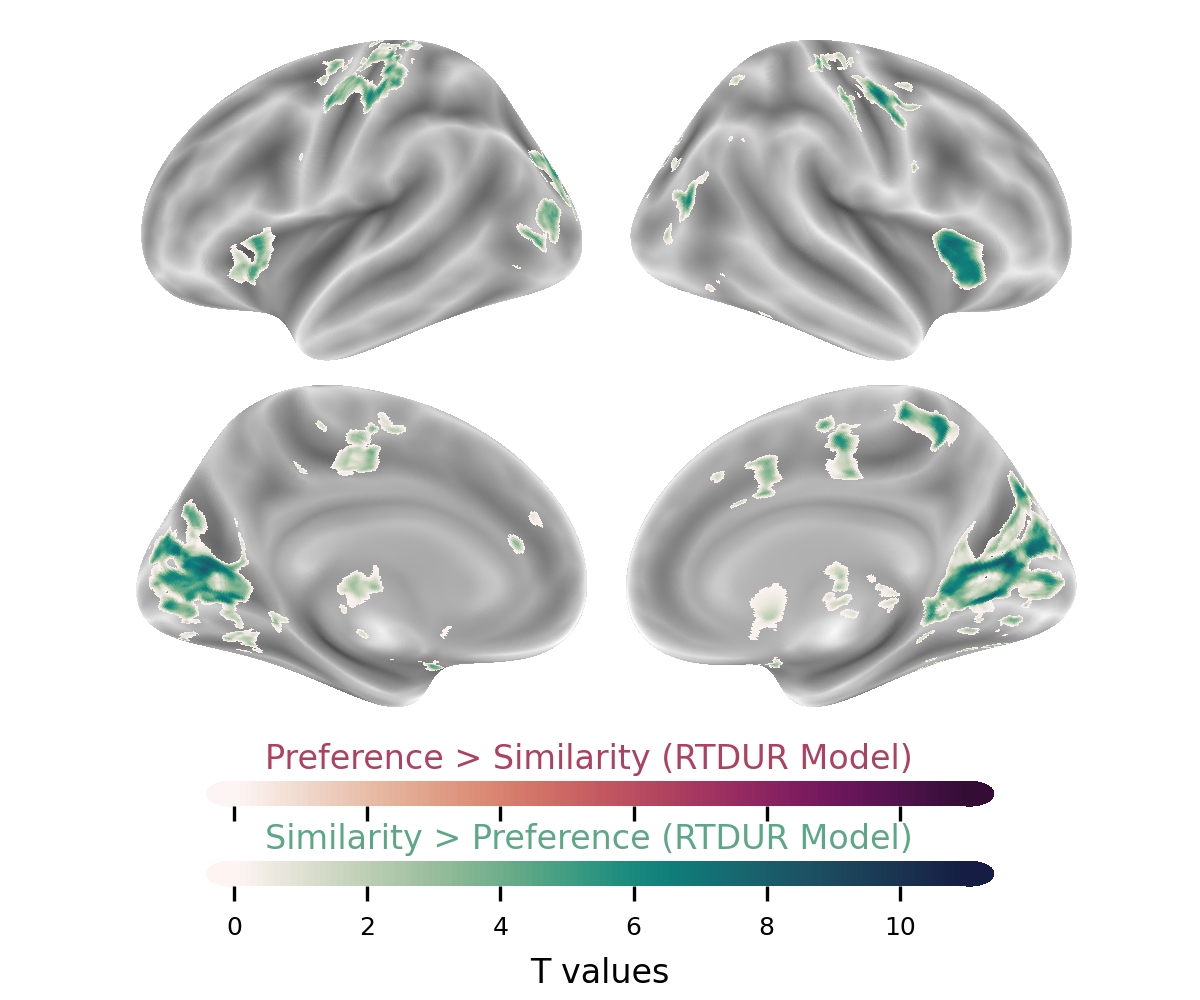

#### Supplementary Table 4.1: Judgement RT Model: Preference vs Similarity, *P_fwe_ < 0.05*, Cluster Table.

| Task | Anatomical label | Hemisphere | x | y | z | cluster mean T value | mm3 |
| --- | --- | --- | --- | --- | --- | --- | --- |
| Similarity > Preference | 33.98% Calcarine  30.82% Calcarine  9.54% Cuneus  9.03% Cuneus  7.80% Lingual  7.48% Lingual | L  R  L  R  R  L | 11.5 | -66.5 | 7.5 | 6.6 | 12408 |
|  | 49.23% Caudate  27.86% Putamen  19.27% no label | R  R  n/a | 13.5 | 9.5 | 7.5 | 7.07 | 7264 |
|  | 51.10% Caudate  32.97% Putamen  12.09% no label | L  L  n/a | -14.5 | 15.5 | 9.5 | 6.92 | 5824 |
|  | 60.38% Postcentral  34.23% Precentral  5.39% no label | L  L  n/a | -48.5 | -14.5 | 51.5 | 6.36 | 2968 |
|  | 85.12% Precentral  12.81% Frontal Mid 2 | R  R | 43.5 | -12.5 | 51.5 | 6.44 | 1936 |
|  | 53.85% Occipital Mid  44.34% Occipital Sup | L  L | -26.5 | -86.5 | 25.5 | 6.11 | 1768 |
|  | 72.60% Insula  16.44% no label  7.31% Frontal Inf Tri | R  n/a  R | 33.5 | 19.5 | 9.5 | 6.66 | 1752 |
|  | 68.18% Cuneus  22.73% Occipital Sup | R  R | 19.5 | -76.5 | 35.5 | 6.13 | 880 |
|  | 81.00% Precuneus  17.00% Paracentral Lobule | R  R | 7.5 | -46.5 | 55.5 | 6.44 | 800 |
|  | 95.12% Insula | L | -32.5 | 21.5 | -0.5 | 6 | 656 |
|  | 100.00% Thalamus | L | -6.5 | -22.5 | 7.5 | 6.14 | 600 |
|  | 100.00% Thalamus | R | 13.5 | -20.5 | 9.5 | 6.17 | 552 |
|  | 100.00% Occipital Mid | L | -38.5 | -80.5 | 9.5 | 6.04 | 480 |
|  | 67.31% Temporal Mid  32.69% Occipital Mid | R  R | 45.5 | -72.5 | 15.5 | 6.2 | 416 |
|  | 81.25% Cingulate Mid  18.75% Supp Motor Area | R  R | 9.5 | -18.5 | 45.5 | 6.11 | 384 |
|  | 84.09% Occipital Mid  15.91% Temporal Mid | L  L | -48.5 | -78.5 | 3.5 | 6.08 | 352 |
|  | 100.00% Lingual | R | 13.5 | -78.5 | -4.5 | 6.05 | 312 |
|  | 79.49% Occipital Sup  20.51% Cuneus | L  L | -14.5 | -78.5 | 27.5 | 6.27 | 312 |
|  | 79.31% Postcentral  20.69% Paracentral Lobule | L  L | -20.5 | -28.5 | 71.5 | 6.15 | 232 |
|  | 74.07% Postcentral  25.93% Precentral | R  R | 31.5 | -28.5 | 63.5 | 6.04 | 216 |
|  | 100.00% Occipital Sup | R | 19.5 | -88.5 | 29.5 | 5.95 | 208 |
|  | 96.15% Precentral | L | -26.5 | -12.5 | 57.5 | 6.13 | 208 |
|  | 91.67% Fusiform  8.33% Occipital Inf | R  R | 35.5 | -72.5 | -12.5 | 6.01 | 192 |
|  | 86.36% Cingulate Mid  13.64% Cingulate Mid | R  L | 11.5 | 7.5 | 41.5 | 5.91 | 176 |
|  | 65.00% Supp Motor Area  35.00% Supp Motor Area | R  L | 1.5 | -10.5 | 55.5 | 6.06 | 160 |
|  | 100.00% Fusiform | R | 37.5 | -46.5 | -20.5 | 6.12 | 152 |
|  | 50.00% Frontal Inf Oper  50.00% Precentral | R  R | 55.5 | 11.5 | 37.5 | 5.99 | 144 |
|  | 62.50% Occipital Mid  37.50% Occipital Sup | L  L | -20.5 | -96.5 | 15.5 | 5.96 | 128 |
|  | 93.75% Cingulate Mid  6.25% no label | L  n/a | -10.5 | -16.5 | 43.5 | 5.92 | 128 |
|  | 92.31% Postcentral  7.69% no label | R  n/a | 25.5 | -26.5 | 57.5 | 5.86 | 104 |
|  | 58.33% Cingulate Mid  33.33% Supp Motor Area  8.33% Paracentral Lobule | L  L  L | -8.5 | -22.5 | 49.5 | 6.03 | 96 |
|  | 100.00% Lingual | L | -24.5 | -64.5 | -8.5 | 6.13 | 88 |
|  | 100.00% Lingual | L | -16.5 | -44.5 | -8.5 | 6.13 | 88 |
|  | 100.00% Frontal Sup 2 | L | -26.5 | -6.5 | 67.5 | 6.13 | 88 |
|  | 100.00% Parietal Sup | R | 27.5 | -62.5 | 57.5 | 5.91 | 80 |
|  | 70.00% Supp Motor Area  20.00% Supp Motor Area  10.00% Paracentral Lobule | L  R  L | -0.5 | -20.5 | 51.5 | 5.93 | 80 |
| Cluster defined is 10 voxels | | | | | | | |

#### Supplementary Figure 4.2: Judgement RT Model: Preference vs Similarly, Unthreshold T map.

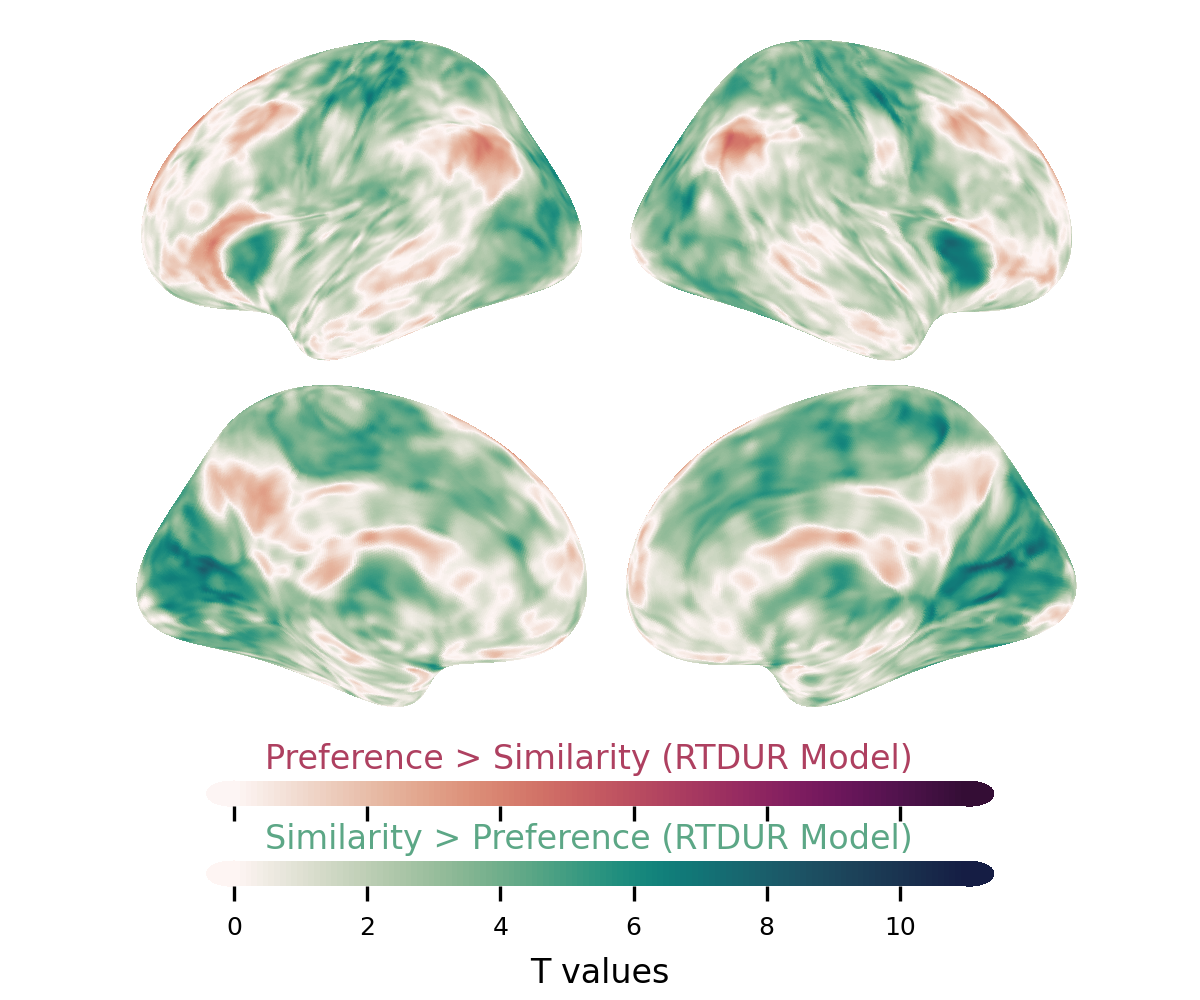

#### Supplementary Figure 4.3: Judgement RT Model: Preference vs Preference-RTDUR, *P_fwe_ < 0.05.*

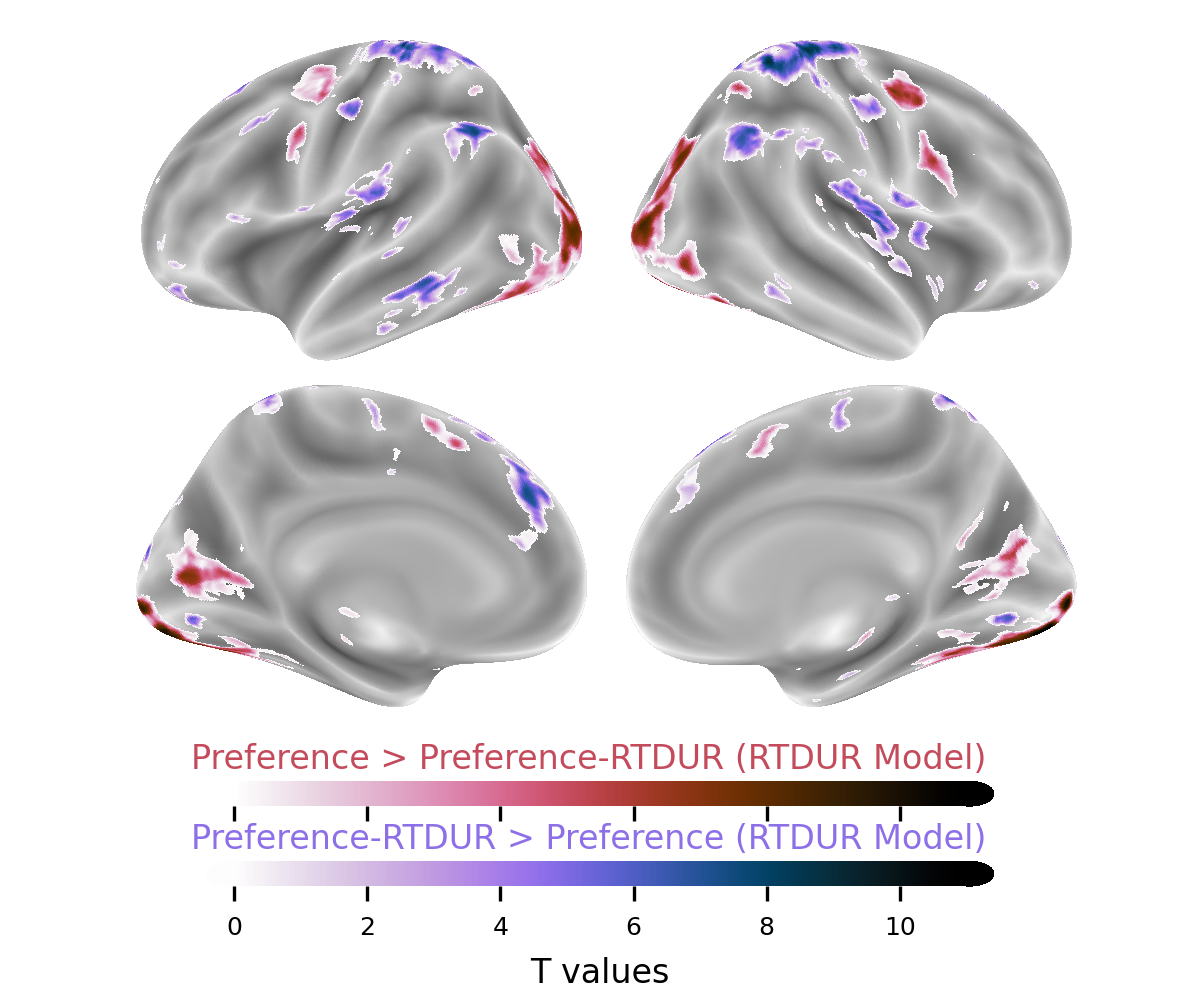

#### Supplementary Table 4.3: Judgement RT Model: Preference vs Preference-RTDUR, *P_fwe_ < 0.05*, Cluster Table.

| Task | Anatomical label | Hemisphere | x | y | z | cluster mean T value | mm3 |
| --- | --- | --- | --- | --- | --- | --- | --- |
| Preference > Preference-RTDUR | 29.38% Occipital Mid  24.15% Fusiform  16.31% Occipital Inf  13.58% Lingual  7.61% Calcarine | R  R  R  R  R | 15.5 | -88.5 | -6.5 | 7.3 | 17560 |
|  | 36.44% Occipital Mid  23.09% Occipital Inf  21.96% Fusiform  8.03% Lingual | L  L  L  L | -32.5 | -84.5 | -8.5 | 7.33 | 16248 |
|  | 84.92% Calcarine  10.46% no label | L  n/a | -24.5 | -66.5 | 9.5 | 6.67 | 2600 |
|  | 85.90% Calcarine  13.68% Cuneus | R  R | 13.5 | -64.5 | 13.5 | 6.5 | 1872 |
|  | 49.46% Frontal Mid 2  46.24% Precentral | R  R | 39.5 | -2.5 | 51.5 | 6.88 | 1488 |
|  | 64.55% Frontal Inf Oper  35.45% Precentral | R  R | 41.5 | 7.5 | 29.5 | 6.17 | 880 |
|  | 76.92% Precentral  15.38% Frontal Sup 2  7.69% Frontal Mid 2 | L  L  L | -44.5 | -6.5 | 47.5 | 6.06 | 728 |
|  | 96.23% Supp Motor Area | L | -6.5 | 5.5 | 61.5 | 6.03 | 424 |
|  | 60.00% Vermis 9  31.11% Vermis 10  8.89% Cerebelum 9 | n/a  n/a  L | 1.5 | -50.5 | -32.5 | 7.12 | 360 |
|  | 60.00% Parietal Inf  20.00% Parietal Sup  17.50% no label | R  R  n/a | 25.5 | -52.5 | 53.5 | 6.04 | 320 |
|  | 100.00% Precentral | L | -46.5 | -0.5 | 37.5 | 5.93 | 288 |
|  | 94.29% Supp Motor Area  5.71% Frontal Sup 2 | R  R | 7.5 | 9.5 | 55.5 | 6.04 | 280 |
|  | 95.00% no label  5.00% Hippocampus | n/a  R | 23.5 | -24.5 | -4.5 | 7.43 | 160 |
|  | 100.00% Cerebelum 6 | L | -8.5 | -76.5 | -18.5 | 6.15 | 80 |
|  | 100.00% no label | n/a | -24.5 | -24.5 | -6.5 | 6.74 | 80 |
| Preference-RTDUR > Preference | 49.77% Postcentral  23.29% Precentral  17.20% Parietal Sup  5.91% Precuneus | R  R  R  R | 17.5 | -28.5 | 61.5 | 6.96 | 8792 |
|  | 53.03% Postcentral  25.14% Parietal Sup  7.08% Precuneus  6.94% Precentral  5.06% Paracentral Lobule | L  L  L  L  L | -28.5 | -40.5 | 67.5 | 6.8 | 5536 |
|  | 48.61% Parietal Inf  25.83% Angular  20.28% no label  5.28% Postcentral | L  L  n/a  L | -54.5 | -60.5 | 41.5 | 6.51 | 2880 |
|  | 50.44% Cerebelum Crus2  45.77% Cerebelum Crus1 | R  R | 37.5 | -70.5 | -42.5 | 6.4 | 2744 |
|  | 88.77% Frontal Sup Medial  7.02% Cingulate Ant | L  L | -2.5 | 35.5 | 37.5 | 6.48 | 2280 |
|  | 67.02% Parietal Inf  28.07% Angular | R  R | 53.5 | -48.5 | 47.5 | 6.29 | 2280 |
|  | 54.88% Rolandic Oper  35.77% Insula  5.28% Temporal Sup | R  R  R | 41.5 | -14.5 | 17.5 | 6.32 | 1968 |
|  | 91.58% Temporal Mid  8.42% Temporal Inf | L  L | -66.5 | -28.5 | -8.5 | 6.39 | 1616 |
|  | 32.39% SupraMarginal  24.65% Temporal Sup  24.65% Rolandic Oper  14.08% Postcentral | L  L  L  L | -50.5 | -24.5 | 17.5 | 6.18 | 1136 |
|  | 56.10% Frontal Sup 2  30.08% Frontal Sup Medial  8.94% Supp Motor Area | L  L  L | -16.5 | 29.5 | 57.5 | 6.14 | 984 |
|  | 57.80% Frontal Sup Medial  42.20% Frontal Sup 2 | R  R | 11.5 | 33.5 | 57.5 | 6.34 | 872 |
|  | 59.26% Cerebelum Crus1  40.74% Cerebelum Crus2 | L  L | -36.5 | -76.5 | -32.5 | 6.11 | 432 |
|  | 79.25% Frontal Mid 2  18.87% no label | L  n/a | -50.5 | 15.5 | 47.5 | 6.29 | 424 |
|  | 92.45% Rolandic Oper  5.66% Heschl | R  R | 59.5 | -2.5 | 9.5 | 6.24 | 424 |
|  | 55.32% Insula  44.68% Rolandic Oper | L  L | -36.5 | -18.5 | 21.5 | 6.2 | 376 |
|  | 100.00% Lingual | L | -8.5 | -80.5 | -4.5 | 6.16 | 376 |
|  | 68.89% Frontal Mid 2  17.78% Frontal Inf Orb 2  11.11% no label | L  L  n/a | -50.5 | 49.5 | -2.5 | 6.11 | 360 |
|  | 86.36% Precentral  11.36% no label | R  n/a | 39.5 | -16.5 | 41.5 | 6.5 | 352 |
|  | 100.00% Lingual | R | 11.5 | -76.5 | -4.5 | 6.23 | 344 |
|  | 66.67% Supp Motor Area  23.81% Supp Motor Area | R  L | -0.5 | -16.5 | 61.5 | 5.92 | 336 |
|  | 97.30% OFCant | L | -32.5 | 41.5 | -14.5 | 6.25 | 296 |
|  | 61.11% Cuneus  33.33% Occipital Sup  5.56% Calcarine | L  L  L | -8.5 | -92.5 | 15.5 | 6.31 | 288 |
|  | 91.43% Caudate  8.57% no label | L  n/a | -14.5 | -2.5 | 17.5 | 6.56 | 280 |
|  | 71.88% Cuneus  21.88% Occipital Sup  6.25% Calcarine | R  R  R | 13.5 | -92.5 | 19.5 | 6.15 | 256 |
|  | 78.12% Postcentral  21.88% no label | L  n/a | -38.5 | -20.5 | 37.5 | 6.66 | 256 |
|  | 73.33% no label  26.67% Frontal Mid 2 | n/a  L | -38.5 | 61.5 | 5.5 | 6.27 | 240 |
|  | 75.00% Rolandic Oper  17.86% no label  7.14% Frontal Inf Oper | L  n/a  L | -62.5 | 1.5 | 5.5 | 6.16 | 224 |
|  | 100.00% Caudate | L | -12.5 | 11.5 | 13.5 | 6.42 | 176 |
|  | 100.00% Insula | R | 35.5 | 5.5 | 9.5 | 6.23 | 168 |
|  | 90.48% Cerebelum Crus1  9.52% Cerebelum Crus2 | R  R | 13.5 | -80.5 | -26.5 | 6.05 | 168 |
|  | 100.00% SupraMarginal | R | 61.5 | -36.5 | 37.5 | 5.85 | 160 |
|  | 84.21% Precentral  15.79% Postcentral | R  R | 41.5 | -24.5 | 67.5 | 6.04 | 152 |
|  | 100.00% SupraMarginal | R | 65.5 | -28.5 | 29.5 | 5.9 | 128 |
|  | 100.00% Supp Motor Area | L | -8.5 | 15.5 | 67.5 | 5.98 | 120 |
|  | 46.67% Frontal Inf Orb 2  33.33% no label  20.00% Temporal Pole Sup | R  n/a  R | 53.5 | 17.5 | -4.5 | 6.16 | 120 |
|  | 85.71% Frontal Sup Medial  14.29% no label | L  n/a | -6.5 | 39.5 | 55.5 | 6 | 112 |
|  | 53.85% Postcentral  38.46% Precentral  7.69% no label | R  R  n/a | 61.5 | -10.5 | 41.5 | 6.03 | 104 |
|  | 76.92% Frontal Inf Tri  23.08% no label | L  n/a | -50.5 | 45.5 | 11.5 | 6.08 | 104 |
|  | 83.33% Temporal Inf  16.67% Temporal Mid | R  R | 59.5 | -42.5 | -14.5 | 5.88 | 96 |
| Cluster defined is 10 voxels | | | | | | | |

#### Supplementary Figure 4.4: Judgement RT Model: Preference vs Preference-RTDUR, Unthreshold T map.

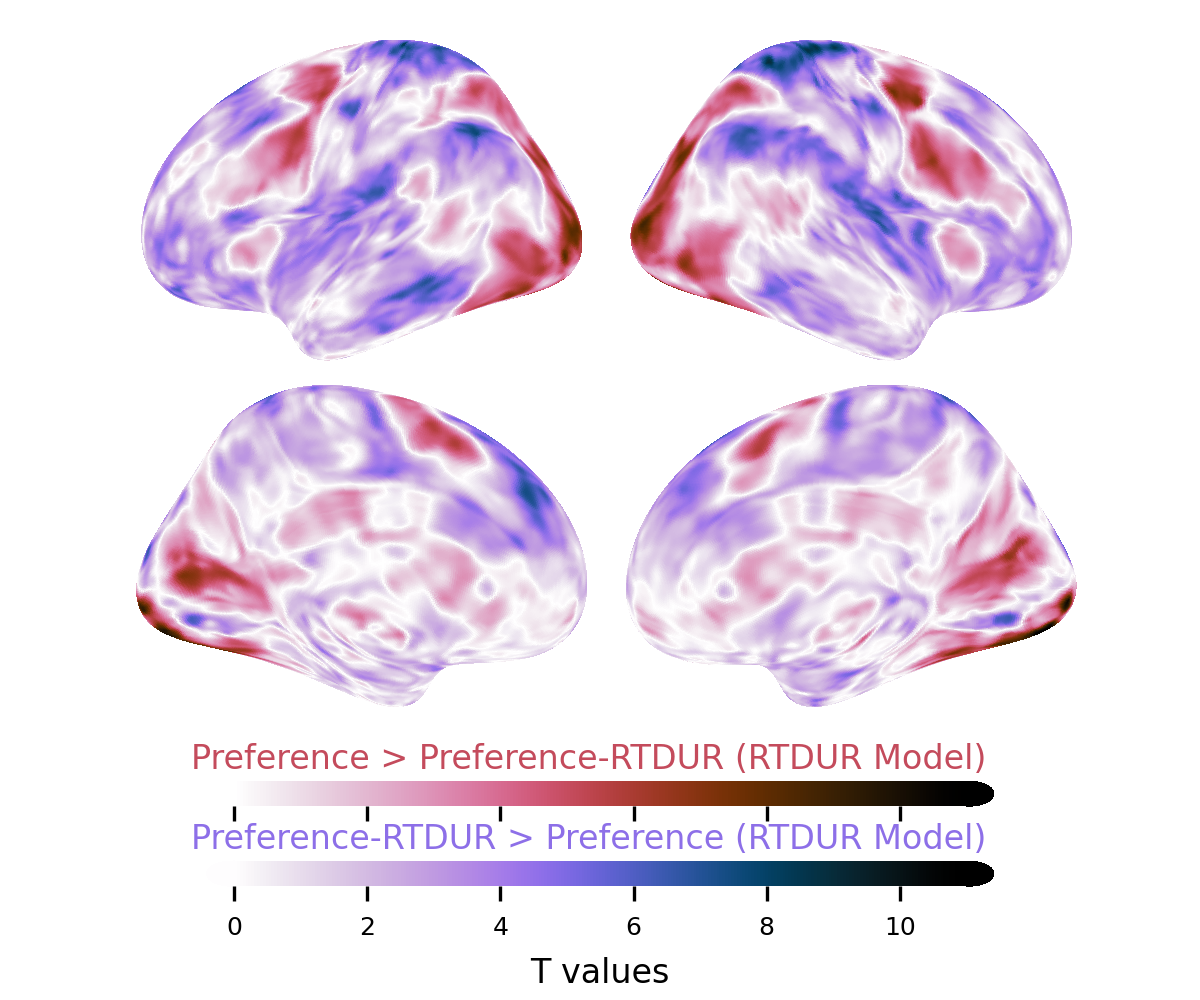

#### Supplementary Figure 4.5: Judgement RT Model: Similarity vs Similarity-RTDUR, *P_fwe_ < 0.05.*

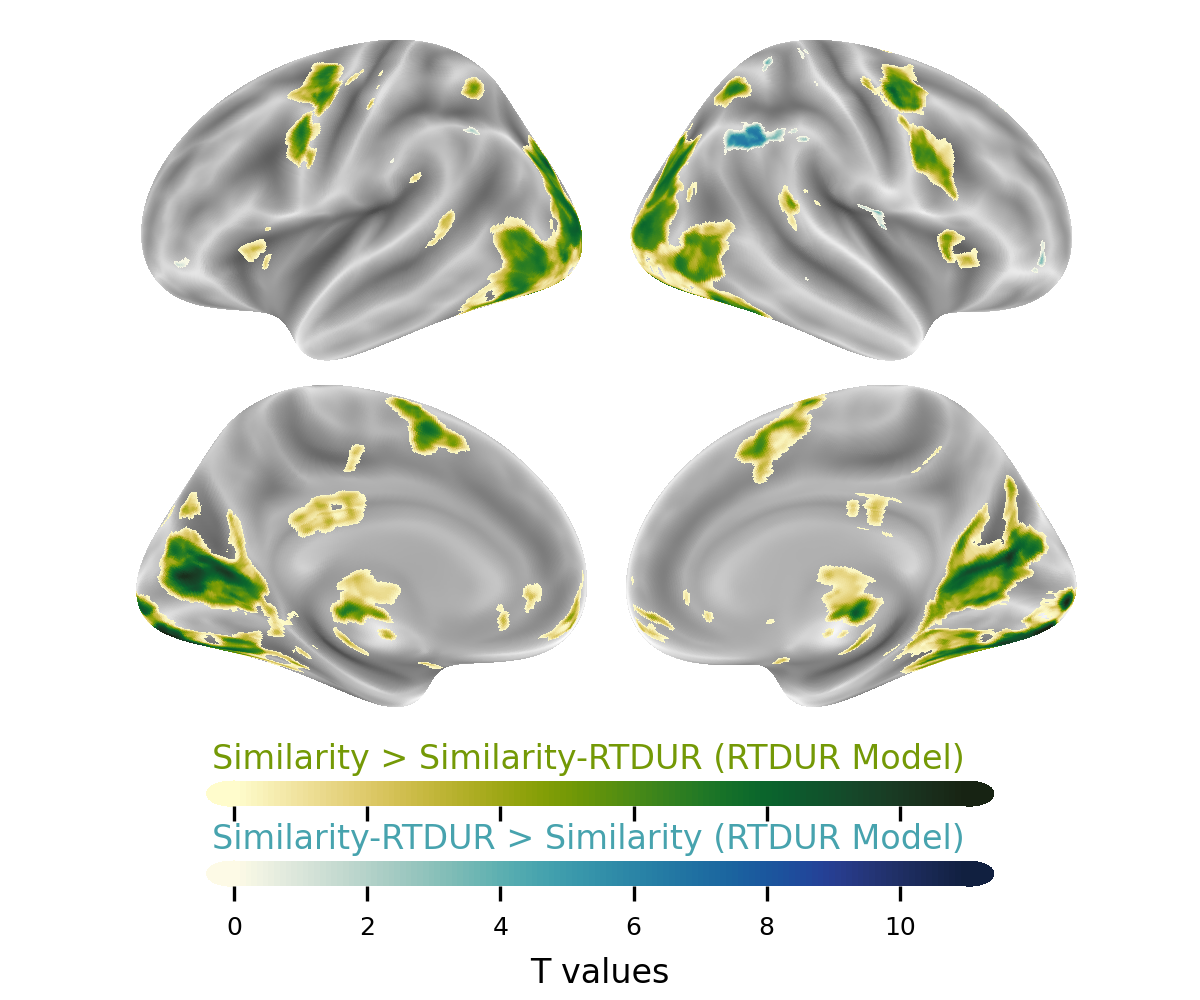

#### Supplementary Table 4.5: Judgement RT Model: Similarity vs Similarity-RTDUR, *P_fwe_ < 0.05*, Cluster Table.

| Task | Anatomical label | Hemisphere | x | y | z | cluster mean T value | mm3 |
| --- | --- | --- | --- | --- | --- | --- | --- |
| Similarity > Similarity-RTDUR | 28.01% Occipital Mid  24.67% Fusiform  13.56% Occipital Inf  11.05% Lingual  5.71% Calcarine  5.20% Temporal Mid | R  R  R  R  R  R | 17.5 | -88.5 | -6.5 | 7.08 | 28016 |
|  | 37.79% Occipital Mid  20.43% Fusiform  18.16% Occipital Inf  7.88% Lingual  5.11% no label | L  L  L  L  n/a | -32.5 | -84.5 | -8.5 | 7.26 | 27096 |
|  | 35.90% Calcarine  29.58% Calcarine  8.19% Cuneus  6.85% Lingual  6.81% Lingual | L  R  R  R  L | 13.5 | -64.5 | 11.5 | 7.25 | 17984 |
|  | 36.04% Thalamus  34.09% Thalamus  29.87% no label | L  R  n/a | 5.5 | -28.5 | -4.5 | 6.66 | 4928 |
|  | 48.60% Putamen  29.37% Caudate  15.38% no label  6.64% Pallidum | R  R  n/a  R | 17.5 | 9.5 | -0.5 | 7.1 | 4576 |
|  | 84.56% Precentral  5.79% Postcentral | L  L | -44.5 | -4.5 | 49.5 | 6.46 | 4560 |
|  | 65.43% Putamen  17.73% Caudate  8.87% no label  7.98% Pallidum | L  L  n/a  L | -18.5 | 11.5 | 1.5 | 7.04 | 4512 |
|  | 47.86% Precentral  37.04% Frontal Mid 2  8.55% Frontal Sup 2  6.55% no label | R  R  R  n/a | 39.5 | -6.5 | 51.5 | 6.75 | 2808 |
|  | 94.23% Supp Motor Area | R | 7.5 | 5.5 | 59.5 | 6.52 | 2496 |
|  | 98.23% Supp Motor Area | L | -6.5 | 3.5 | 57.5 | 6.47 | 2256 |
|  | 61.75% Precentral  36.25% Frontal Inf Oper | R  R | 39.5 | 7.5 | 27.5 | 6.16 | 2008 |
|  | 42.51% Frontal Med Orb  34.01% Frontal Med Orb  12.96% Frontal Sup Medial  6.07% Cingulate Ant | L  R  L  L | -0.5 | 55.5 | -10.5 | 6.33 | 1976 |
|  | 55.40% Vermis 9  20.14% Vermis 10  15.11% Cerebelum 9  9.35% Cerebelum 9 | n/a  n/a  L  R | 1.5 | -52.5 | -34.5 | 7.73 | 1112 |
|  | 49.52% Parietal Sup  29.52% Parietal Inf  16.19% no label | R  R  n/a | 25.5 | -52.5 | 53.5 | 6.48 | 840 |
|  | 100.00% Temporal Sup | R | 65.5 | -40.5 | 13.5 | 5.91 | 384 |
|  | 52.08% Parietal Inf  47.92% Parietal Sup | L  L | -28.5 | -54.5 | 53.5 | 6.26 | 384 |
|  | 72.09% no label  18.60% Hippocampus  9.30% Thalamus | n/a  R  R | 23.5 | -24.5 | -6.5 | 6.77 | 344 |
|  | 79.49% no label  20.51% Hippocampus | n/a  L | -24.5 | -24.5 | -6.5 | 7.59 | 312 |
|  | 87.10% Temporal Mid  6.45% Temporal Sup  6.45% no label | L  L  n/a | -54.5 | -44.5 | 9.5 | 5.83 | 248 |
|  | 100.00% no label | n/a | 1.5 | -30.5 | -44.5 | 6.01 | 232 |
|  | 44.00% no label  32.00% Cingulate Post  24.00% Cingulate Mid | n/a  L  L | -4.5 | -38.5 | 27.5 | 5.75 | 200 |
|  | 100.00% Insula | R | 33.5 | 17.5 | 5.5 | 5.95 | 200 |
|  | 41.67% Cuneus  37.50% Occipital Sup  20.83% no label | L  L  n/a | -6.5 | -88.5 | 41.5 | 6.36 | 192 |
|  | 75.00% Cuneus  25.00% Occipital Sup | L  L | -12.5 | -76.5 | 29.5 | 5.94 | 160 |
|  | 85.00% Cerebelum 10  15.00% Cerebelum 9 | L  L | -18.5 | -38.5 | -46.5 | 6.26 | 160 |
|  | 100.00% no label | n/a | -0.5 | -20.5 | -18.5 | 6.44 | 128 |
|  | 100.00% Precentral | R | 55.5 | -0.5 | 45.5 | 5.83 | 112 |
|  | 100.00% Cerebelum 6 | R | 11.5 | -72.5 | -18.5 | 6.07 | 96 |
|  | 100.00% Cerebelum 10 | R | 19.5 | -38.5 | -46.5 | 5.99 | 88 |
| Similarity-RTDUR > Similarity | 73.42% Parietal Inf  22.07% Angular | R  R | 57.5 | -50.5 | 39.5 | 6.25 | 1776 |
|  | 100.00% Postcentral | R | 27.5 | -40.5 | 69.5 | 6.08 | 256 |
|  | 81.82% Parietal Inf  18.18% no label | L  n/a | -48.5 | -48.5 | 57.5 | 5.98 | 176 |
|  | 60.00% Postcentral  40.00% Parietal Sup | R  R | 49.5 | -38.5 | 59.5 | 5.97 | 80 |
|  | 100.00% no label | n/a | -70.5 | -24.5 | 27.5 | 5.84 | 80 |
| Cluster defined is 10 voxels | | | | | | | |

#### Supplementary Figure 4.6: Judgement RT Model: Similarity vs Similarity-RTDUR, Unthreshold T map.

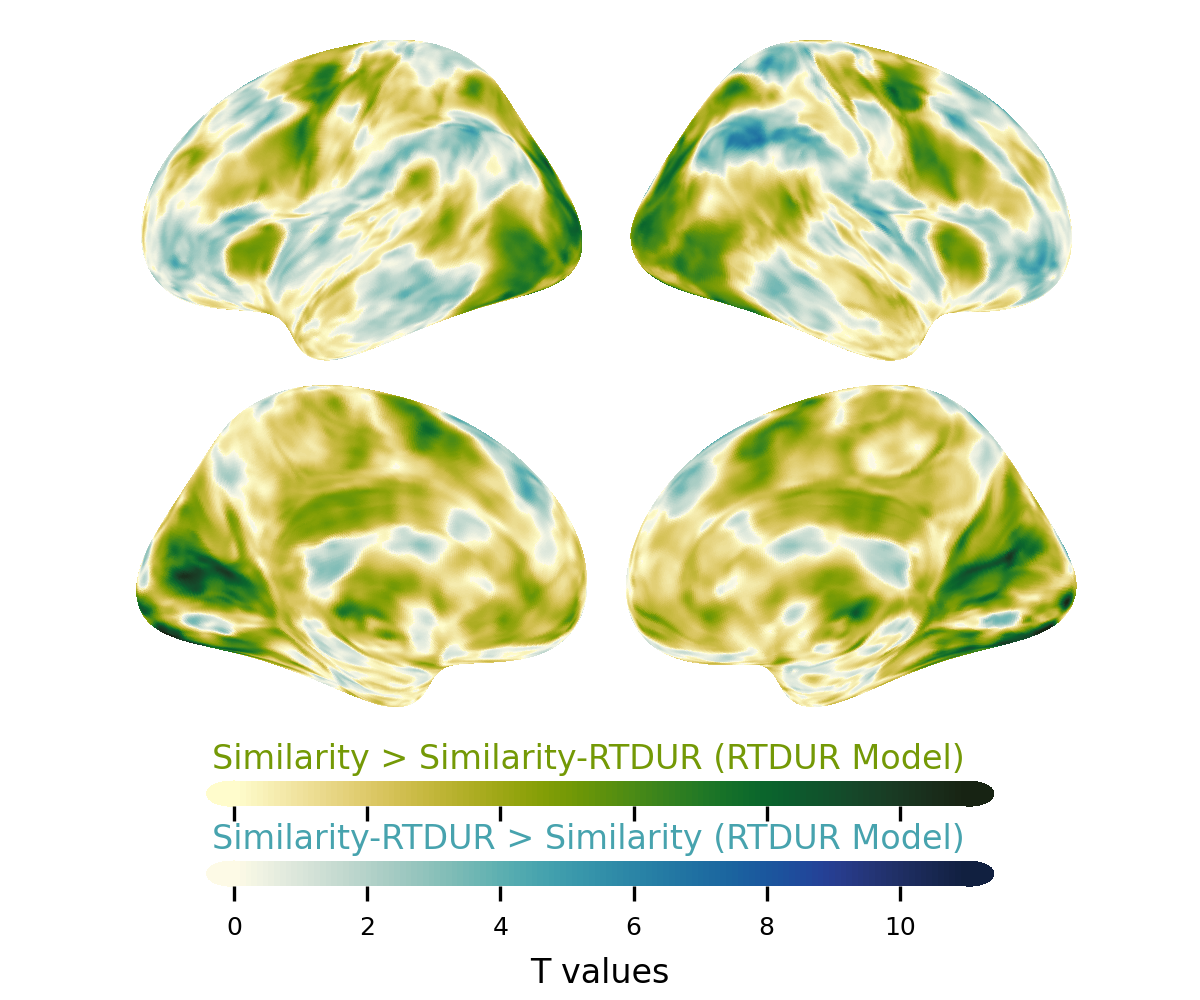

### Supplementary 5: fMRI Judgement Model with Cue-onset.

#### Supplementary Figure 5.1: Judgement Cue-onset Model: Preference Cue-onset vs Similarity Cue-onset, *P_fwe_ < 0.05.*

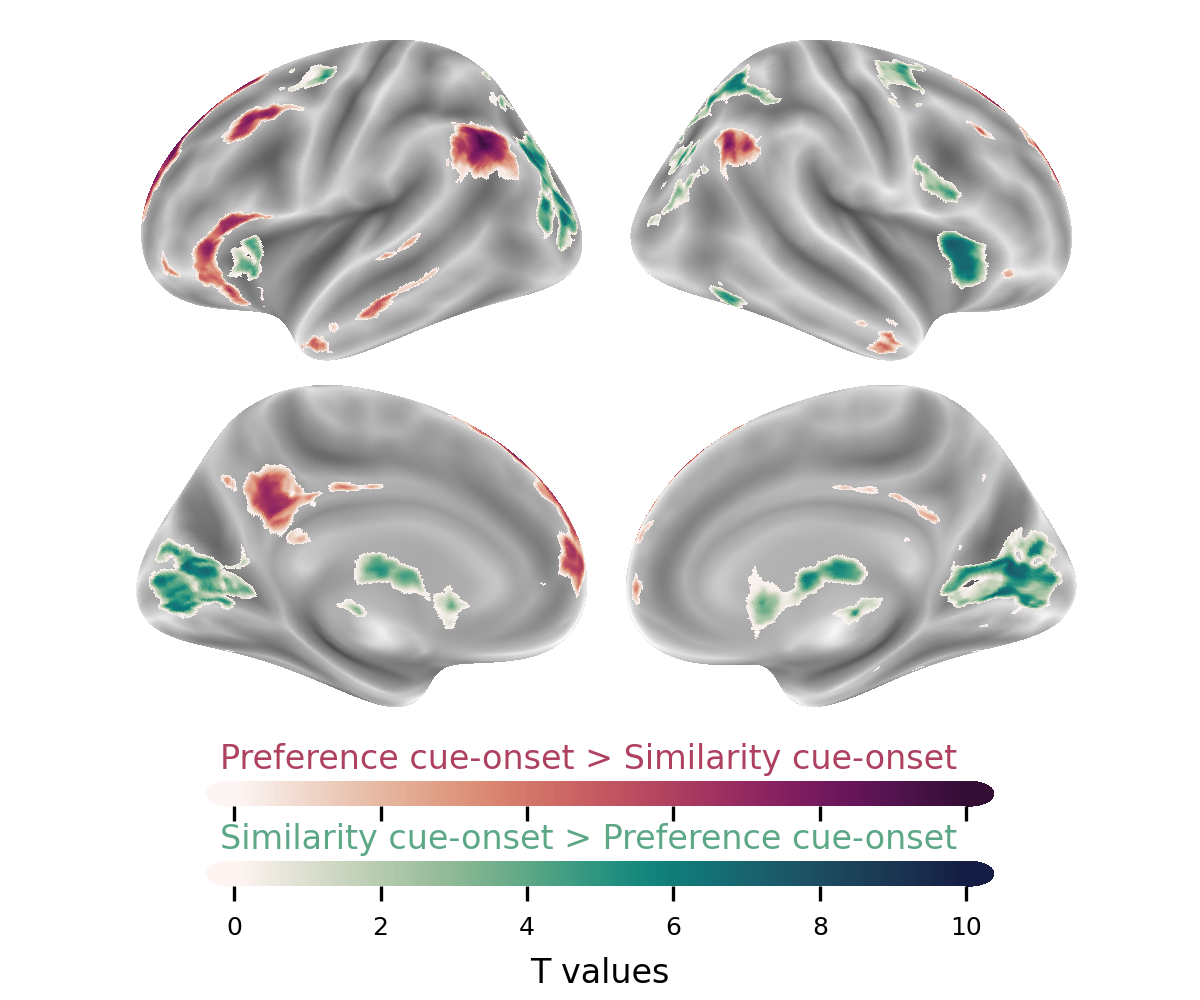

#### Supplementary Table 5.1: Judgement Cue-onset Model :Preference Cue-Onset vs Similarity Cue-Onset, *P_fwe_ < 0.05*, Cluster Table.

| Task condition | Anatomical label | Hemisphere | x | y | z | Cluster mean | mm^3^ |
| --- | --- | --- | --- | --- | --- | --- | --- |
| Preference cue > Similarity cue | 61.35% Frontal Sup 2  32.64% Frontal Sup Medial | L  L | -14.5 | 39.5 | 53.5 | 6.9 | 13432 |
|  | 59.04% Angular  30.23% no label  8.19% Parietal Inf | L  n/a  L | -50.5 | -62.5 | 35.5 | 7.2 | 8496 |
|  | 43.96% Frontal Inf Tri  29.67% Frontal Inf Orb 2  10.20% no label  8.63% OFCpost  6.12% OFClat | L  L  n/a  L  L | -56.5 | 27.5 | -4.5 | 6.72 | 5096 |
|  | 88.08% Frontal Mid 2  9.33% no label | L  n/a | -38.5 | 23.5 | 47.5 | 6.66 | 3088 |
|  | 77.49% Angular  18.71% Parietal Inf | R  R | 51.5 | -60.5 | 35.5 | 6.67 | 2736 |
|  | 65.18% Frontal Sup 2  27.68% Frontal Sup Medial | R  R | 17.5 | 35.5 | 53.5 | 6.52 | 2688 |
|  | 72.26% Frontal Sup 2  27.74% Frontal Sup Medial | R  R | 13.5 | 61.5 | 23.5 | 6.19 | 2336 |
|  | 39.92% Cingulate Post  34.22% Cingulate Mid  20.53% Precuneus  5.32% no label | L  L  L  n/a | -6.5 | -50.5 | 31.5 | 6.41 | 2104 |
|  | 54.55% Cerebelum Crus2  45.45% Cerebelum Crus1 | R  R | 33.5 | -78.5 | -40.5 | 6.13 | 968 |
|  | 95.58% Temporal Mid | L | -66.5 | -20.5 | -12.5 | 6.1 | 904 |
|  | 88.66% Cerebelum Crus2  11.34% Cerebelum Crus1 | L  L | -30.5 | -82.5 | -40.5 | 6.13 | 776 |
|  | 54.26% Temporal Inf  20.21% no label  15.96% Temporal Mid  9.57% Temporal Pole Mid | L  n/a  L  L | -50.5 | -0.5 | -34.5 | 6.24 | 752 |
|  | 53.33% Temporal Mid  45.33% Temporal Inf | R  R | 53.5 | -4.5 | -28.5 | 6.02 | 600 |
|  | 65.12% Frontal Mid 2  18.60% no label  16.28% OFClat | L  n/a  L | -40.5 | 57.5 | -2.5 | 6.05 | 344 |
|  | 100.00% Cerebelum Crus1 | R | 45.5 | -68.5 | -36.5 | 5.96 | 192 |
|  | 100.00% Cerebelum Crus1 | R | 25.5 | -88.5 | -30.5 | 6.03 | 160 |
|  | 95.00% Frontal Mid 2  5.00% no label | R  n/a | 41.5 | 29.5 | 43.5 | 5.94 | 160 |
|  | 100.00% Cingulate Mid | L | -2.5 | -24.5 | 37.5 | 6.22 | 136 |
|  | 100.00% Frontal Sup Medial | R | 7.5 | 55.5 | 3.5 | 5.94 | 88 |
| Similarity cue > Preference cue | 44.25% Calcarine  39.01% Calcarine  6.08% Lingual | R  L  L | -4.5 | -70.5 | 11.5 | 6.33 | 8552 |
|  | 35.57% no label  34.19% Caudate  15.23% Insula  9.16% Putamen | n/a  R  R  R | 9.5 | 7.5 | 5.5 | 6.81 | 7512 |
|  | 93.09% Occipital Mid | L | -28.5 | -76.5 | 21.5 | 6.44 | 4400 |
|  | 56.74% Caudate  21.24% Putamen  18.65% no label | L  L  n/a | -8.5 | 7.5 | 3.5 | 6.84 | 3088 |
|  | 63.16% Parietal Sup  20.65% Parietal Inf  12.55% Occipital Sup | R  R  R | 25.5 | -58.5 | 61.5 | 6.25 | 1976 |
|  | 72.73% Frontal Inf Oper  21.82% Precentral  5.45% Frontal Inf Tri | R  R  R | 53.5 | 11.5 | 37.5 | 6.05 | 880 |
|  | 59.09% Frontal Sup 2  24.55% Precentral  11.82% Frontal Mid 2 | R  R  R | 27.5 | -6.5 | 53.5 | 6.2 | 880 |
|  | 68.37% Vermis 9  11.22% Vermis 10  7.14% Vermis 8  7.14% no label  5.10% Cerebelum 9 | n/a  n/a  n/a  n/a  L | 3.5 | -54.5 | -30.5 | 6.25 | 784 |
|  | 100.00% no label | n/a | 3.5 | -30.5 | -2.5 | 6.79 | 496 |
|  | 98.25% Occipital Mid | L | -44.5 | -80.5 | 13.5 | 6.41 | 456 |
|  | 92.45% Insula | L | -28.5 | 27.5 | 1.5 | 6.07 | 424 |
|  | 80.95% Occipital Mid  19.05% Occipital Sup | R  R | 31.5 | -80.5 | 35.5 | 6.24 | 336 |
|  | 95.12% Occipital Mid | R | 35.5 | -70.5 | 23.5 | 6.14 | 328 |
|  | 100.00% Occipital Mid | R | 43.5 | -78.5 | 19.5 | 6.07 | 320 |
|  | 57.58% Precentral  27.27% Frontal Sup 2  15.15% Frontal Mid 2 | L  L  L | -28.5 | -8.5 | 49.5 | 6.02 | 264 |
|  | 100.00% Temporal Inf | R | 47.5 | -58.5 | -12.5 | 6.22 | 216 |
|  | 52.17% Cerebelum 7b  47.83% Cerebelum 8 | R  R | 9.5 | -74.5 | -42.5 | 6.27 | 184 |
|  | 100.00% Parietal Sup | L | -22.5 | -64.5 | 47.5 | 5.88 | 104 |
|  | 90.00% Occipital Mid  10.00% Temporal Mid | R  R | 47.5 | -72.5 | 23.5 | 6.1 | 80 |
| Cluster defined is 10 voxels | | | | | | | |

#### Supplementary Figure 5.2: Judgement Cue-onset Model: Preference Cue-Onset vs Similarity Cue-Onset, *Unthreshold T map.*

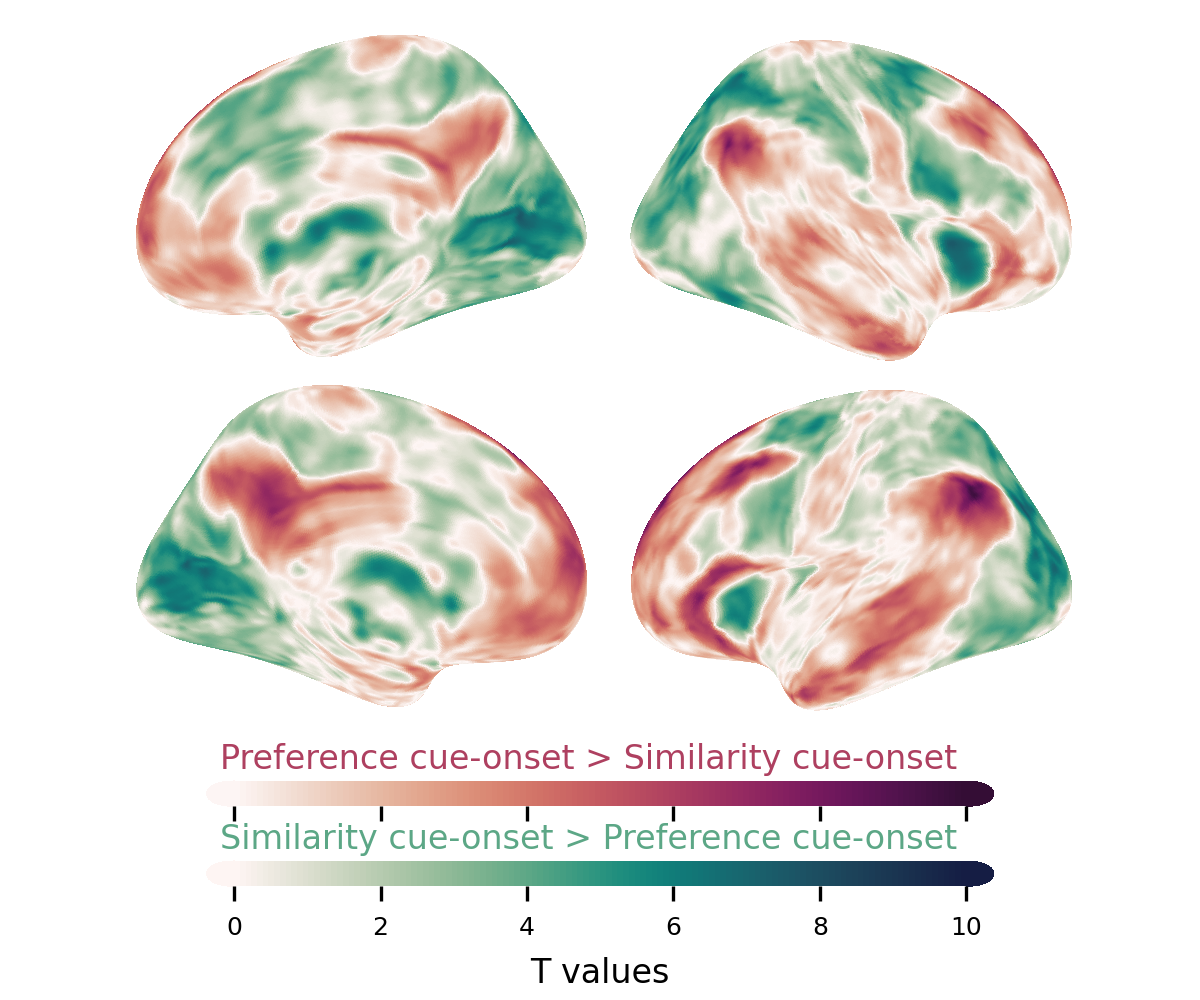

### Supplementary 6: fMRI Condition Model with RTDUR as Covariates.

#### Supplementary Figure 6.1: Condition RT Model: Switch Trial vs Repeated Trial, P_fdr_ <0.01.

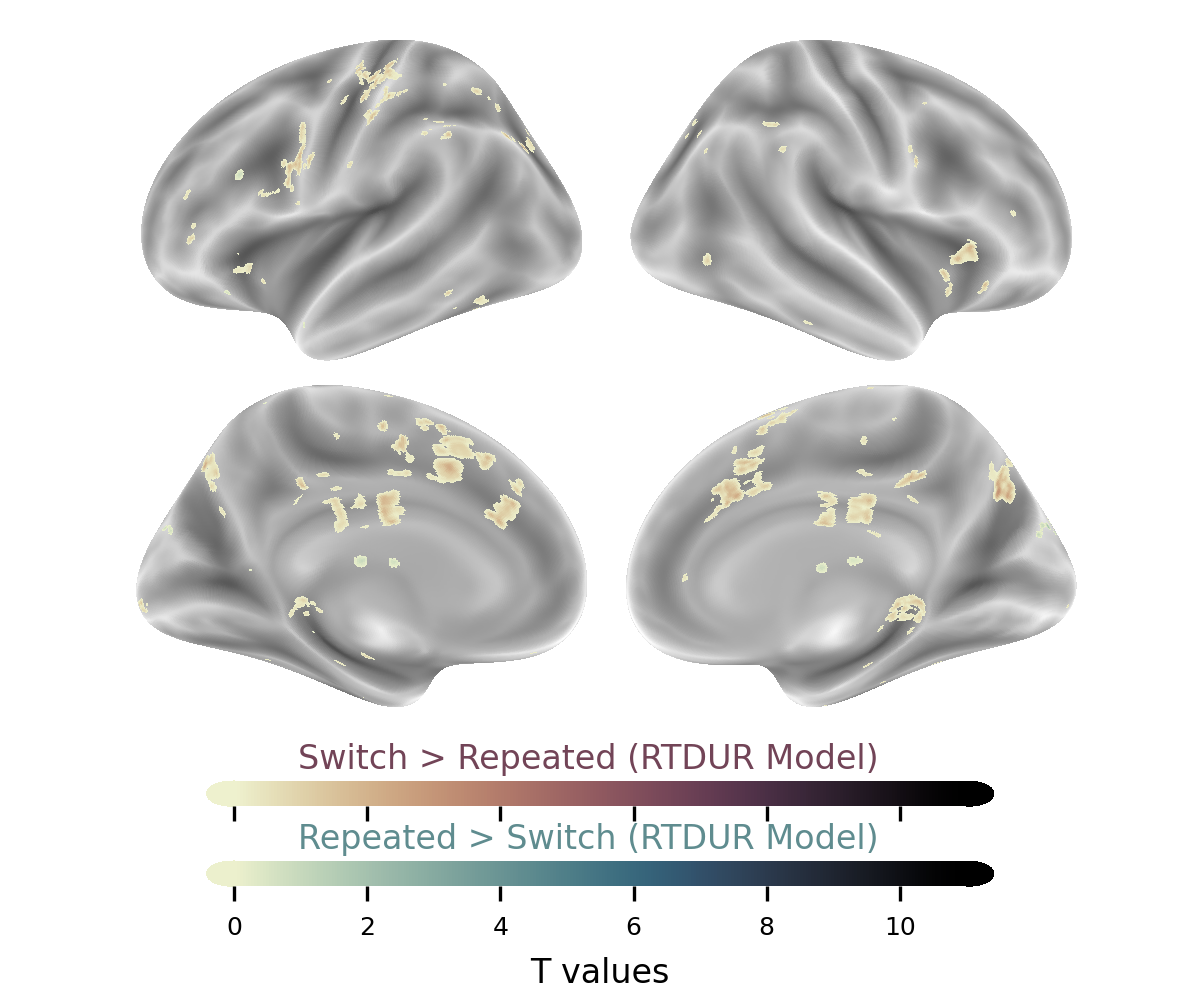

#### Supplementary Table 6.1: Condition RT Model: Switch Trial vs Repeated Trial, P_fdr_ <0.01, Cluster Table.

| Task | Anatomical label | Hemisphere | x | y | z | cluster mean T value | ^mm3^ |
| --- | --- | --- | --- | --- | --- | --- | --- |
| Switch > Repeated | 52.94% Cingulate Mid  47.06% Cingulate Mid | R  L | 1.5 | -40.5 | 37.5 | 3.66 | 136 |
|  | 62.50% Precuneus  37.50% Cuneus | R  R | 11.5 | -68.5 | 35.5 | 4.14 | 128 |
|  | 92.86% Precentral  7.14% Frontal Inf Oper | L  L | -52.5 | 5.5 | 33.5 | 3.75 | 112 |
|  | 91.67% Cingulate Ant  8.33% Frontal Sup Medial | L  L | -8.5 | 27.5 | 25.5 | 3.24 | 96 |
|  | 100.00% Supp Motor Area | R | 7.5 | 5.5 | 61.5 | 3.59 | 80 |
|  | 100.00% Occipital Mid | L | -30.5 | -70.5 | 29.5 | 3.56 | 80 |
|  | 60.00% Precentral  40.00% Frontal Inf Oper | L  L | -46.5 | 5.5 | 27.5 | 4.11 | 80 |
| Cluster defined is 10 voxels | | | | | | | |

#### Supplementary Figure 6.2: Condition RT Model: Switch Trial vs Repeated Trial, Unthreshold T map.

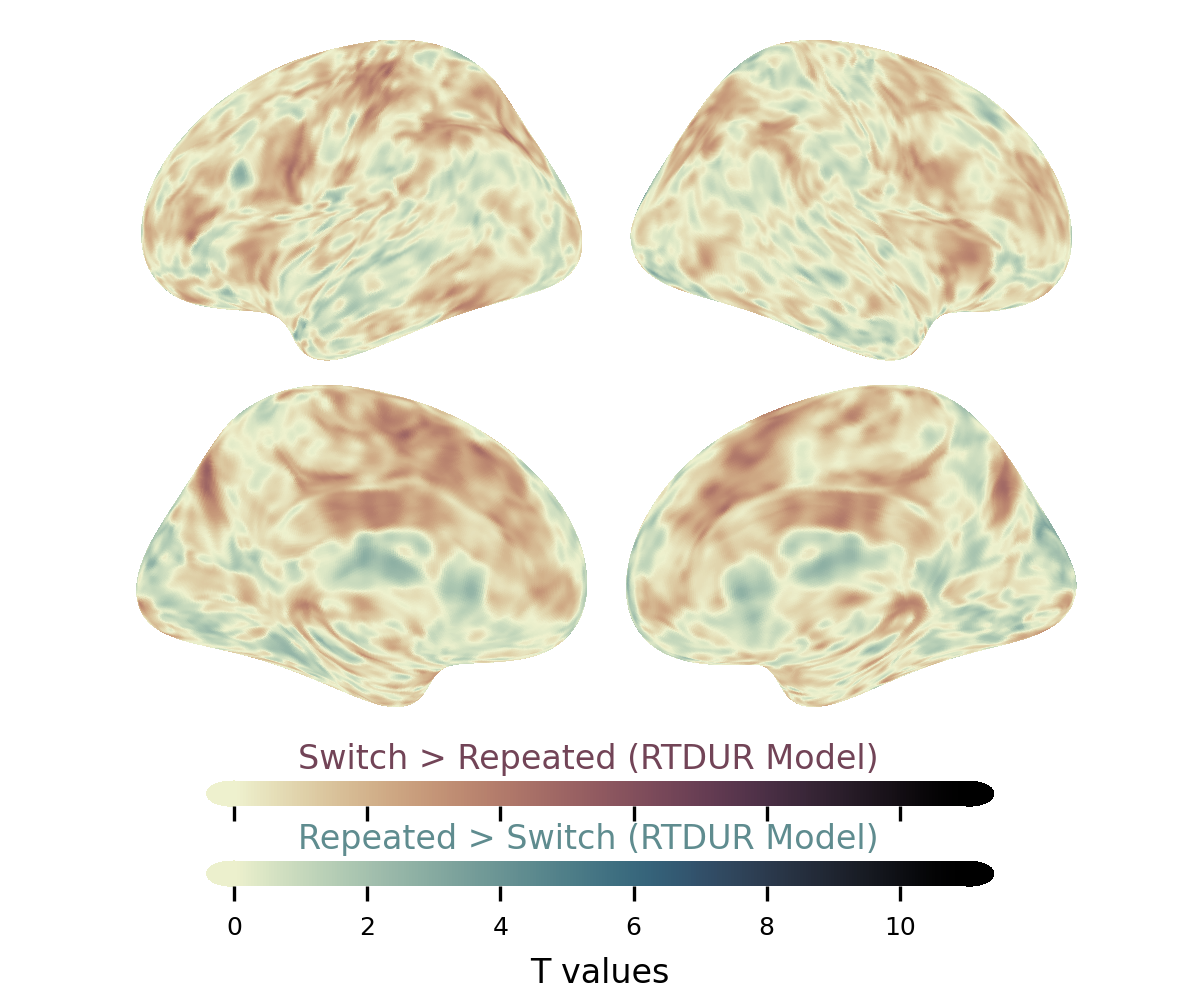

### Supplementary 7: fMRI Condition Model with Cue-onset.

#### Supplementary Figure 7.1: Condition Cue-onset Model: Switch Trial's Cue-onset vs Repeated Trial's Cue-onset, *P_fwe_ < 0.05.*

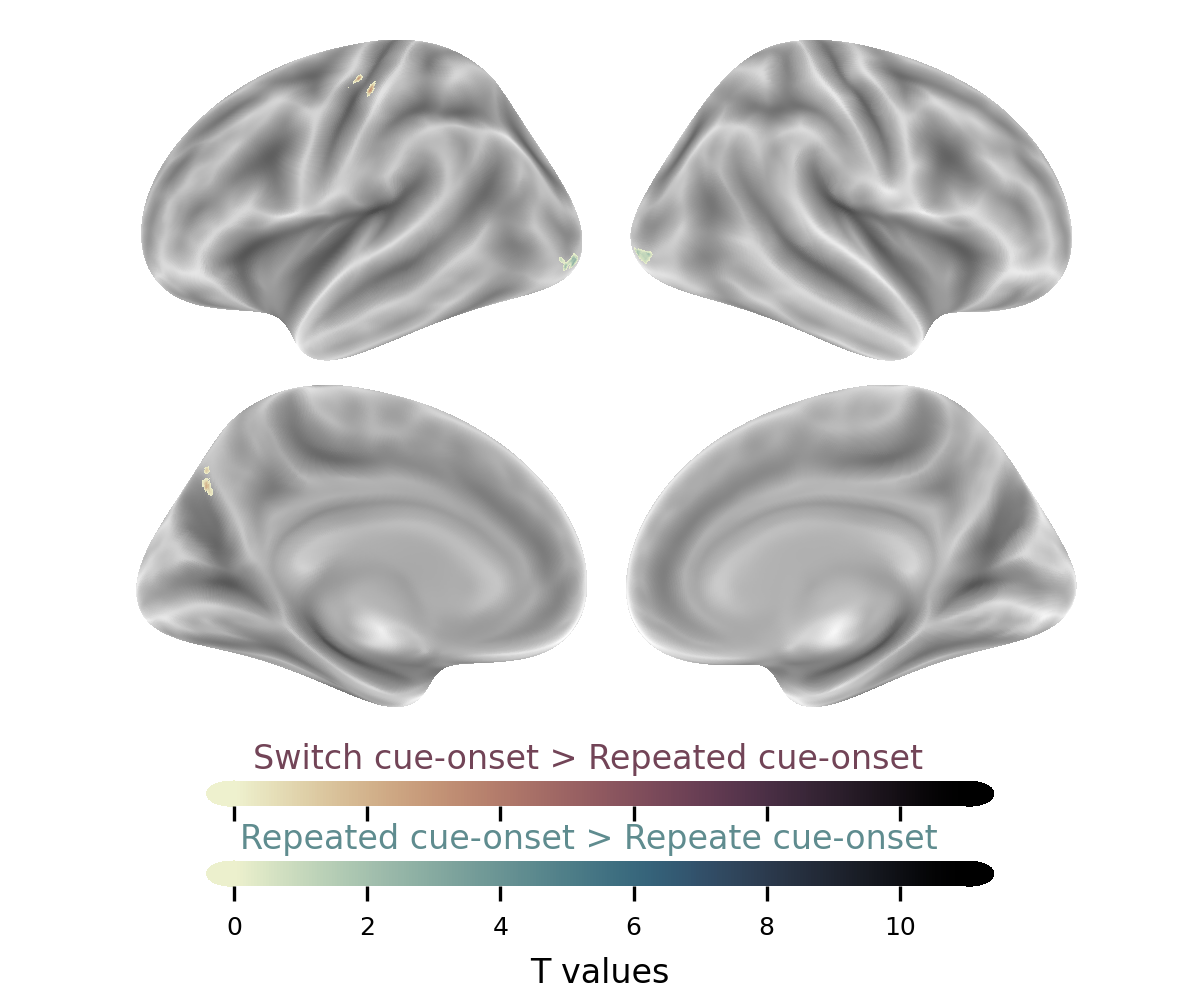

#### Supplementary Table 7.1: Condition Cue-onset Model: Switch Trial's Cue-Onset vs Repeated trial's Cue-Onset, *P_fwe_ < 0.05*, Cluster Table.

| Task | Anatomical label | Hemisphere | x | y | z | cluster mean T value | mm3 |
| --- | --- | --- | --- | --- | --- | --- | --- |
| Repeated cue > Switch cue | 52.00% no label  44.00% Occipital Mid | n/a  L | -28.5 | -100.5 | -10.5 | 5.97 | 200 |
|  | 100.00% Occipital Inf | R | 35.5 | -94.5 | -2.5 | 6.02 | 104 |
| Cluster defined is 10 voxels | | | | | | | |

#### Supplementary Figure 7.2: Condition Cue-onset Model: Switch Trial's Cue-Onset vs Repeated trial's Cue-Onset, *Unthreshold T map.*

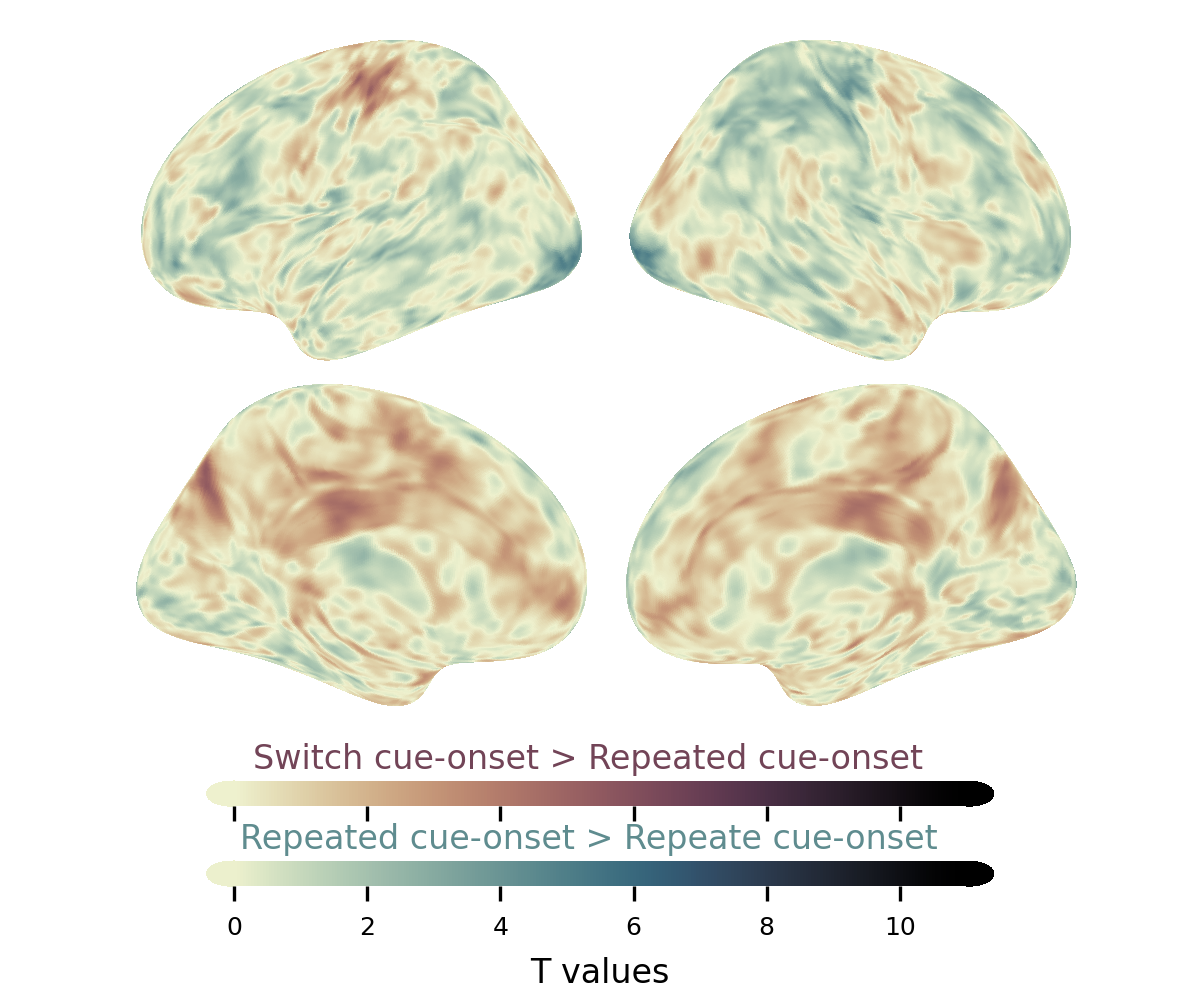

### Supplementary 8: fMRI Judgement X Condition Model with RTDUR as Covariates.

#### Supplementary Figure 8.1: Interaction RT Model: Preference Switch Cost Contrast, P_fdr_ < 0.01

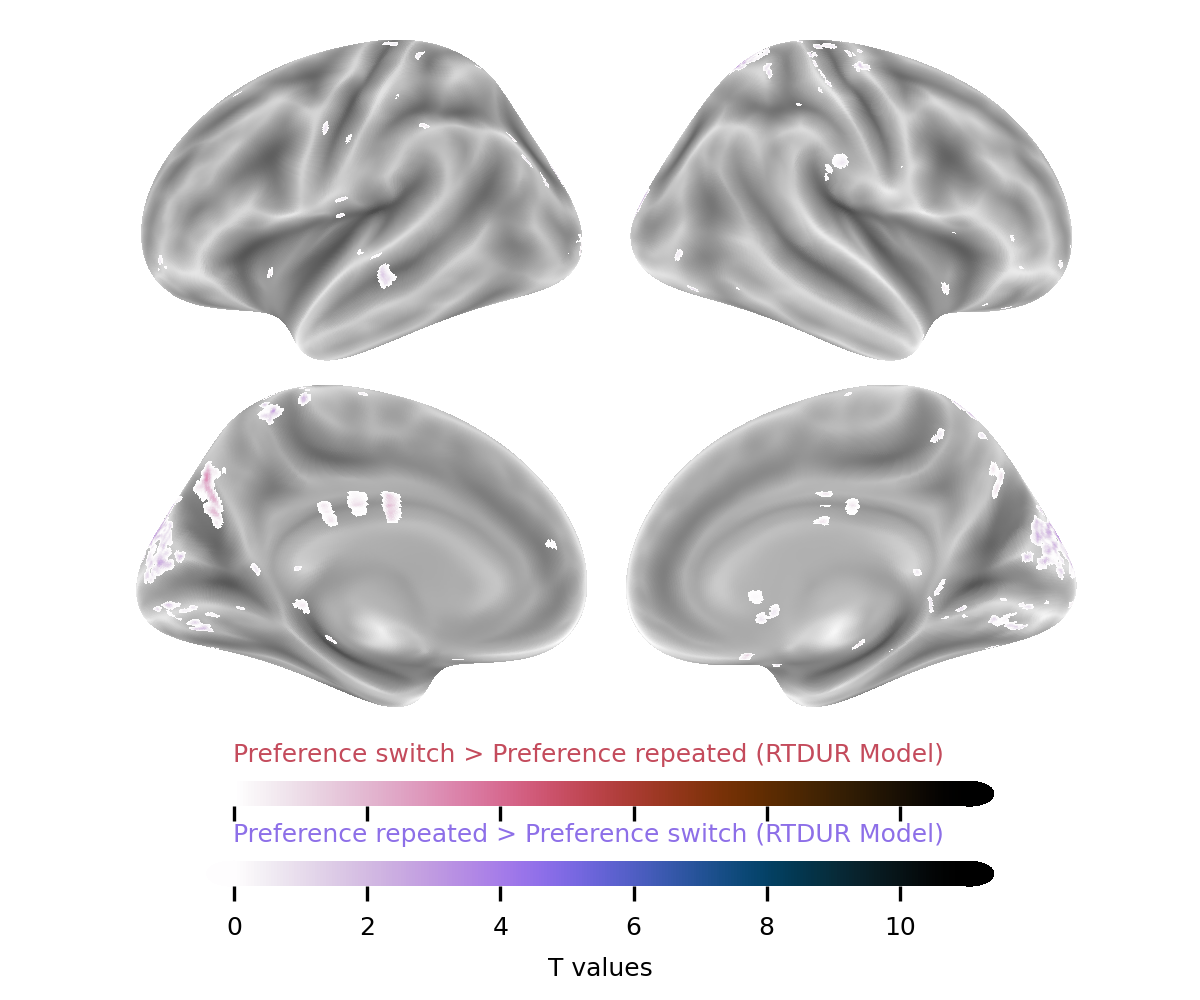

#### Supplementary Table 8.1: Interaction RT Model: Preference Switch Cost Contrast, *P_fdr_ < 0.01*, Cluster Table.

| Task | Anatomical label | Hemisphere | x | y | z | cluster mean T value | mm3 |
| --- | --- | --- | --- | --- | --- | --- | --- |
| Preference switch > Preference repeated | 100.00% Precuneus | L | -8.5 | -72.5 | 37.5 | 3.86 | 248 |
| Preference repeated > Preference switch | 89.80% Cuneus  6.12% Occipital Sup | L  L | -8.5 | -92.5 | 27.5 | 3.86 | 392 |
|  | 90.24% Cuneus  7.32% Cuneus | R  L | 11.5 | -92.5 | 23.5 | 4.05 | 328 |
|  | 83.78% Parietal Sup  16.22% Precuneus | R  R | 17.5 | -58.5 | 67.5 | 3.68 | 296 |
|  | 66.67% Cuneus  33.33% Cuneus | L  R | 3.5 | -86.5 | 25.5 | 3.96 | 120 |
|  | 92.86% Occipital Sup  7.14% Cuneus | R  R | 17.5 | -92.5 | 15.5 | 4.14 | 112 |
|  | 85.71% Calcarine  14.29% Cuneus | L  L | -0.5 | -84.5 | 11.5 | 3.8 | 112 |
|  | 100.00% Precuneus | L | -6.5 | -46.5 | 61.5 | 3.75 | 104 |
| Cluster defined is 10 voxels | | | | | | | |

#### Supplementary Figure 8.2: Interaction RT Model: Preference Switch Cost Contrast, Unthreshold T map.

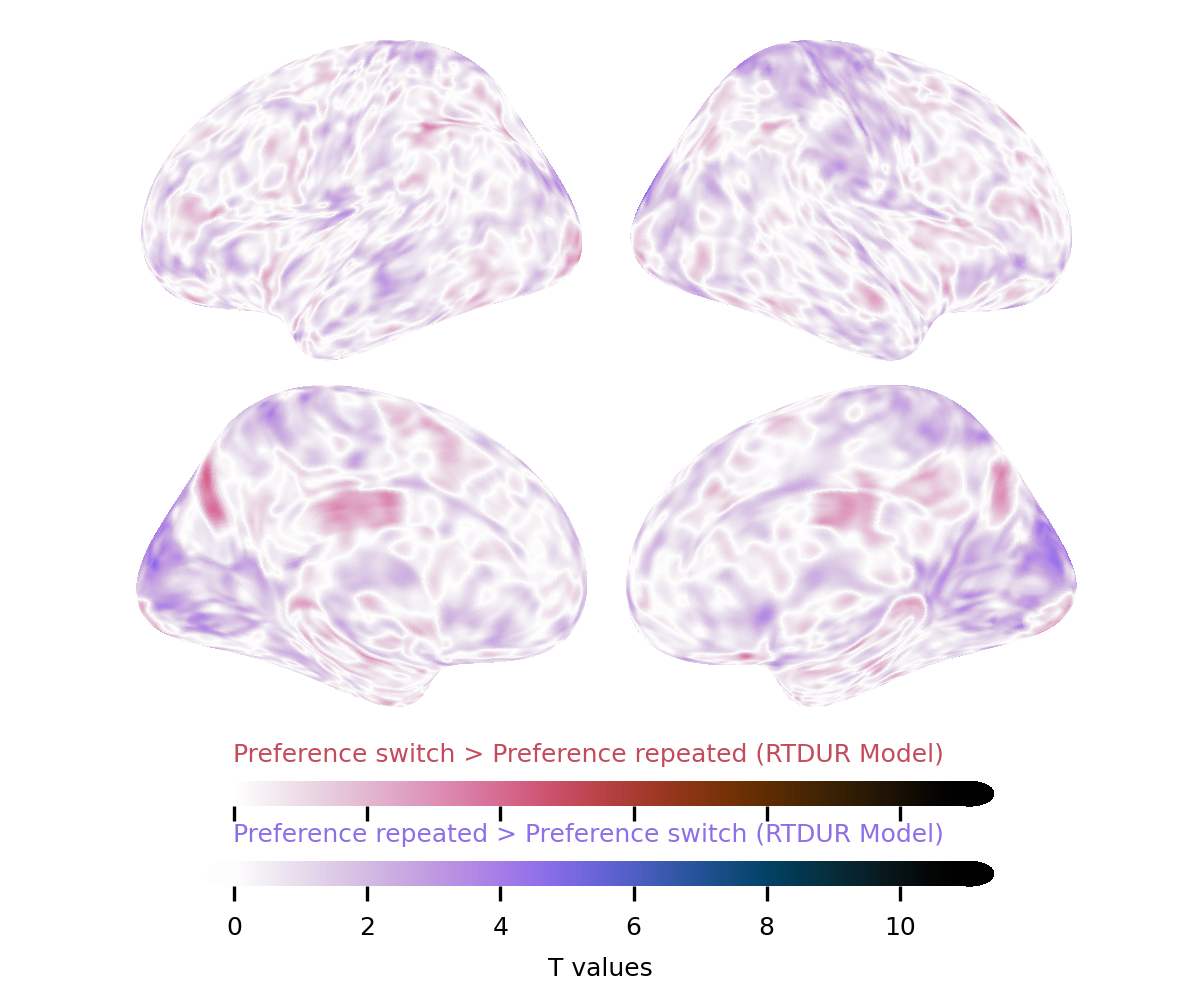

#### Supplementary Figure 8.3: Interaction RT Model: Similarity Switch Cost Contrast, *P_fwe_ < 0.05*.

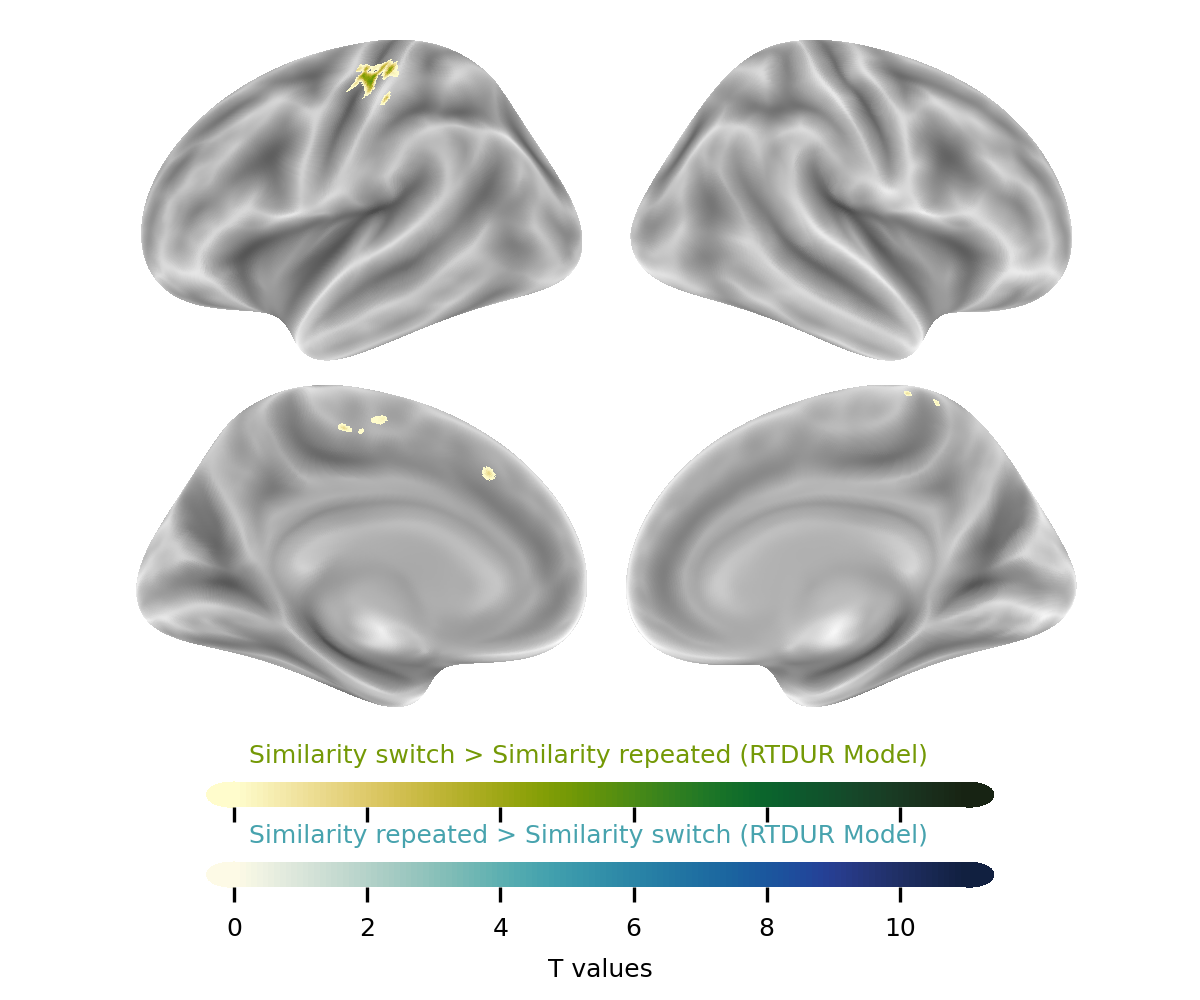

#### Supplementary Table 8.3: Interaction RT Model: Similarity Switch Cost Contrast, *P_fwe_ < 0.05*., Cluster Table.

| Task | Anatomical label | Hemisphere | x | y | z | cluster mean T value | mm3 |
| --- | --- | --- | --- | --- | --- | --- | --- |
| Similarity switch > Similarity repeat | 76.32% Postcentral  23.68% Precentral | L  L | -38.5 | -26.5 | 47.5 | 6.06 | 304 |
|  | 95.00% Precentral  5.00% Postcentral | L  L | -38.5 | -26.5 | 61.5 | 6.17 | 160 |

#### Supplementary Figure 8.4: Interaction RT Model: Similarity Switch Cost Contrast, *Unthreshold T map.*

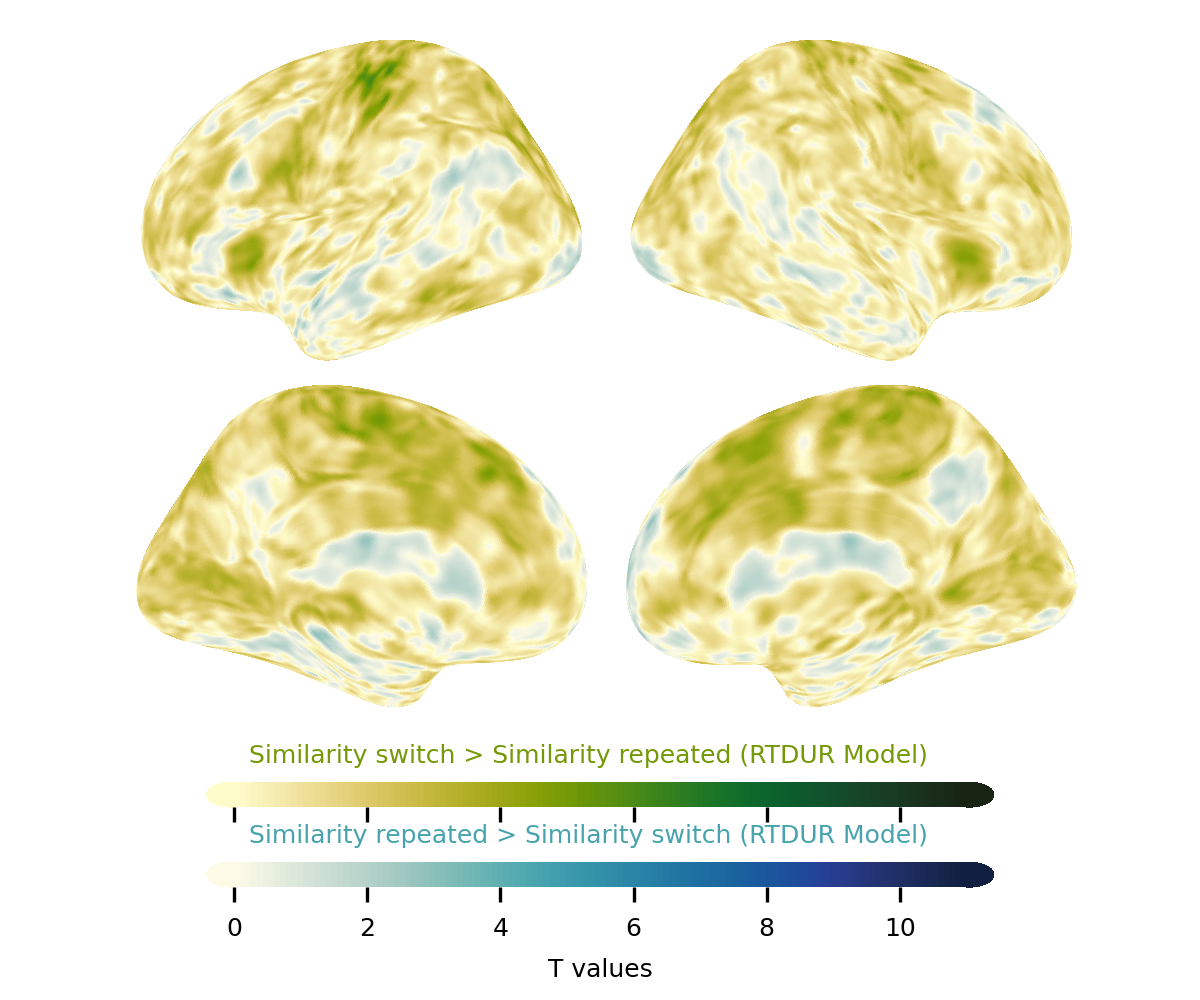

### Supplementary 9: fMRI Judgement X Condition Model with Cue onset.

#### Supplementary Figure 9.1: Interaction Cue-onset Model: Preference Switch Cost Contrast with Cue-onset, *P_fwe_ < 0.05*.

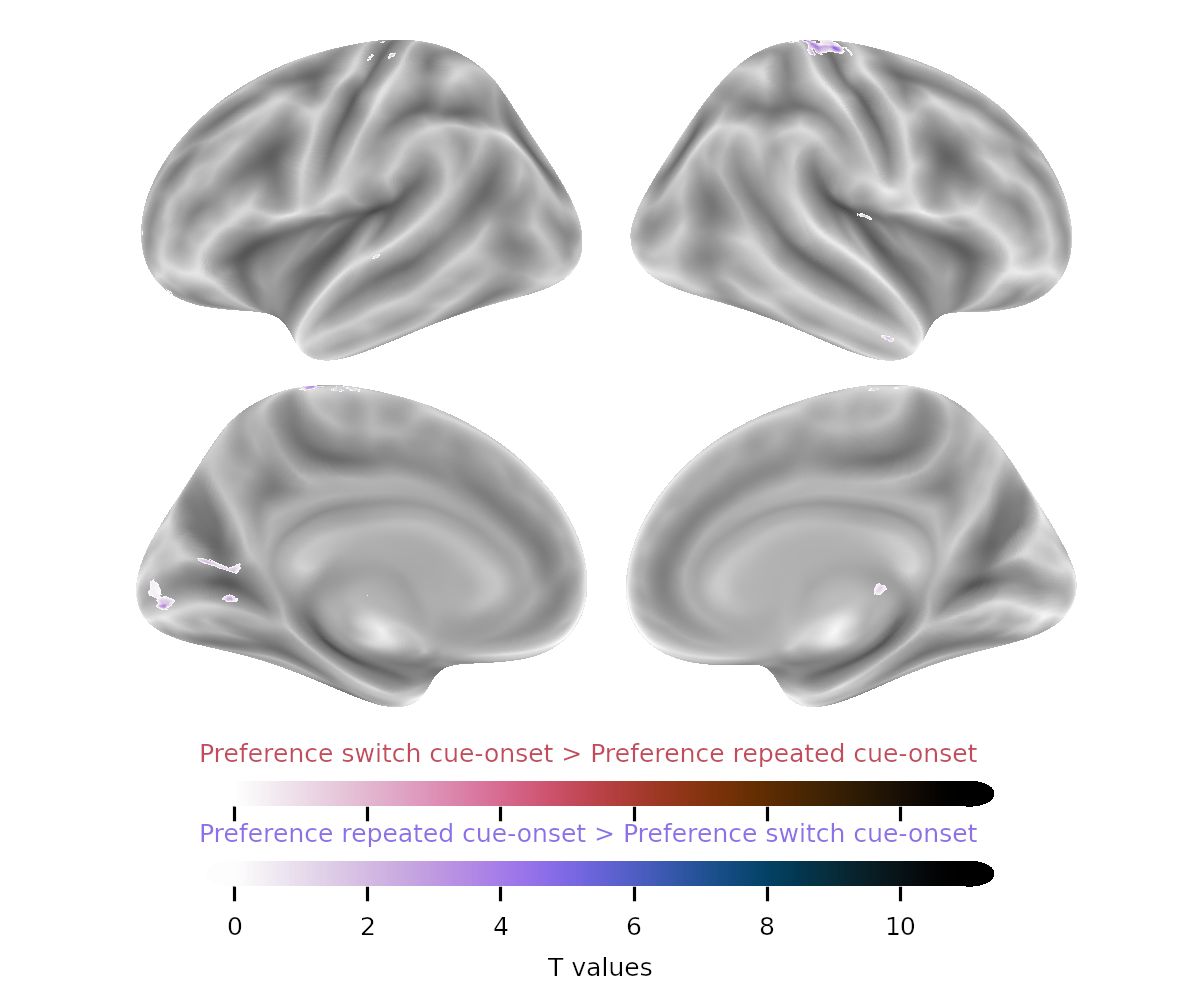

#### Supplementary Table 9.1: Interaction Cue-onset Model: Preference Switch Cost Contrast with Cue-onset, *P_fwe_ < 0.05*, Cluster Table

| Task | Anatomical label | Hemisphere | x | y | z | cluster mean T value | mm3 |
| --- | --- | --- | --- | --- | --- | --- | --- |
| Preference repeated Cue-Onset > Preference switch Cue-Onset | 61.73% Cerebelum Crus1  38.27% Cerebelum Crus2 | R  R | 31.5 | -82.5 | -32.5 | 6.51 | 3136 |
|  | 75.00% Cerebelum Crus2  25.00% Cerebelum Crus1 | L  L | -26.5 | -84.5 | -34.5 | 6.03 | 416 |
|  | 59.18% Precentral  20.41% Postcentral  20.41% no label | R  R  n/a | 11.5 | -32.5 | 73.5 | 5.94 | 392 |
|  | 65.00% Paracentral Lobule  35.00% Precuneus | L  L | -4.5 | -40.5 | 73.5 | 5.96 | 160 |
|  | 92.86% Caudate  7.14% no label | R  n/a | 9.5 | 9.5 | 3.5 | 5.97 | 112 |
|  | 100.00% Calcarine | L | -14.5 | -64.5 | 5.5 | 5.86 | 96 |
|  | 100.00% Thalamus | R | 13.5 | -18.5 | 7.5 | 5.96 | 80 |
|  | 100.00% Calcarine | L | -6.5 | -92.5 | -0.5 | 5.84 | 80 |
| Cluster defined is 10 voxels | | | | | | | |

#### Supplementary Figure 9.2: Interaction Cue-onset Model: Preference Switch Cost Contrast with Cue-onset, *Unthreshold T map.*.

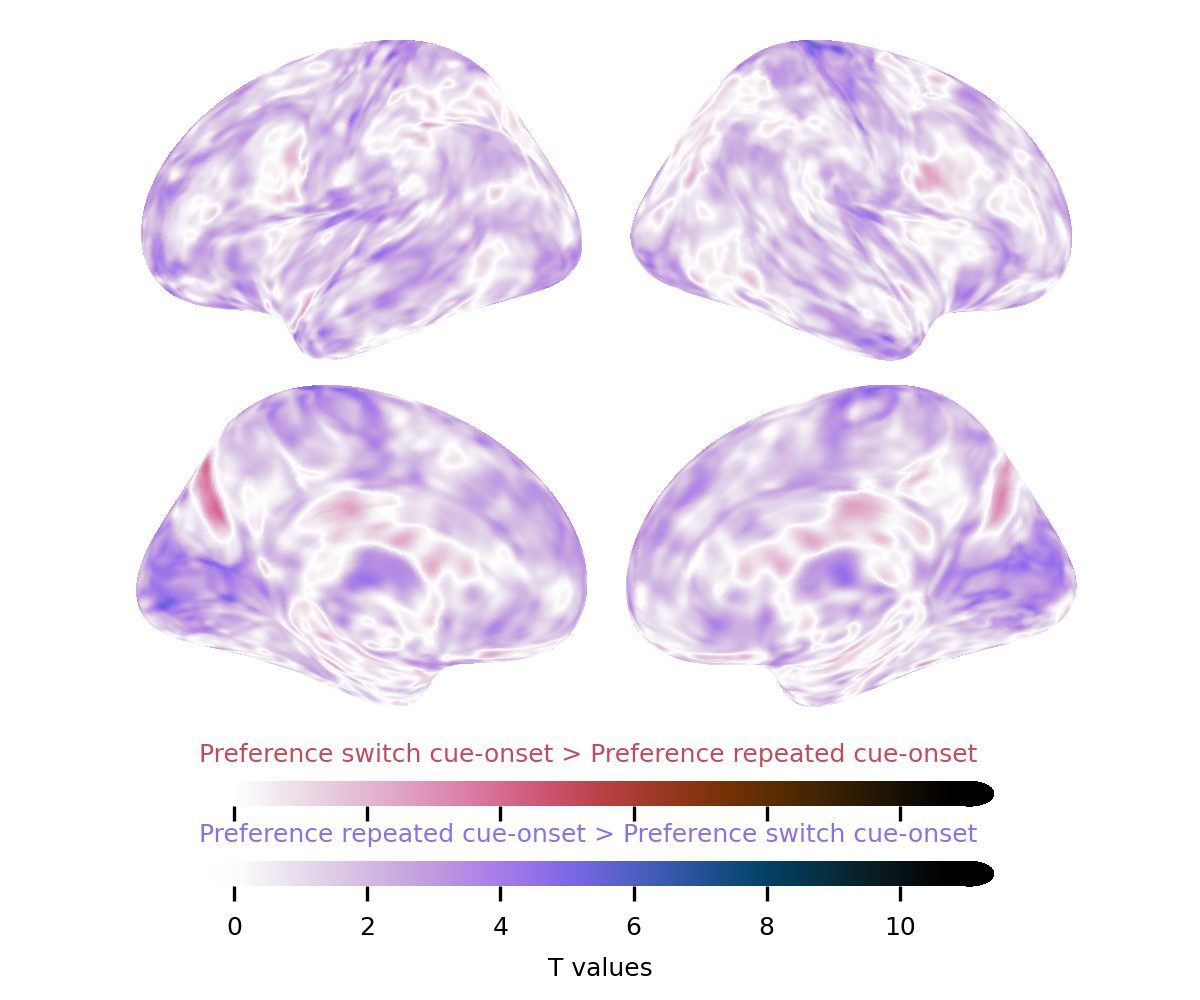

#### Supplementary Figure 9.3: Interaction Cue-onset Model: Similarity Switch Cost Contrast with Cue-onset, *P_fwe_ < 0.05*.

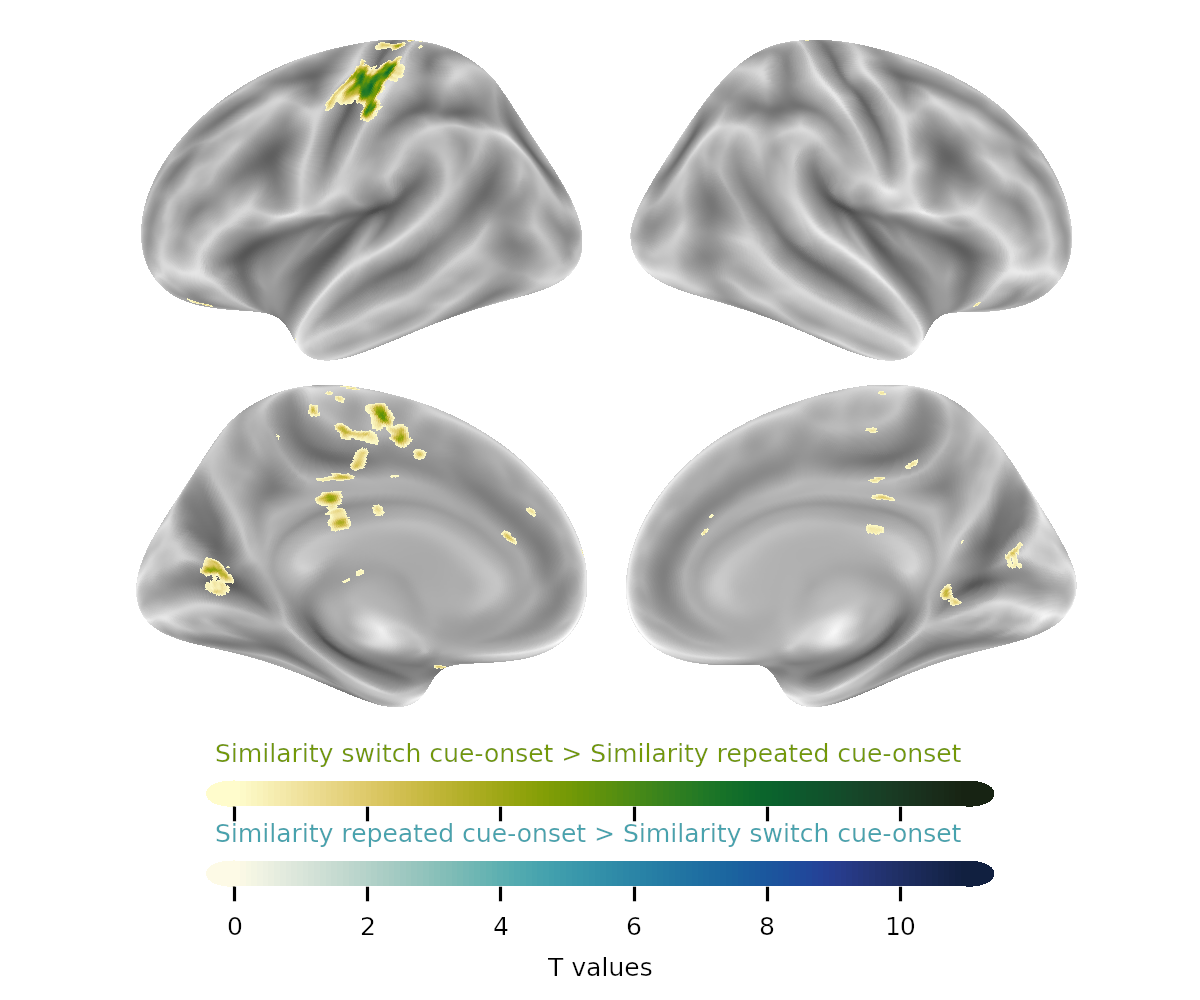

#### Supplementary Table 9.3: Interaction Cue-onset Model: Similarity Switch Cost Contrast with Cue-onset, *P_fwe_ < 0.05*, Cluster Table.

| Task | Anatomical label | Hemisphere | x | y | z | cluster mean T value | mm3 |
| --- | --- | --- | --- | --- | --- | --- | --- |
| Similarity switch onset-cue > Similarity repeated onset-cue | 59.95% Postcentral  39.04% Precentral | L  L | -40.5 | -22.5 | 51.5 | 6.7 | 3176 |
|  | 56.19% Cerebelum Crus1  43.81% Cerebelum Crus2 | R  R | 21.5 | -86.5 | -38.5 | 6.24 | 1808 |
|  | 68.92% Caudate  14.86% Putamen  12.16% no label | R  R  n/a | 11.5 | 15.5 | -8.5 | 5.99 | 592 |
|  | 100.00% Supp Motor Area | L | -4.5 | -14.5 | 59.5 | 6.29 | 216 |
|  | 100.00% Calcarine | L | -20.5 | -66.5 | 9.5 | 5.97 | 208 |
|  | 100.00% Thalamus | L | -10.5 | -18.5 | 7.5 | 6.02 | 176 |
|  | 90.48% Cingulate Mid  9.52% Paracentral Lobule | L  L | -4.5 | -26.5 | 47.5 | 6.04 | 168 |
|  | 60.00% Supp Motor Area  40.00% Cingulate Mid | L  L | -2.5 | -6.5 | 51.5 | 5.99 | 160 |
|  | 50.00% no label  35.00% Olfactory  10.00% Insula  5.00% Putamen | n/a  L  L  L | -24.5 | 9.5 | -14.5 | 6.16 | 160 |
|  | 81.25% Putamen  18.75% Pallidum | L  L | -16.5 | 9.5 | -2.5 | 6.08 | 128 |
|  | 78.57% Postcentral  14.29% no label  7.14% Paracentral Lobule | L  n/a  L | -18.5 | -30.5 | 61.5 | 6.24 | 112 |
|  | 41.67% no label  33.33% Cingulate Mid  25.00% Cingulate Mid | n/a  L  R | -2.5 | -28.5 | 29.5 | 5.98 | 96 |
|  | 27.27% OFCant  27.27% OFCmed  27.27% no label  9.09% OFCpost  9.09% Rectus | L  L  n/a  L  L | -16.5 | 33.5 | -14.5 | 6.27 | 88 |
| Cluster defined is 10 voxels | | | | | | | |

#### Supplementary Figure 9.4: Interaction Cue-onset Model: Similarity Switch Cost Contrast, *Unthreshold T map.*.

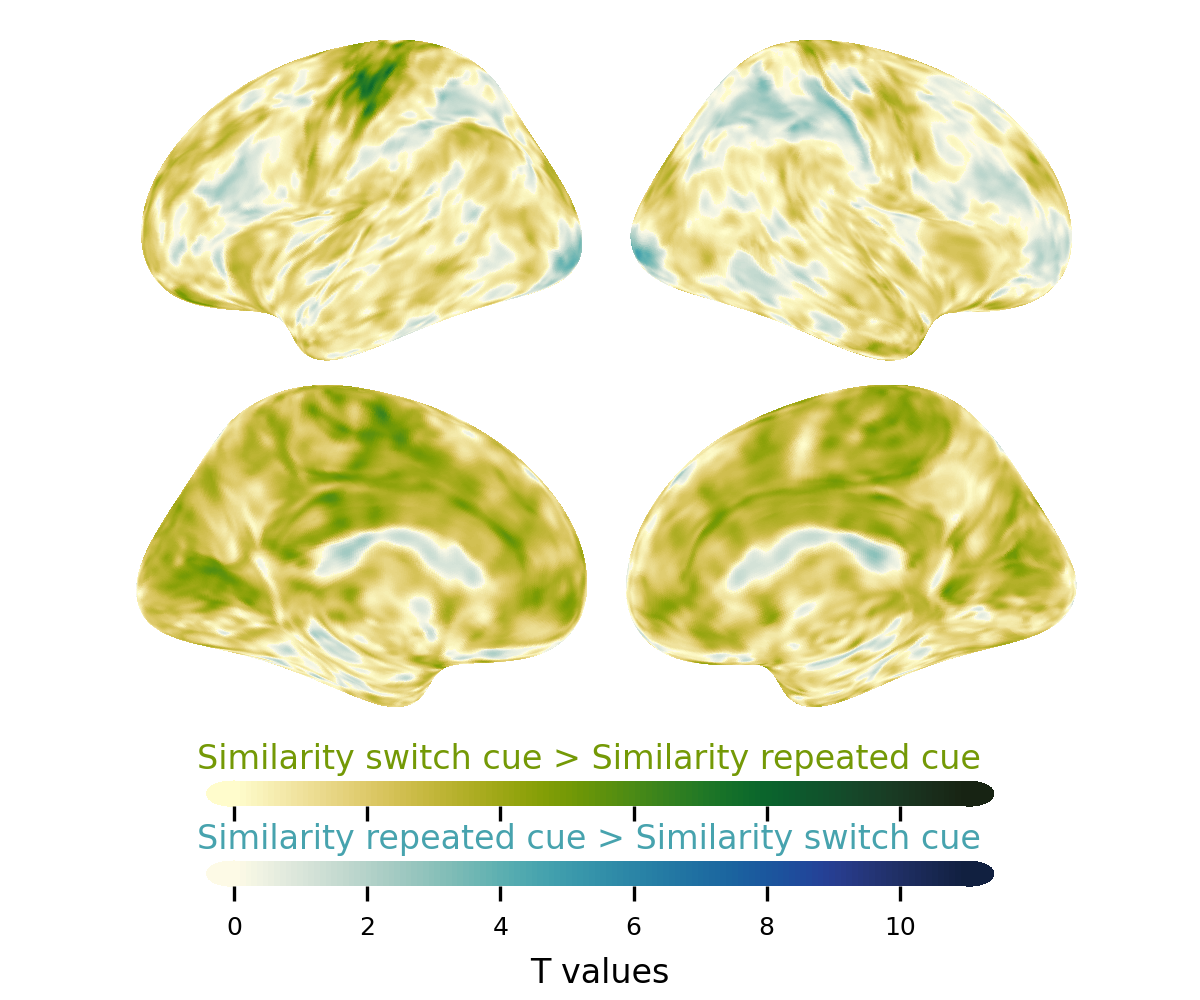

### Supplementary 10: Conjunction Analysis for Switch Trials.

#### Supplementary Figure 10.1:Interaction Model: Similarity Switch Condition, one sample two tailed t-test, *P_fwe_ < 0.05*

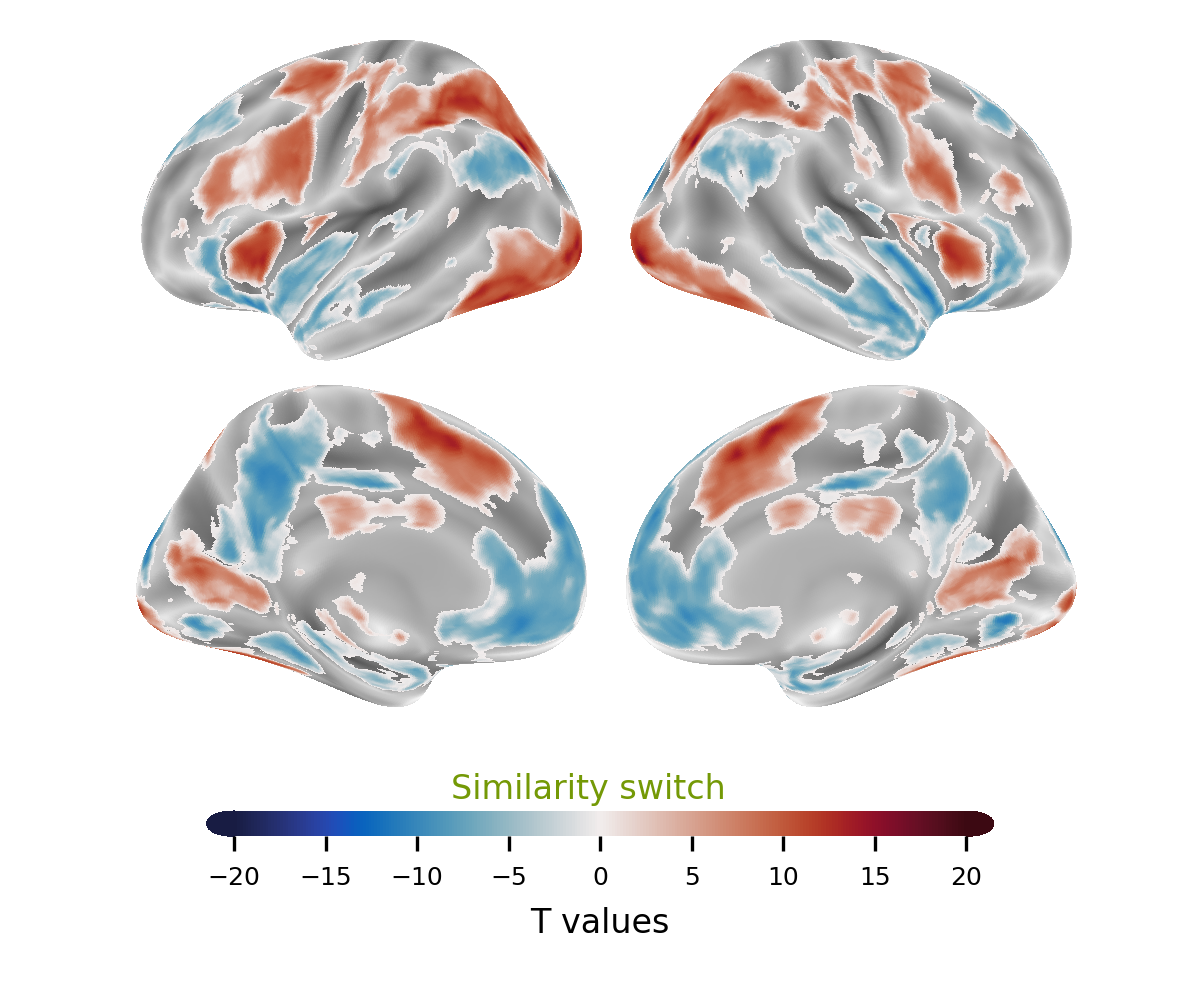

#### Supplementary Table 10.1: Interaction Model: Similarity Switch Condition, one sample two tailed t-test, *P_fwe_ < 0.05,* Cluster Table.

| Task condition | Anatomical label | Hemisphere | x | y | z | cluster mean T value | mm^3^ |
| --- | --- | --- | --- | --- | --- | --- | --- |
| Similarity switch | 15.41% Occipital Inf  14.17% Cerebelum 6  10.25% Fusiform  7.89% Temporal Inf  6.53% Cerebelum Crus1  6.34% Occipital Mid  5.73% Cerebelum 8 | R  R  R  R  R  R  R | 29.5 | -94.5 | -8.5 | 8.23 | 47072 |
|  | 18.95% Occipital Mid  17.71% Occipital Inf  16.45% Fusiform  13.69% no label  9.08% Cerebelum 6  7.87% Cerebelum Crus1  6.88% Temporal Inf | L  L  L  n/a  L  L  L | -22.5 | -98.5 | -4.5 | 8.7 | 35592 |
|  | 37.72% Parietal Inf  23.00% Parietal Sup  11.05% Postcentral  9.31% Occipital Mid  8.03% no label | L  L  L  L  n/a | -46.5 | -42.5 | 49.5 | 8.4 | 32664 |
|  | 20.06% Parietal Sup  17.48% Postcentral  15.42% Parietal Inf  11.67% Precentral  8.78% no label  8.58% Occipital Mid  7.75% Angular | R  R  R  R  n/a  R  R | 33.5 | -68.5 | 31.5 | 7.88 | 27960 |
|  | 47.02% Precentral  19.23% Frontal Inf Tri  10.50% Frontal Inf Oper  8.29% Frontal Sup 2  8.11% Frontal Mid 2  6.21% no label | L  L  L  L  L  n/a | -28.5 | -6.5 | 53.5 | 7.87 | 23464 |
|  | 39.33% Supp Motor Area  31.49% Supp Motor Area  10.88% Cingulate Mid  7.23% Frontal Sup Medial | L  R  R  L | -0.5 | 17.5 | 45.5 | 8.37 | 21032 |
|  | 31.83% Precuneus  19.12% Precuneus  18.92% Cingulate Mid  16.63% Cingulate Mid  7.99% no label | R  L  L  R  n/a | -6.5 | -40.5 | 45.5 | -7.16 | 16112 |
|  | 34.28% Precentral  27.91% Frontal Inf Oper  16.34% Frontal Mid 2  13.37% Frontal Sup 2  6.65% no label | R  R  R  R  n/a | 33.5 | -2.5 | 57.5 | 7.57 | 11552 |
|  | 22.14% Temporal Mid  16.00% Temporal Sup  15.71% Temporal Pole Sup  14.64% Temporal Pole Mid  13.14% Insula  10.36% no label  6.79% OFCpost | R  R  R  R  R  n/a  R | 53.5 | -0.5 | -18.5 | -7.52 | 11200 |
|  | 29.74% Cuneus  23.70% Cuneus  19.42% Occipital Sup  17.35% Occipital Sup | L  R  R  L | -6.5 | -94.5 | 17.5 | -8.12 | 10464 |
|  | 24.48% Cingulate Ant  21.62% Cingulate Ant  20.54% Frontal Med Orb  16.91% Frontal Med Orb  10.27% Frontal Sup Medial | L  R  R  L  R | 7.5 | 35.5 | -4.5 | -6.48 | 10360 |
|  | 39.49% Putamen  31.04% Insula  9.68% Caudate  9.50% no label  6.10% Frontal Inf Tri | L  L  L  n/a  L | -34.5 | 17.5 | 3.5 | 7.75 | 9176 |
|  | 42.22% Calcarine  36.01% Calcarine  7.93% Lingual  5.70% Lingual | L  R  R  L | 11.5 | -62.5 | 7.5 | 7.04 | 7864 |
|  | 36.77% Putamen  22.53% Insula  17.38% no label  17.26% Caudate | R  R  n/a  R | 31.5 | 19.5 | 7.5 | 7.81 | 7136 |
|  | 40.43% Precuneus  26.68% Cuneus  16.53% no label  9.66% Calcarine | L  L  n/a  L | -12.5 | -56.5 | 13.5 | -8.34 | 4888 |
|  | 51.22% no label  31.45% Occipital Mid  15.25% Angular | n/a  L  L | -40.5 | -82.5 | 33.5 | -7.53 | 4248 |
|  | 46.83% Temporal Mid  29.75% Temporal Pole Sup  10.74% Temporal Sup  9.09% Temporal Pole Mid | L  L  L  L | -56.5 | 1.5 | -16.5 | -6.52 | 2904 |
|  | 88.12% Lingual  6.35% Cerebelum 6 | L  L | -12.5 | -80.5 | -4.5 | -7.77 | 2896 |
|  | 32.10% ParaHippocampal  22.73% Lingual  20.17% no label  18.18% Fusiform | L  L  n/a  L | -28.5 | -40.5 | -6.5 | -7.85 | 2816 |
|  | 53.44% Angular  39.06% Occipital Mid | R  R | 49.5 | -72.5 | 25.5 | -7.57 | 2560 |
|  | 41.36% Temporal Sup  38.98% Insula  15.59% no label | L  L  n/a | -40.5 | -6.5 | -6.5 | -7.18 | 2360 |
|  | 94.96% Lingual | R | 13.5 | -76.5 | -4.5 | -7.99 | 2224 |
|  | 36.07% Fusiform  26.48% ParaHippocampal  21.00% Lingual  16.44% no label | R  R  R  n/a | 31.5 | -52.5 | -2.5 | -6.84 | 1752 |
|  | 58.17% Frontal Sup Medial  34.62% Frontal Sup Medial | L  R | 1.5 | 53.5 | 27.5 | -6.37 | 1664 |
|  | 100.00% Thalamus | L | -12.5 | -18.5 | 5.5 | 8.09 | 1272 |
|  | 56.49% no label  33.12% Temporal Sup  10.39% SupraMarginal | n/a  L  L | -68.5 | -28.5 | 15.5 | -6.57 | 1232 |
|  | 50.79% no label  23.02% Cingulate Ant  19.05% Cingulate Mid  5.56% Cingulate Mid | n/a  L  L  R | -4.5 | 5.5 | 29.5 | 6.62 | 1008 |
|  | 36.70% no label  22.94% Olfactory  21.10% Caudate  11.01% Olfactory  8.26% Caudate | n/a  R  L  L  R | -2.5 | 11.5 | -4.5 | -6.48 | 872 |
|  | 78.90% Frontal Inf Orb 2  19.27% Frontal Inf Tri | R  R | 45.5 | 35.5 | -8.5 | -6.38 | 872 |
|  | 100.00% Thalamus | R | 11.5 | -18.5 | 9.5 | 7.18 | 832 |
|  | 69.07% Frontal Sup 2  30.93% Frontal Mid 2 | R  R | 23.5 | 29.5 | 43.5 | -6.48 | 776 |
|  | 51.72% Vermis 9  25.29% Vermis 10  18.39% Cerebelum 9 | n/a  n/a  L | -2.5 | -50.5 | -34.5 | 7.4 | 696 |
|  | 90.36% Cerebelum 9  9.64% no label | R  n/a | 11.5 | -46.5 | -52.5 | -6.56 | 664 |
|  | 70.51% Frontal Inf Tri  29.49% Frontal Mid 2 | L  L | -48.5 | 45.5 | 5.5 | 6.24 | 624 |
|  | 53.97% no label  26.98% Hippocampus  19.05% Thalamus | n/a  L  L | -18.5 | -30.5 | -2.5 | 6.96 | 504 |
|  | 50.79% Frontal Inf Tri  49.21% Frontal Mid 2 | R  R | 49.5 | 33.5 | 23.5 | 6.19 | 504 |
|  | 93.55% Cerebelum 9  6.45% no label | L  n/a | -8.5 | -46.5 | -46.5 | -6.74 | 496 |
|  | 71.67% Insula  28.33% OFCpost | L  L | -28.5 | 17.5 | -18.5 | -6.84 | 480 |
|  | 71.93% no label  22.81% Thalamus  5.26% Hippocampus | n/a  R  R | 23.5 | -26.5 | -2.5 | 7.79 | 456 |
|  | 98.21% Temporal Sup | R | 55.5 | -24.5 | 11.5 | -6.78 | 448 |
|  | 67.31% Rolandic Oper  32.69% Insula | L  L | -40.5 | -6.5 | 13.5 | 6.74 | 416 |
|  | 97.92% Hippocampus | L | -32.5 | -14.5 | -18.5 | -6.26 | 384 |
|  | 46.67% Hippocampus  26.67% Amygdala  24.44% ParaHippocampal | R  R  R | 27.5 | -4.5 | -26.5 | -6.48 | 360 |
|  | 51.52% SupraMarginal  48.48% Angular | R  R | 53.5 | -48.5 | 27.5 | -6.37 | 264 |
|  | 56.25% Paracentral Lobule  43.75% Precuneus | L  L | -6.5 | -40.5 | 71.5 | 6.46 | 256 |
|  | 82.14% Temporal Mid  17.86% Temporal Sup | L  L | -50.5 | -44.5 | 9.5 | 6.3 | 224 |
|  | 100.00% Frontal Mid 2 | L | -36.5 | 55.5 | 11.5 | 6.12 | 200 |
|  | 100.00% Cerebelum 8 | R | 35.5 | -48.5 | -50.5 | 6.1 | 200 |
|  | 100.00% Cingulate Mid | R | 5.5 | 1.5 | 31.5 | 6.78 | 192 |
|  | 69.57% Rolandic Oper  30.43% Insula | R  R | 39.5 | -4.5 | 13.5 | 6.2 | 184 |
|  | 100.00% Postcentral | R | 27.5 | -40.5 | 71.5 | -6.22 | 128 |
|  | 53.33% Postcentral  46.67% Parietal Sup | L  L | -28.5 | -42.5 | 69.5 | -6.21 | 120 |
|  | 92.31% Precuneus  7.69% Postcentral | R  R | 5.5 | -52.5 | 71.5 | -6.13 | 104 |
|  | 91.67% Cerebelum 4 5  8.33% Cerebelum 6 | L  L | -14.5 | -50.5 | -22.5 | 6.1 | 96 |
| Cluster defined is 10 voxels | | | | | | | |

#### Supplementary Figure 10.2: Interaction Model: Preference Switch Condition, one sample two tailed t-test, *P_fwe_ < 0.05.*

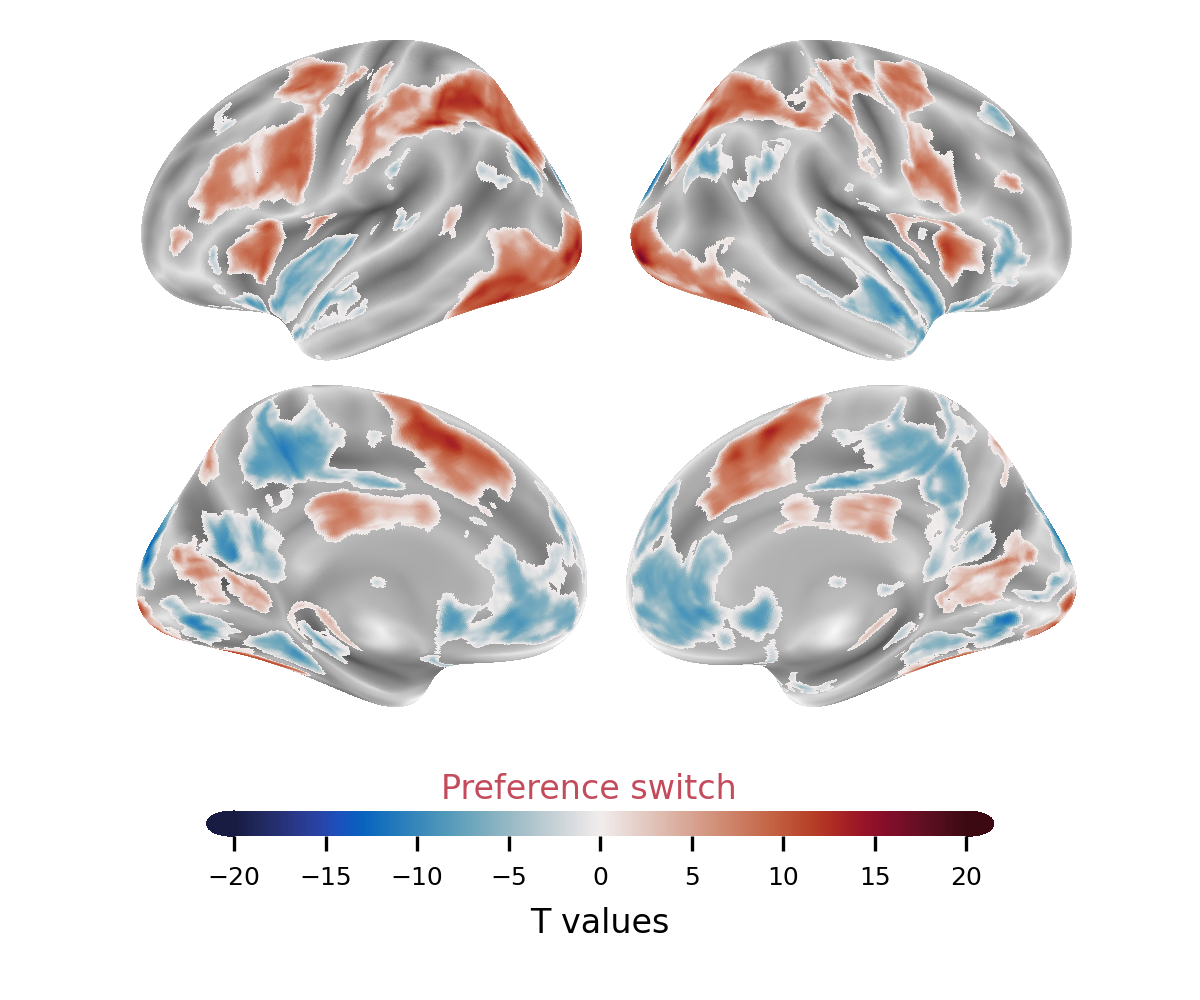

#### Supplementary Table 10.2: Interaction Model: Preference Switch Condition, one sample two tailed t-test, *P_fwe_ < 0.05,* Cluster Table.

| Task condition | Anatomical label | Hemisphere | x | y | z | cluster mean T value | mm3 |
| --- | --- | --- | --- | --- | --- | --- | --- |
| Preference switch | 10.46% Frontal Sup Medial  7.84% Frontal Sup Medial  7.32% no label  6.13% Cingulate Ant  5.94% Frontal Sup 2  5.61% Temporal Mid | L  R  n/a  L  L  R | 41.5 | -6.5 | -12.5 | -7.12 | 107808 |
|  | 7.89% Occipital Inf  7.70% Occipital Mid  7.45% Cerebelum 6  7.02% Cerebelum 6  6.96% Occipital Inf  6.91% no label  6.40% Fusiform  5.73% Fusiform | R  L  R  L  L  n/a  L  R | 29.5 | -94.5 | -8.5 | 8.38 | 90816 |
|  | 16.23% Precentral  14.66% Parietal Sup  14.11% Postcentral  10.32% Parietal Inf  8.61% no label  7.90% Frontal Inf Oper  6.05% Occipital Mid | R  R  R  R  n/a  R  R | 31.5 | -68.5 | 31.5 | 8.01 | 45880 |
|  | 24.52% Precuneus  20.41% Precuneus  17.33% Cingulate Mid  14.18% Cingulate Mid  8.62% no label | L  R  L  R  n/a | -12.5 | -56.5 | 13.5 | -7.47 | 33512 |
|  | 38.72% Parietal Inf  25.39% Parietal Sup  11.49% Occipital Mid  8.00% Postcentral  7.97% no label | L  L  L  L  n/a | -28.5 | -70.5 | 29.5 | 8.62 | 30496 |
|  | 48.42% Precentral  18.39% Frontal Inf Tri  11.76% Frontal Inf Oper  8.86% Frontal Sup 2  6.41% Frontal Mid 2  5.75% no label | L  L  L  L  L  n/a | -30.5 | -4.5 | 57.5 | 7.95 | 21840 |
|  | 39.65% Supp Motor Area  32.30% Supp Motor Area  10.21% Cingulate Mid  7.47% Frontal Sup Medial | L  R  R  L | -8.5 | 11.5 | 51.5 | 8.57 | 19592 |
|  | 43.30% no label  34.32% Angular  7.23% Occipital Mid  6.99% Temporal Sup | n/a  L  L  L | -48.5 | -72.5 | 37.5 | -7.32 | 13728 |
|  | 30.65% Cuneus  23.79% Cuneus  18.95% Occipital Sup  16.77% Occipital Sup | L  R  R  L | -6.5 | -94.5 | 17.5 | -7.96 | 9920 |
|  | 62.12% Angular  13.95% SupraMarginal  7.32% Parietal Inf  6.03% Occipital Mid | R  R  R  R | 49.5 | -70.5 | 41.5 | -6.84 | 8088 |
|  | 40.96% Calcarine  39.74% Calcarine  6.81% Lingual | L  R  R | 13.5 | -64.5 | 9.5 | 7.1 | 7872 |
|  | 27.73% Insula  25.06% Putamen  22.94% no label  18.15% Caudate | R  R  n/a  R | 31.5 | 19.5 | 5.5 | 8.08 | 7184 |
|  | 58.00% Cerebelum Crus2  28.57% Cerebelum Crus1  13.43% no label | L  L  n/a | -28.5 | -86.5 | -40.5 | -7.13 | 4648 |
|  | 52.32% Putamen  27.99% Caudate  18.92% no label | L  L  n/a | -16.5 | 1.5 | 17.5 | 7.21 | 4144 |
|  | 52.88% Cerebelum Crus2  46.52% Cerebelum Crus1 | R  R | 21.5 | -84.5 | -32.5 | -6.94 | 4024 |
|  | 81.68% Insula  10.56% Frontal Inf Tri | L  L | -32.5 | 19.5 | 3.5 | 8.79 | 3712 |
|  | 34.11% ParaHippocampal  22.40% Lingual  19.01% no label  18.23% Fusiform | L  L  n/a  L | -34.5 | -36.5 | -12.5 | -7.69 | 3072 |
|  | 68.22% Frontal Sup 2  28.86% Frontal Mid 2 | R  R | 21.5 | 29.5 | 43.5 | -6.83 | 2744 |
|  | 89.20% Lingual  6.79% Cerebelum 6 | L  L | -12.5 | -76.5 | -6.5 | -7.73 | 2592 |
|  | 92.98% Lingual | R | 13.5 | -74.5 | -6.5 | -8.04 | 2280 |
|  | 61.73% Frontal Mid 2  38.27% Frontal Inf Tri | R  R | 49.5 | 35.5 | 25.5 | 6.82 | 1944 |
|  | 37.39% Fusiform  31.53% ParaHippocampal  17.12% Lingual  11.71% no label | R  R  R  n/a | 33.5 | -52.5 | -4.5 | -7.01 | 1776 |
|  | 68.27% Hippocampus  30.29% Amygdala | L  L | -20.5 | -2.5 | -20.5 | -6.57 | 1664 |
|  | 92.22% Cerebelum 9  7.78% no label | R  n/a | 9.5 | -46.5 | -52.5 | -6.8 | 720 |
|  | 67.11% Postcentral  31.58% Parietal Sup | R  R | 23.5 | -40.5 | 73.5 | -6.51 | 608 |
|  | 61.76% Postcentral  36.76% Parietal Sup | L  L | -28.5 | -42.5 | 69.5 | -6.49 | 544 |
|  | 47.69% Insula  46.15% Rolandic Oper  6.15% no label | R  R  n/a | 39.5 | -2.5 | 11.5 | 6.69 | 520 |
|  | 80.65% no label  16.13% Thalamus | n/a  R | 23.5 | -24.5 | -4.5 | 8.26 | 496 |
|  | 61.40% Rolandic Oper  29.82% Insula  8.77% no label | L  L  n/a | -40.5 | -6.5 | 13.5 | 6.74 | 456 |
|  | 58.82% no label  31.37% Hippocampus  9.80% Thalamus | n/a  L  L | -24.5 | -26.5 | -4.5 | 6.93 | 408 |
|  | 69.05% Cingulate Ant  23.81% Cingulate Mid  7.14% no label | L  L  n/a | -4.5 | 5.5 | 29.5 | 6.67 | 336 |
|  | 100.00% Thalamus | L | -10.5 | -18.5 | 5.5 | 7.03 | 320 |
|  | 96.55% Cerebelum 9 | L | -6.5 | -46.5 | -48.5 | -6.55 | 232 |
|  | 96.43% Rolandic Oper | R | 43.5 | -14.5 | 17.5 | -6.28 | 224 |
|  | 92.59% Cingulate Mid | R | 5.5 | 1.5 | 31.5 | 7.18 | 216 |
|  | 100.00% Cerebelum Crus2 | R | 47.5 | -54.5 | -42.5 | -6.23 | 216 |
|  | 80.00% Temporal Inf  20.00% no label | L  n/a | -52.5 | 5.5 | -42.5 | -6.03 | 120 |
|  | 71.43% Temporal Mid  28.57% no label | L  n/a | -68.5 | -48.5 | -0.5 | -6.05 | 112 |
|  | 85.71% Insula  7.14% Putamen  7.14% no label | R  R  n/a | 33.5 | 7.5 | 11.5 | -6.41 | 112 |
|  | 100.00% Precentral | L | -40.5 | -14.5 | 57.5 | 5.93 | 80 |

#### Supplementary Figure 10.3: Switch Conjunction Analysis, *P_fwe_ < 0.05.*

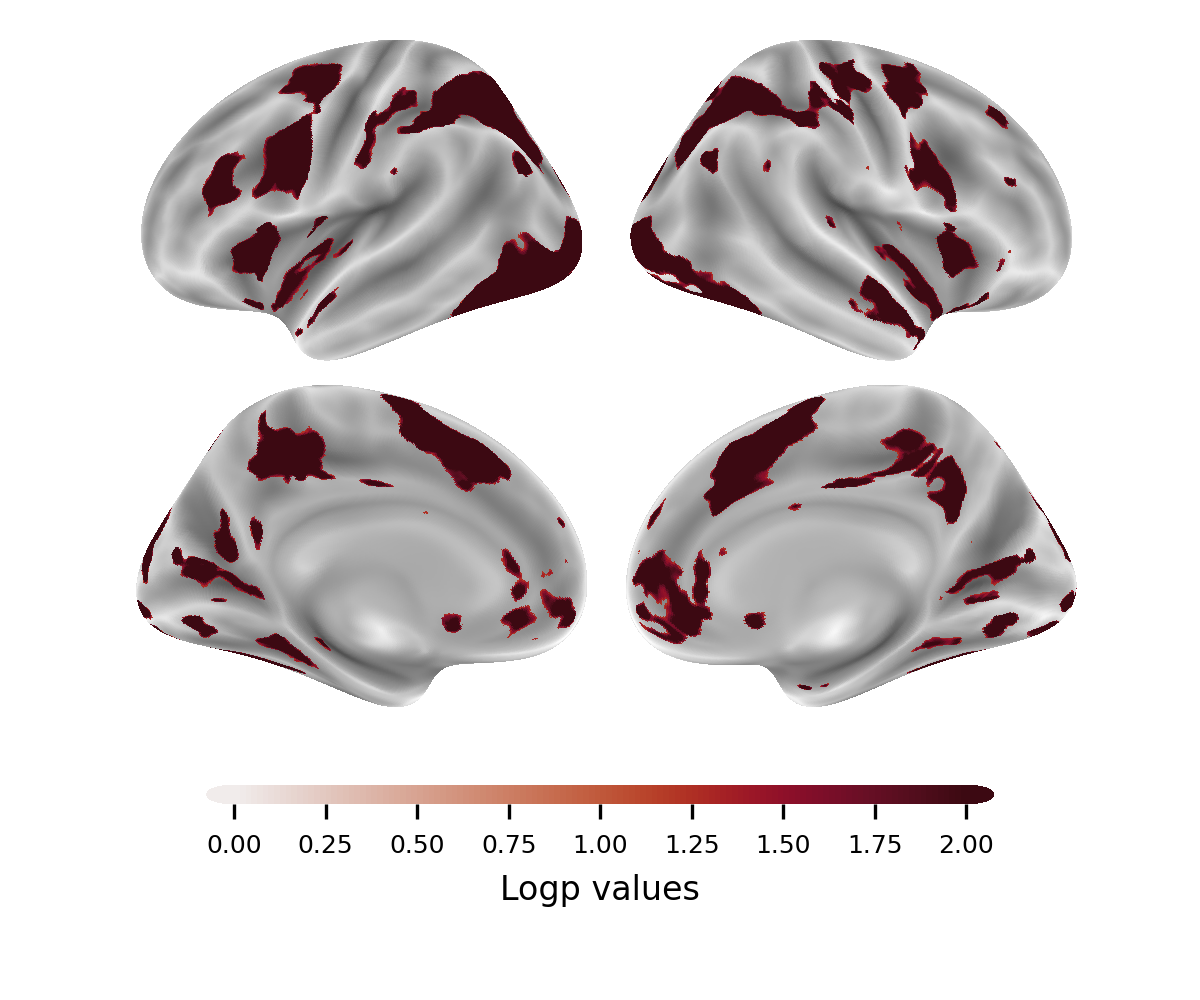

#### Supplementary Table 10.3: Switch Conjunction Analysis, *P_fwe_ < 0.05,* Cluster Table.

| Task condition | Anatomical label | Hemisphere | x | y | z | cluster mean log P value | mm3 |
| --- | --- | --- | --- | --- | --- | --- | --- |
| Switch-similiarity & Preference-switch | 16.13% Occipital Inf  13.93% Cerebelum 6  10.77% Fusiform  7.97% Temporal Inf  6.78% Occipital Mid  5.72% Cerebelum 8 | R  R  R  R  R  R | 23.5 | -76.5 | -18.5 | 2.54 | 43880 |
|  | 19.39% Occipital Mid  18.26% Occipital Inf  16.60% Fusiform  13.45% no label  9.35% Cerebelum 6  6.90% Cerebelum Crus1  6.78% Temporal Inf | L  L  L  n/a  L  L  L | -36.5 | -78.5 | -12.5 | 2.58 | 34208 |
|  | 40.84% Parietal Inf  25.73% Parietal Sup  10.65% Occipital Mid  7.60% no label  7.48% Postcentral | L  L  L  n/a  L | -34.5 | -56.5 | 45.5 | 2.56 | 28544 |
|  | 20.85% Parietal Sup  16.08% Parietal Inf  15.54% Postcentral  11.33% Precentral  9.10% no label  8.95% Occipital Mid  8.02% Angular | R  R  R  R  n/a  R  R | 33.5 | -50.5 | 47.5 | 2.52 | 26824 |
|  | 49.31% Precentral  18.28% Frontal Inf Tri  11.38% Frontal Inf Oper  8.77% Frontal Sup 2  6.25% Frontal Mid 2  5.71% no label | L  L  L  L  L  n/a | -44.5 | 5.5 | 37.5 | 2.54 | 20880 |
|  | 40.98% Supp Motor Area  32.93% Supp Motor Area  10.52% Cingulate Mid  6.93% Frontal Sup Medial | L  R  R  L | -0.5 | 7.5 | 51.5 | 2.58 | 18704 |
|  | 30.86% Precuneus  19.90% Precuneus  19.64% Cingulate Mid  17.19% Cingulate Mid  7.12% no label | R  L  L  R  n/a | 1.5 | -46.5 | 37.5 | 2.4 | 15400 |
|  | 32.84% Precentral  28.52% Frontal Inf Oper  16.70% Frontal Mid 2  13.66% Frontal Sup 2  6.79% no label | R  R  R  R  n/a | 41.5 | 1.5 | 41.5 | 2.51 | 11304 |
|  | 22.23% Temporal Mid  15.67% Temporal Sup  15.38% Temporal Pole Sup  14.94% Temporal Pole Mid  13.41% Insula  10.20% no label  6.92% OFCpost | R  R  R  R  R  n/a  R | 45.5 | -0.5 | -18.5 | 2.45 | 10976 |
|  | 24.55% Cingulate Ant  21.76% Cingulate Ant  20.44% Frontal Med Orb  16.94% Frontal Med Orb  10.33% Frontal Sup Medial | L  R  R  L  R | 1.5 | 41.5 | -2.5 | 2.22 | 10296 |
|  | 31.20% Cuneus  23.93% Cuneus  18.80% Occipital Sup  17.69% Occipital Sup | L  R  R  L | 1.5 | -92.5 | 23.5 | 2.52 | 9360 |
|  | 43.50% Calcarine  38.12% Calcarine  7.06% Lingual | L  R  R | -0.5 | -68.5 | 9.5 | 2.43 | 7136 |
|  | 30.01% Putamen  26.83% Insula  19.78% no label  17.84% Caudate | R  R  n/a  R | 25.5 | 11.5 | 5.5 | 2.5 | 5784 |
|  | 45.82% Precuneus  26.24% Cuneus  13.12% no label  10.27% Calcarine | L  L  n/a  L | -12.5 | -58.5 | 17.5 | 2.52 | 4208 |
|  | 60.68% Putamen  21.36% Caudate  17.05% no label | L  L  n/a | -22.5 | 1.5 | 9.5 | 2.43 | 3520 |
|  | 50.26% no label  28.57% Occipital Mid  18.37% Angular | n/a  L  L | -44.5 | -78.5 | 33.5 | 2.46 | 3136 |
|  | 85.20% Insula  10.20% Frontal Inf Tri | L  L | -34.5 | 17.5 | 3.5 | 2.56 | 3136 |
|  | 33.94% ParaHippocampal  23.55% Lingual  19.27% no label  18.04% Fusiform | L  L  n/a  L | -32.5 | -42.5 | -10.5 | 2.53 | 2616 |
|  | 47.84% Temporal Mid  26.54% Temporal Pole Sup  12.04% Temporal Sup  10.19% Temporal Pole Mid | L  L  L  L | -56.5 | 1.5 | -20.5 | 2.2 | 2592 |
|  | 89.24% Lingual  6.96% Cerebelum 6 | L  L | -14.5 | -78.5 | -8.5 | 2.5 | 2528 |
|  | 41.87% Temporal Sup  39.10% Insula  15.22% no label | L  L  n/a | -42.5 | -10.5 | -8.5 | 2.45 | 2312 |
|  | 95.06% Lingual | R | 13.5 | -74.5 | -6.5 | 2.55 | 2104 |
|  | 68.18% Angular  25.91% Occipital Mid  5.91% no label | R  R  n/a | 47.5 | -72.5 | 29.5 | 2.43 | 1760 |
|  | 58.17% Frontal Sup Medial  34.62% Frontal Sup Medial | L  R | -0.5 | 53.5 | 27.5 | 2.17 | 1664 |
|  | 38.54% Fusiform  28.65% ParaHippocampal  19.79% Lingual  13.02% no label | R  R  R  n/a | 29.5 | -48.5 | -6.5 | 2.38 | 1536 |
|  | 56.00% no label  33.33% Temporal Sup  10.67% SupraMarginal | n/a  L  L | -68.5 | -30.5 | 17.5 | 2.25 | 1200 |
|  | 78.90% Frontal Inf Orb 2  19.27% Frontal Inf Tri | R  R | 45.5 | 33.5 | -10.5 | 2.18 | 872 |
|  | 69.07% Frontal Sup 2  30.93% Frontal Mid 2 | R  R | 21.5 | 27.5 | 41.5 | 2.27 | 776 |
|  | 32.29% no label  25.00% Olfactory  23.96% Caudate  12.50% Olfactory  6.25% Caudate | n/a  R  L  L  R | -0.5 | 11.5 | -6.5 | 2.28 | 768 |
|  | 51.72% Vermis 9  25.29% Vermis 10  18.39% Cerebelum 9 | n/a  n/a  L | -0.5 | -52.5 | -34.5 | 2.51 | 696 |
|  | 50.79% Frontal Inf Tri  49.21% Frontal Mid 2 | R  R | 49.5 | 33.5 | 23.5 | 2.06 | 504 |
|  | 71.67% Insula  28.33% OFCpost | L  L | -30.5 | 15.5 | -20.5 | 2.36 | 480 |
|  | 94.74% Cerebelum 9  5.26% no label | R  n/a | 9.5 | -46.5 | -50.5 | 2.28 | 456 |
|  | 100.00% Temporal Sup | R | 51.5 | -26.5 | 9.5 | 2.39 | 424 |
|  | 87.76% Postcentral  12.24% SupraMarginal | R  R | 61.5 | -12.5 | 25.5 | 2.11 | 392 |
|  | 79.17% no label  18.75% Thalamus | n/a  R | 21.5 | -26.5 | -4.5 | 2.56 | 384 |
|  | 47.73% Hippocampus  27.27% Amygdala  22.73% ParaHippocampal | R  R  R | 25.5 | -4.5 | -28.5 | 2.25 | 352 |
|  | 71.43% Rolandic Oper  28.57% Insula | L  L | -40.5 | -6.5 | 13.5 | 2.41 | 336 |
|  | 97.62% Hippocampus | L | -32.5 | -14.5 | -20.5 | 2.08 | 336 |
|  | 58.54% no label  29.27% Hippocampus  12.20% Thalamus | n/a  L  L | -22.5 | -28.5 | -6.5 | 2.43 | 328 |
|  | 100.00% Thalamus | L | -12.5 | -18.5 | 3.5 | 2.47 | 320 |
|  | 51.52% SupraMarginal  48.48% Angular | R  R | 51.5 | -48.5 | 27.5 | 2.18 | 264 |
|  | 69.70% Cingulate Ant  21.21% Cingulate Mid  9.09% no label | L  L  n/a | -4.5 | 3.5 | 27.5 | 2.33 | 264 |
|  | 96.55% Cerebelum 9 | L | -8.5 | -48.5 | -48.5 | 2.28 | 232 |
|  | 100.00% Cingulate Mid | R | 5.5 | -0.5 | 29.5 | 2.35 | 192 |
|  | 69.57% Rolandic Oper  30.43% Insula | R  R | 39.5 | -4.5 | 11.5 | 2.09 | 184 |
|  | 53.33% Postcentral  46.67% Parietal Sup | L  L | -28.5 | -42.5 | 69.5 | 2.08 | 120 |
|  | 92.31% Precuneus  7.69% Postcentral | R  R | 5.5 | -52.5 | 71.5 | 2.05 | 104 |
|  | 100.00% Postcentral | R | 25.5 | -40.5 | 71.5 | 2.16 | 104 |
|  | 90.91% Cerebelum 4 5  9.09% Cerebelum 6 | L  L | -14.5 | -50.5 | -22.5 | 1.91 | 88 |
|  | 100.00% Precentral | L | -40.5 | -14.5 | 57.5 | 1.93 | 80 |

### Supplementary 11: Conjunction Analysis for Repeated Trials

#### Supplementary Figure 11.1: Interaction Model: Similarity Repeated condition, one sample two tailed t-test, *P_fwe_ < 0.05.*

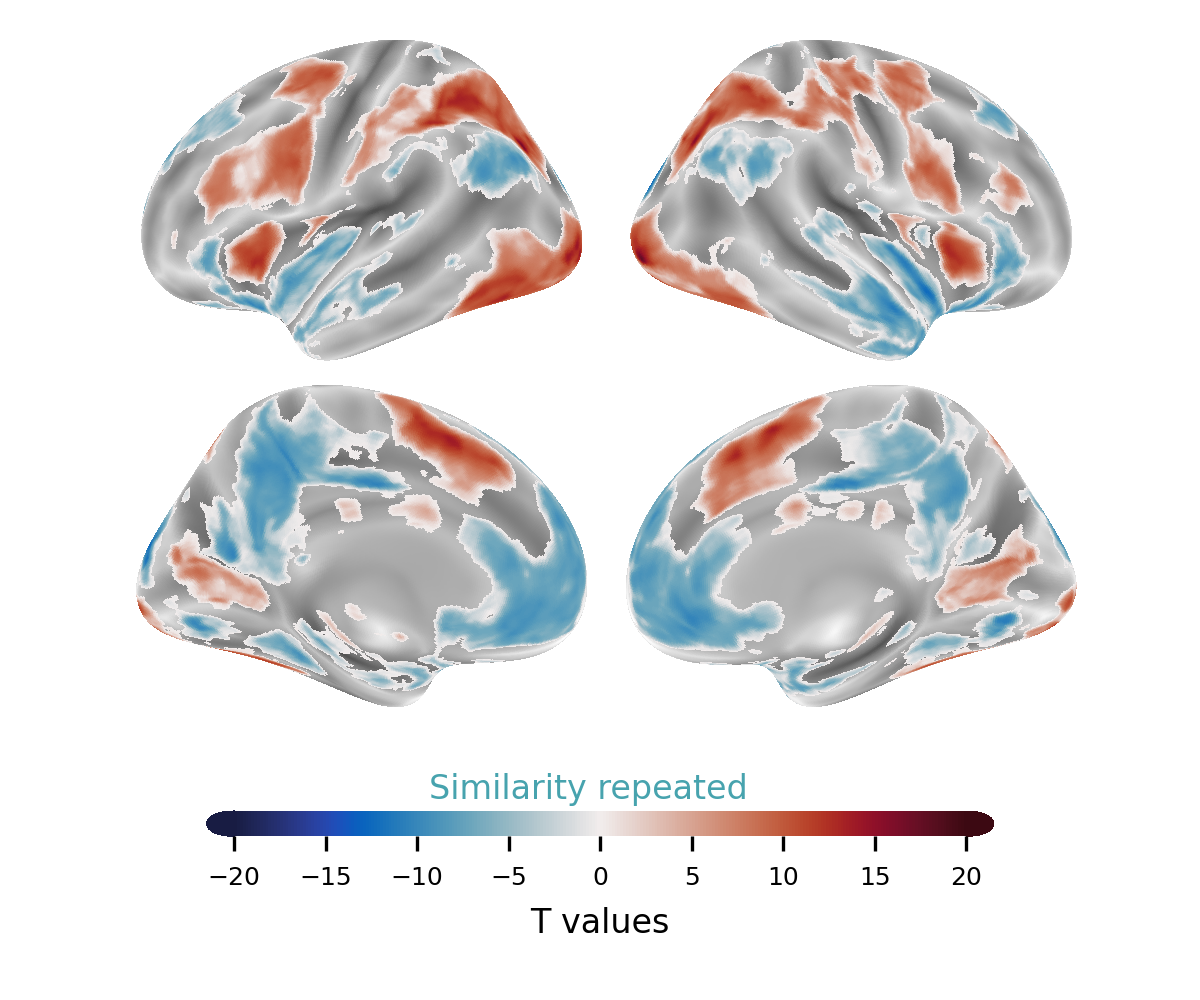

#### Supplementary Table 11.1: Interaction Model: Similarity Repeated condition, one sample two tailed t-test, *P_fwe_ < 0.05,* Cluster Table.

| Task condition | Anatomical label | Hemisphere | x | y | z | cluster mean | mm3 |
| --- | --- | --- | --- | --- | --- | --- | --- |
| Similarity repeated | 15.41% Occipital Inf  14.17% Cerebelum 6  10.25% Fusiform  7.89% Temporal Inf  6.53% Cerebelum Crus1  6.34% Occipital Mid  5.73% Cerebelum 8 | R  R  R  R  R  R  R | 29.5 | -94.5 | -8.5 | 8.23 | 47072 |
|  | 18.95% Occipital Mid  17.71% Occipital Inf  16.45% Fusiform  13.69% no label  9.08% Cerebelum 6  7.87% Cerebelum Crus1  6.88% Temporal Inf | L  L  L  n/a  L  L  L | -22.5 | -98.5 | -4.5 | 8.7 | 35592 |
|  | 37.72% Parietal Inf  23.00% Parietal Sup  11.05% Postcentral  9.31% Occipital Mid  8.03% no label | L  L  L  L  n/a | -46.5 | -42.5 | 49.5 | 8.4 | 32664 |
|  | 20.06% Parietal Sup  17.48% Postcentral  15.42% Parietal Inf  11.67% Precentral  8.78% no label  8.58% Occipital Mid  7.75% Angular | R  R  R  R  n/a  R  R | 33.5 | -68.5 | 31.5 | 7.88 | 27960 |
|  | 47.02% Precentral  19.23% Frontal Inf Tri  10.50% Frontal Inf Oper  8.29% Frontal Sup 2  8.11% Frontal Mid 2  6.21% no label | L  L  L  L  L  n/a | -28.5 | -6.5 | 53.5 | 7.87 | 23464 |
|  | 39.33% Supp Motor Area  31.49% Supp Motor Area  10.88% Cingulate Mid  7.23% Frontal Sup Medial | L  R  R  L | -0.5 | 17.5 | 45.5 | 8.37 | 21032 |
|  | 31.83% Precuneus  19.12% Precuneus  18.92% Cingulate Mid  16.63% Cingulate Mid  7.99% no label | R  L  L  R  n/a | -6.5 | -40.5 | 45.5 | -7.16 | 16112 |
|  | 34.28% Precentral  27.91% Frontal Inf Oper  16.34% Frontal Mid 2  13.37% Frontal Sup 2  6.65% no label | R  R  R  R  n/a | 33.5 | -2.5 | 57.5 | 7.57 | 11552 |
|  | 22.14% Temporal Mid  16.00% Temporal Sup  15.71% Temporal Pole Sup  14.64% Temporal Pole Mid  13.14% Insula  10.36% no label  6.79% OFCpost | R  R  R  R  R  n/a  R | 53.5 | -0.5 | -18.5 | -7.52 | 11200 |
|  | 29.74% Cuneus  23.70% Cuneus  19.42% Occipital Sup  17.35% Occipital Sup | L  R  R  L | -6.5 | -94.5 | 17.5 | -8.12 | 10464 |
|  | 24.48% Cingulate Ant  21.62% Cingulate Ant  20.54% Frontal Med Orb  16.91% Frontal Med Orb  10.27% Frontal Sup Medial | L  R  R  L  R | 7.5 | 35.5 | -4.5 | -6.48 | 10360 |
|  | 39.49% Putamen  31.04% Insula  9.68% Caudate  9.50% no label  6.10% Frontal Inf Tri | L  L  L  n/a  L | -34.5 | 17.5 | 3.5 | 7.75 | 9176 |
|  | 42.22% Calcarine  36.01% Calcarine  7.93% Lingual  5.70% Lingual | L  R  R  L | 11.5 | -62.5 | 7.5 | 7.04 | 7864 |
|  | 36.77% Putamen  22.53% Insula  17.38% no label  17.26% Caudate | R  R  n/a  R | 31.5 | 19.5 | 7.5 | 7.81 | 7136 |
|  | 40.43% Precuneus  26.68% Cuneus  16.53% no label  9.66% Calcarine | L  L  n/a  L | -12.5 | -56.5 | 13.5 | -8.34 | 4888 |
|  | 51.22% no label  31.45% Occipital Mid  15.25% Angular | n/a  L  L | -40.5 | -82.5 | 33.5 | -7.53 | 4248 |
|  | 46.83% Temporal Mid  29.75% Temporal Pole Sup  10.74% Temporal Sup  9.09% Temporal Pole Mid | L  L  L  L | -56.5 | 1.5 | -16.5 | -6.52 | 2904 |
|  | 88.12% Lingual  6.35% Cerebelum 6 | L  L | -12.5 | -80.5 | -4.5 | -7.77 | 2896 |
|  | 32.10% ParaHippocampal  22.73% Lingual  20.17% no label  18.18% Fusiform | L  L  n/a  L | -28.5 | -40.5 | -6.5 | -7.85 | 2816 |
|  | 53.44% Angular  39.06% Occipital Mid | R  R | 49.5 | -72.5 | 25.5 | -7.57 | 2560 |
|  | 41.36% Temporal Sup  38.98% Insula  15.59% no label | L  L  n/a | -40.5 | -6.5 | -6.5 | -7.18 | 2360 |
|  | 94.96% Lingual | R | 13.5 | -76.5 | -4.5 | -7.99 | 2224 |
|  | 36.07% Fusiform  26.48% ParaHippocampal  21.00% Lingual  16.44% no label | R  R  R  n/a | 31.5 | -52.5 | -2.5 | -6.84 | 1752 |
|  | 58.17% Frontal Sup Medial  34.62% Frontal Sup Medial | L  R | 1.5 | 53.5 | 27.5 | -6.37 | 1664 |
|  | 100.00% Thalamus | L | -12.5 | -18.5 | 5.5 | 8.09 | 1272 |
|  | 56.49% no label  33.12% Temporal Sup  10.39% SupraMarginal | n/a  L  L | -68.5 | -28.5 | 15.5 | -6.57 | 1232 |
|  | 50.79% no label  23.02% Cingulate Ant  19.05% Cingulate Mid  5.56% Cingulate Mid | n/a  L  L  R | -4.5 | 5.5 | 29.5 | 6.62 | 1008 |
|  | 36.70% no label  22.94% Olfactory  21.10% Caudate  11.01% Olfactory  8.26% Caudate | n/a  R  L  L  R | -2.5 | 11.5 | -4.5 | -6.48 | 872 |
|  | 78.90% Frontal Inf Orb 2  19.27% Frontal Inf Tri | R  R | 45.5 | 35.5 | -8.5 | -6.38 | 872 |
|  | 100.00% Thalamus | R | 11.5 | -18.5 | 9.5 | 7.18 | 832 |
|  | 69.07% Frontal Sup 2  30.93% Frontal Mid 2 | R  R | 23.5 | 29.5 | 43.5 | -6.48 | 776 |
|  | 51.72% Vermis 9  25.29% Vermis 10  18.39% Cerebelum 9 | n/a  n/a  L | -2.5 | -50.5 | -34.5 | 7.4 | 696 |
|  | 90.36% Cerebelum 9  9.64% no label | R  n/a | 11.5 | -46.5 | -52.5 | -6.56 | 664 |
|  | 70.51% Frontal Inf Tri  29.49% Frontal Mid 2 | L  L | -48.5 | 45.5 | 5.5 | 6.24 | 624 |
|  | 53.97% no label  26.98% Hippocampus  19.05% Thalamus | n/a  L  L | -18.5 | -30.5 | -2.5 | 6.96 | 504 |
|  | 50.79% Frontal Inf Tri  49.21% Frontal Mid 2 | R  R | 49.5 | 33.5 | 23.5 | 6.19 | 504 |
|  | 93.55% Cerebelum 9  6.45% no label | L  n/a | -8.5 | -46.5 | -46.5 | -6.74 | 496 |
|  | 71.67% Insula  28.33% OFCpost | L  L | -28.5 | 17.5 | -18.5 | -6.84 | 480 |
|  | 71.93% no label  22.81% Thalamus  5.26% Hippocampus | n/a  R  R | 23.5 | -26.5 | -2.5 | 7.79 | 456 |
|  | 98.21% Temporal Sup | R | 55.5 | -24.5 | 11.5 | -6.78 | 448 |
|  | 67.31% Rolandic Oper  32.69% Insula | L  L | -40.5 | -6.5 | 13.5 | 6.74 | 416 |
|  | 97.92% Hippocampus | L | -32.5 | -14.5 | -18.5 | -6.26 | 384 |
|  | 46.67% Hippocampus  26.67% Amygdala  24.44% ParaHippocampal | R  R  R | 27.5 | -4.5 | -26.5 | -6.48 | 360 |
|  | 51.52% SupraMarginal  48.48% Angular | R  R | 53.5 | -48.5 | 27.5 | -6.37 | 264 |
|  | 56.25% Paracentral Lobule  43.75% Precuneus | L  L | -6.5 | -40.5 | 71.5 | 6.46 | 256 |
|  | 82.14% Temporal Mid  17.86% Temporal Sup | L  L | -50.5 | -44.5 | 9.5 | 6.3 | 224 |
|  | 100.00% Frontal Mid 2 | L | -36.5 | 55.5 | 11.5 | 6.12 | 200 |
|  | 100.00% Cerebelum 8 | R | 35.5 | -48.5 | -50.5 | 6.1 | 200 |
|  | 100.00% Cingulate Mid | R | 5.5 | 1.5 | 31.5 | 6.78 | 192 |
|  | 69.57% Rolandic Oper  30.43% Insula | R  R | 39.5 | -4.5 | 13.5 | 6.2 | 184 |
|  | 100.00% Postcentral | R | 27.5 | -40.5 | 71.5 | -6.22 | 128 |
|  | 53.33% Postcentral  46.67% Parietal Sup | L  L | -28.5 | -42.5 | 69.5 | -6.21 | 120 |
|  | 92.31% Precuneus  7.69% Postcentral | R  R | 5.5 | -52.5 | 71.5 | -6.13 | 104 |
|  | 91.67% Cerebelum 4 5  8.33% Cerebelum 6 | L  L | -14.5 | -50.5 | -22.5 | 6.1 | 96 |

#### Supplementary Figure 11.2: Interaction Model: Preference Repeated condition, one sample two tailed t-test, *P_fwe_ < 0.05.*

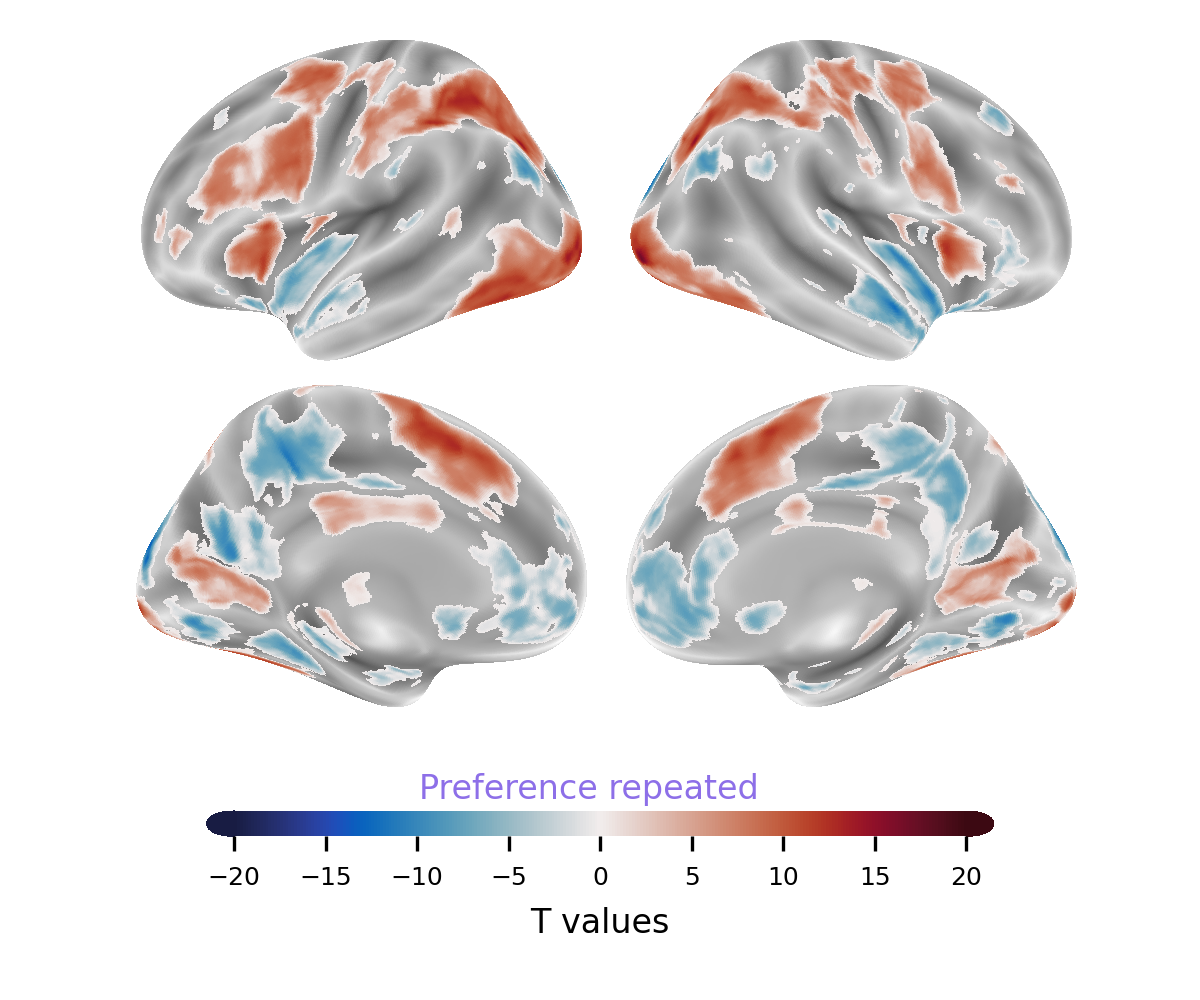

#### Supplementary Table 11.2: Interaction Model: Preference Repeated condition, one sample two tailed t-test, *P_fwe_ < 0.05*, Cluster Table.

| Task condition | Anatomical label | Hemisphere | x | y | z | cluster mean T value | mm3 |
| --- | --- | --- | --- | --- | --- | --- | --- |
| Preference repeated | 10.46% Frontal Sup Medial  7.84% Frontal Sup Medial  7.32% no label  6.13% Cingulate Ant  5.94% Frontal Sup 2  5.61% Temporal Mid | L  R  n/a  L  L  R | 41.5 | -6.5 | -12.5 | -7.12 | 107808 |
|  | 7.89% Occipital Inf  7.70% Occipital Mid  7.45% Cerebelum 6  7.02% Cerebelum 6  6.96% Occipital Inf  6.91% no label  6.40% Fusiform  5.73% Fusiform | R  L  R  L  L  n/a  L  R | 29.5 | -94.5 | -8.5 | 8.38 | 90816 |
|  | 16.23% Precentral  14.66% Parietal Sup  14.11% Postcentral  10.32% Parietal Inf  8.61% no label  7.90% Frontal Inf Oper  6.05% Occipital Mid | R  R  R  R  n/a  R  R | 31.5 | -68.5 | 31.5 | 8.01 | 45880 |
|  | 24.52% Precuneus  20.41% Precuneus  17.33% Cingulate Mid  14.18% Cingulate Mid  8.62% no label | L  R  L  R  n/a | -12.5 | -56.5 | 13.5 | -7.47 | 33512 |
|  | 38.72% Parietal Inf  25.39% Parietal Sup  11.49% Occipital Mid  8.00% Postcentral  7.97% no label | L  L  L  L  n/a | -28.5 | -70.5 | 29.5 | 8.62 | 30496 |
|  | 48.42% Precentral  18.39% Frontal Inf Tri  11.76% Frontal Inf Oper  8.86% Frontal Sup 2  6.41% Frontal Mid 2  5.75% no label | L  L  L  L  L  n/a | -30.5 | -4.5 | 57.5 | 7.95 | 21840 |
|  | 39.65% Supp Motor Area  32.30% Supp Motor Area  10.21% Cingulate Mid  7.47% Frontal Sup Medial | L  R  R  L | -8.5 | 11.5 | 51.5 | 8.57 | 19592 |
|  | 43.30% no label  34.32% Angular  7.23% Occipital Mid  6.99% Temporal Sup | n/a  L  L  L | -48.5 | -72.5 | 37.5 | -7.32 | 13728 |
|  | 30.65% Cuneus  23.79% Cuneus  18.95% Occipital Sup  16.77% Occipital Sup | L  R  R  L | -6.5 | -94.5 | 17.5 | -7.96 | 9920 |
|  | 62.12% Angular  13.95% SupraMarginal  7.32% Parietal Inf  6.03% Occipital Mid | R  R  R  R | 49.5 | -70.5 | 41.5 | -6.84 | 8088 |
|  | 40.96% Calcarine  39.74% Calcarine  6.81% Lingual | L  R  R | 13.5 | -64.5 | 9.5 | 7.1 | 7872 |
|  | 27.73% Insula  25.06% Putamen  22.94% no label  18.15% Caudate | R  R  n/a  R | 31.5 | 19.5 | 5.5 | 8.08 | 7184 |
|  | 58.00% Cerebelum Crus2  28.57% Cerebelum Crus1  13.43% no label | L  L  n/a | -28.5 | -86.5 | -40.5 | -7.13 | 4648 |
|  | 52.32% Putamen  27.99% Caudate  18.92% no label | L  L  n/a | -16.5 | 1.5 | 17.5 | 7.21 | 4144 |
|  | 52.88% Cerebelum Crus2  46.52% Cerebelum Crus1 | R  R | 21.5 | -84.5 | -32.5 | -6.94 | 4024 |
|  | 81.68% Insula  10.56% Frontal Inf Tri | L  L | -32.5 | 19.5 | 3.5 | 8.79 | 3712 |
|  | 34.11% ParaHippocampal  22.40% Lingual  19.01% no label  18.23% Fusiform | L  L  n/a  L | -34.5 | -36.5 | -12.5 | -7.69 | 3072 |
|  | 68.22% Frontal Sup 2  28.86% Frontal Mid 2 | R  R | 21.5 | 29.5 | 43.5 | -6.83 | 2744 |
|  | 89.20% Lingual  6.79% Cerebelum 6 | L  L | -12.5 | -76.5 | -6.5 | -7.73 | 2592 |
|  | 92.98% Lingual | R | 13.5 | -74.5 | -6.5 | -8.04 | 2280 |
|  | 61.73% Frontal Mid 2  38.27% Frontal Inf Tri | R  R | 49.5 | 35.5 | 25.5 | 6.82 | 1944 |
|  | 37.39% Fusiform  31.53% ParaHippocampal  17.12% Lingual  11.71% no label | R  R  R  n/a | 33.5 | -52.5 | -4.5 | -7.01 | 1776 |
|  | 68.27% Hippocampus  30.29% Amygdala | L  L | -20.5 | -2.5 | -20.5 | -6.57 | 1664 |
|  | 92.22% Cerebelum 9  7.78% no label | R  n/a | 9.5 | -46.5 | -52.5 | -6.8 | 720 |
|  | 67.11% Postcentral  31.58% Parietal Sup | R  R | 23.5 | -40.5 | 73.5 | -6.51 | 608 |
|  | 61.76% Postcentral  36.76% Parietal Sup | L  L | -28.5 | -42.5 | 69.5 | -6.49 | 544 |
|  | 47.69% Insula  46.15% Rolandic Oper  6.15% no label | R  R  n/a | 39.5 | -2.5 | 11.5 | 6.69 | 520 |
|  | 80.65% no label  16.13% Thalamus | n/a  R | 23.5 | -24.5 | -4.5 | 8.26 | 496 |
|  | 61.40% Rolandic Oper  29.82% Insula  8.77% no label | L  L  n/a | -40.5 | -6.5 | 13.5 | 6.74 | 456 |
|  | 58.82% no label  31.37% Hippocampus  9.80% Thalamus | n/a  L  L | -24.5 | -26.5 | -4.5 | 6.93 | 408 |
|  | 69.05% Cingulate Ant  23.81% Cingulate Mid  7.14% no label | L  L  n/a | -4.5 | 5.5 | 29.5 | 6.67 | 336 |
|  | 100.00% Thalamus | L | -10.5 | -18.5 | 5.5 | 7.03 | 320 |
|  | 96.55% Cerebelum 9 | L | -6.5 | -46.5 | -48.5 | -6.55 | 232 |
|  | 96.43% Rolandic Oper | R | 43.5 | -14.5 | 17.5 | -6.28 | 224 |
|  | 92.59% Cingulate Mid | R | 5.5 | 1.5 | 31.5 | 7.18 | 216 |
|  | 100.00% Cerebelum Crus2 | R | 47.5 | -54.5 | -42.5 | -6.23 | 216 |
|  | 80.00% Temporal Inf  20.00% no label | L  n/a | -52.5 | 5.5 | -42.5 | -6.03 | 120 |
|  | 71.43% Temporal Mid  28.57% no label | L  n/a | -68.5 | -48.5 | -0.5 | -6.05 | 112 |
|  | 85.71% Insula  7.14% Putamen  7.14% no label | R  R  n/a | 33.5 | 7.5 | 11.5 | -6.41 | 112 |
|  | 100.00% Precentral | L | -40.5 | -14.5 | 57.5 | 5.93 | 80 |

#### Supplementary Figure 11.3: Repeated Conjunction Analysis, *P_fwe_ < 0.05*.

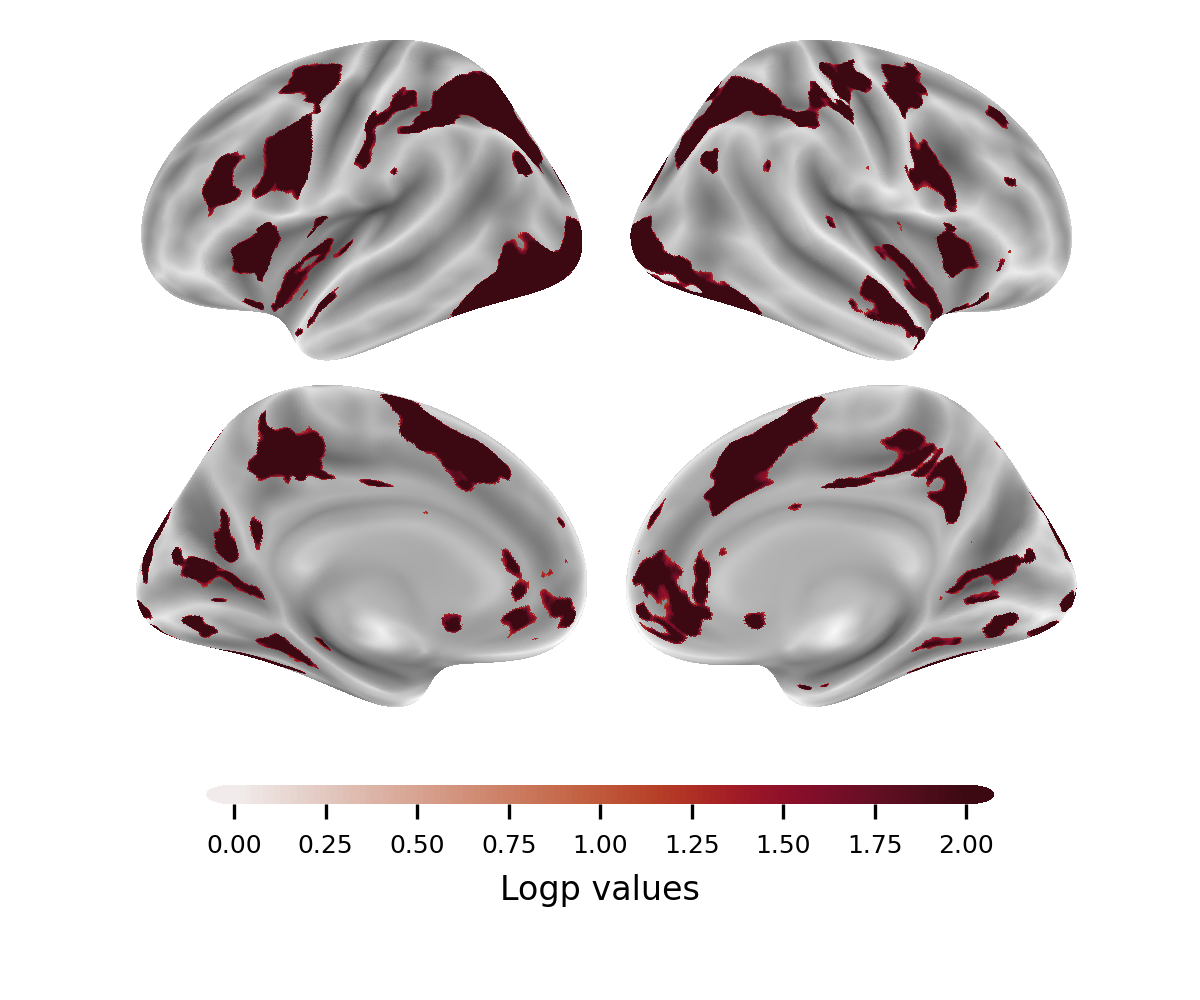

#### Supplementary Table 11.3: Repeated Conjunction Analysis, *P_fwe_ < 0.05*, Cluster Table.

| Task condition | Anatomical label | Hemisphere | x | y | z | cluster mean T value | mm3 |
| --- | --- | --- | --- | --- | --- | --- | --- |
| Similarity repeated & Preference repeated | 16.13% Occipital Inf  13.93% Cerebelum 6  10.77% Fusiform  7.97% Temporal Inf  6.78% Occipital Mid  5.72% Cerebelum 8 | R  R  R  R  R  R | 23.5 | -76.5 | -18.5 | 2.54 | 43880 |
|  | 19.39% Occipital Mid  18.26% Occipital Inf  16.60% Fusiform  13.45% no label  9.35% Cerebelum 6  6.90% Cerebelum Crus1  6.78% Temporal Inf | L  L  L  n/a  L  L  L | -36.5 | -78.5 | -12.5 | 2.58 | 34208 |
|  | 40.84% Parietal Inf  25.73% Parietal Sup  10.65% Occipital Mid  7.60% no label  7.48% Postcentral | L  L  L  n/a  L | -34.5 | -56.5 | 45.5 | 2.56 | 28544 |
|  | 20.85% Parietal Sup  16.08% Parietal Inf  15.54% Postcentral  11.33% Precentral  9.10% no label  8.95% Occipital Mid  8.02% Angular | R  R  R  R  n/a  R  R | 33.5 | -50.5 | 47.5 | 2.52 | 26824 |
|  | 49.31% Precentral  18.28% Frontal Inf Tri  11.38% Frontal Inf Oper  8.77% Frontal Sup 2  6.25% Frontal Mid 2  5.71% no label | L  L  L  L  L  n/a | -44.5 | 5.5 | 37.5 | 2.54 | 20880 |
|  | 40.98% Supp Motor Area  32.93% Supp Motor Area  10.52% Cingulate Mid  6.93% Frontal Sup Medial | L  R  R  L | -0.5 | 7.5 | 51.5 | 2.58 | 18704 |
|  | 30.86% Precuneus  19.90% Precuneus  19.64% Cingulate Mid  17.19% Cingulate Mid  7.12% no label | R  L  L  R  n/a | 1.5 | -46.5 | 37.5 | 2.4 | 15400 |
|  | 32.84% Precentral  28.52% Frontal Inf Oper  16.70% Frontal Mid 2  13.66% Frontal Sup 2  6.79% no label | R  R  R  R  n/a | 41.5 | 1.5 | 41.5 | 2.51 | 11304 |
|  | 22.23% Temporal Mid  15.67% Temporal Sup  15.38% Temporal Pole Sup  14.94% Temporal Pole Mid  13.41% Insula  10.20% no label  6.92% OFCpost | R  R  R  R  R  n/a  R | 45.5 | -0.5 | -18.5 | 2.45 | 10976 |
|  | 24.55% Cingulate Ant  21.76% Cingulate Ant  20.44% Frontal Med Orb  16.94% Frontal Med Orb  10.33% Frontal Sup Medial | L  R  R  L  R | 1.5 | 41.5 | -2.5 | 2.22 | 10296 |
|  | 31.20% Cuneus  23.93% Cuneus  18.80% Occipital Sup  17.69% Occipital Sup | L  R  R  L | 1.5 | -92.5 | 23.5 | 2.52 | 9360 |
|  | 43.50% Calcarine  38.12% Calcarine  7.06% Lingual | L  R  R | -0.5 | -68.5 | 9.5 | 2.43 | 7136 |
|  | 30.01% Putamen  26.83% Insula  19.78% no label  17.84% Caudate | R  R  n/a  R | 25.5 | 11.5 | 5.5 | 2.5 | 5784 |
|  | 45.82% Precuneus  26.24% Cuneus  13.12% no label  10.27% Calcarine | L  L  n/a  L | -12.5 | -58.5 | 17.5 | 2.52 | 4208 |
|  | 60.68% Putamen  21.36% Caudate  17.05% no label | L  L  n/a | -22.5 | 1.5 | 9.5 | 2.43 | 3520 |
|  | 50.26% no label  28.57% Occipital Mid  18.37% Angular | n/a  L  L | -44.5 | -78.5 | 33.5 | 2.46 | 3136 |
|  | 85.20% Insula  10.20% Frontal Inf Tri | L  L | -34.5 | 17.5 | 3.5 | 2.56 | 3136 |
|  | 33.94% ParaHippocampal  23.55% Lingual  19.27% no label  18.04% Fusiform | L  L  n/a  L | -32.5 | -42.5 | -10.5 | 2.53 | 2616 |
|  | 47.84% Temporal Mid  26.54% Temporal Pole Sup  12.04% Temporal Sup  10.19% Temporal Pole Mid | L  L  L  L | -56.5 | 1.5 | -20.5 | 2.2 | 2592 |
|  | 89.24% Lingual  6.96% Cerebelum 6 | L  L | -14.5 | -78.5 | -8.5 | 2.5 | 2528 |
|  | 41.87% Temporal Sup  39.10% Insula  15.22% no label | L  L  n/a | -42.5 | -10.5 | -8.5 | 2.45 | 2312 |
|  | 95.06% Lingual | R | 13.5 | -74.5 | -6.5 | 2.55 | 2104 |
|  | 68.18% Angular  25.91% Occipital Mid  5.91% no label | R  R  n/a | 47.5 | -72.5 | 29.5 | 2.43 | 1760 |
|  | 58.17% Frontal Sup Medial  34.62% Frontal Sup Medial | L  R | -0.5 | 53.5 | 27.5 | 2.17 | 1664 |
|  | 38.54% Fusiform  28.65% ParaHippocampal  19.79% Lingual  13.02% no label | R  R  R  n/a | 29.5 | -48.5 | -6.5 | 2.38 | 1536 |
|  | 56.00% no label  33.33% Temporal Sup  10.67% SupraMarginal | n/a  L  L | -68.5 | -30.5 | 17.5 | 2.25 | 1200 |
|  | 78.90% Frontal Inf Orb 2  19.27% Frontal Inf Tri | R  R | 45.5 | 33.5 | -10.5 | 2.18 | 872 |
|  | 69.07% Frontal Sup 2  30.93% Frontal Mid 2 | R  R | 21.5 | 27.5 | 41.5 | 2.27 | 776 |
|  | 32.29% no label  25.00% Olfactory  23.96% Caudate  12.50% Olfactory  6.25% Caudate | n/a  R  L  L  R | -0.5 | 11.5 | -6.5 | 2.28 | 768 |
|  | 51.72% Vermis 9  25.29% Vermis 10  18.39% Cerebelum 9 | n/a  n/a  L | -0.5 | -52.5 | -34.5 | 2.51 | 696 |
|  | 50.79% Frontal Inf Tri  49.21% Frontal Mid 2 | R  R | 49.5 | 33.5 | 23.5 | 2.06 | 504 |
|  | 71.67% Insula  28.33% OFCpost | L  L | -30.5 | 15.5 | -20.5 | 2.36 | 480 |
|  | 94.74% Cerebelum 9  5.26% no label | R  n/a | 9.5 | -46.5 | -50.5 | 2.28 | 456 |
|  | 100.00% Temporal Sup | R | 51.5 | -26.5 | 9.5 | 2.39 | 424 |
|  | 87.76% Postcentral  12.24% SupraMarginal | R  R | 61.5 | -12.5 | 25.5 | 2.11 | 392 |
|  | 79.17% no label  18.75% Thalamus | n/a  R | 21.5 | -26.5 | -4.5 | 2.56 | 384 |
|  | 47.73% Hippocampus  27.27% Amygdala  22.73% ParaHippocampal | R  R  R | 25.5 | -4.5 | -28.5 | 2.25 | 352 |
|  | 71.43% Rolandic Oper  28.57% Insula | L  L | -40.5 | -6.5 | 13.5 | 2.41 | 336 |
|  | 97.62% Hippocampus | L | -32.5 | -14.5 | -20.5 | 2.08 | 336 |
|  | 58.54% no label  29.27% Hippocampus  12.20% Thalamus | n/a  L  L | -22.5 | -28.5 | -6.5 | 2.43 | 328 |
|  | 100.00% Thalamus | L | -12.5 | -18.5 | 3.5 | 2.47 | 320 |
|  | 51.52% SupraMarginal  48.48% Angular | R  R | 51.5 | -48.5 | 27.5 | 2.18 | 264 |
|  | 69.70% Cingulate Ant  21.21% Cingulate Mid  9.09% no label | L  L  n/a | -4.5 | 3.5 | 27.5 | 2.33 | 264 |
|  | 96.55% Cerebelum 9 | L | -8.5 | -48.5 | -48.5 | 2.28 | 232 |
|  | 100.00% Cingulate Mid | R | 5.5 | -0.5 | 29.5 | 2.35 | 192 |
|  | 69.57% Rolandic Oper  30.43% Insula | R  R | 39.5 | -4.5 | 11.5 | 2.09 | 184 |
|  | 53.33% Postcentral  46.67% Parietal Sup | L  L | -28.5 | -42.5 | 69.5 | 2.08 | 120 |
|  | 92.31% Precuneus  7.69% Postcentral | R  R | 5.5 | -52.5 | 71.5 | 2.05 | 104 |
|  | 100.00% Postcentral | R | 25.5 | -40.5 | 71.5 | 2.16 | 104 |
|  | 90.91% Cerebelum 4 5  9.09% Cerebelum 6 | L  L | -14.5 | -50.5 | -22.5 | 1.91 | 88 |
|  | 100.00% Precentral | L | -40.5 | -14.5 | 57.5 | 1.93 | 80 |

### Supplementary 12: fMRIprep output prompt

fMRIprep workflow

Results included in this manuscript come from preprocessing performed using *fMRIPrep* 22.1.0+0.gce344b39.dirty (@fmriprep1; @fmriprep2; RRID:SCR_016216), which is based on *Nipype* 1.8.5 (@nipype1; @nipype2; RRID:SCR_002502).

Preprocessing of B0 inhomogeneity mappings

A total of 1 fieldmaps were found available within the input BIDS structure for this particular subject. A *B0* nonuniformity map (or *fieldmap*) was estimated from the phase-drift map(s) measure with two consecutive GRE (gradient-recalled echo) acquisitions. The corresponding phase-map(s) were phase-unwrapped with prelude (FSL 6.0.5.1:57b01774).

Anatomical data preprocessing

A total of 1 T1-weighted (T1w) images were found within the input BIDS dataset.The T1-weighted (T1w) image was corrected for intensity non-uniformity (INU) with N4BiasFieldCorrection [@n4], distributed with ANTs 2.3.3 [@ants, RRID:SCR_004757], and used as T1w-reference throughout the workflow. The T1w-reference was then skull-stripped with a *Nipype* implementation of the antsBrainExtraction.sh workflow (from ANTs), using OASIS30ANTs as target template. Brain tissue segmentation of cerebrospinal fluid (CSF), white-matter (WM) and gray-matter (GM) was performed on the brain-extracted T1w using fast [FSL 6.0.5.1:57b01774, RRID:SCR_002823, @fsl_fast]. Brain surfaces were reconstructed using recon-all [FreeSurfer 7.2.0, RRID:SCR_001847, @fs_reconall], and the brain mask estimated previously was refined with a custom variation of the method to reconcile ANTs-derived and FreeSurfer-derived segmentations of the cortical gray-matter of Mindboggle [RRID:SCR_002438, @mindboggle]. Volume-based spatial normalization to two standard spaces (MNI152NLin2009cAsym, MNI152NLin6Asym) was performed through nonlinear registration with antsRegistration (ANTs 2.3.3), using brain-extracted versions of both T1w reference and the T1w template. The following templates were selected for spatial normalization: *ICBM 152 Nonlinear Asymmetrical template version 2009c* [@mni152nlin2009casym, RRID:SCR_008796; TemplateFlow ID: MNI152NLin2009cAsym], *FSL's MNI ICBM 152 non-linear 6th Generation Asymmetric Average Brain Stereotaxic Registration Model* [@mni152nlin6asym, RRID:SCR_002823; TemplateFlow ID: MNI152NLin6Asym].

Functional data preprocessing

For each of the 4 BOLD runs found per subject (across all tasks and sessions), the following preprocessing was performed. First, a reference volume and its skull-stripped version were generated using a custom methodology of *fMRIPrep*. Head-motion parameters with respect to the BOLD reference (transformation matrices, and six corresponding rotation and translation parameters) are estimated before any spatiotemporal filtering using mcflirt [FSL 6.0.5.1:57b01774, @mcflirt]. BOLD runs were slice-time corrected to 0.964s (0.5 of slice acquisition range 0s-1.93s) using 3dTshift from AFNI [@afni, RRID:SCR_005927]. The BOLD time-series (including slice-timing correction when applied) were resampled onto their original, native space by applying the transforms to correct for head-motion. These resampled BOLD time-series will be referred to as *preprocessed BOLD in original space*, or just *preprocessed BOLD*. The BOLD reference was then co-registered to the T1w reference using bbregister (FreeSurfer) which implements boundary-based registration [@bbr]. Co-registration was configured with six degrees of freedom. Several confounding time-series were calculated based on the *preprocessed BOLD*: framewise displacement (FD), DVARS and three region-wise global signals. FD was computed using two formulations following Power (absolute sum of relative motions, @power_fd_dvars) and Jenkinson (relative root mean square displacement between affines, @mcflirt). FD and DVARS are calculated for each functional run, both using their implementations in *Nipype* [following the definitions by @power_fd_dvars]. The three global signals are extracted within the CSF, the WM, and the whole-brain masks. Additionally, a set of physiological regressors were extracted to allow for component-based noise correction [*CompCor*, @compcor]. Principal components are estimated after high-pass filtering the *preprocessed BOLD* time-series (using a discrete cosine filter with 128s cut-off) for the two *CompCor* variants: temporal (tCompCor) and anatomical (aCompCor). tCompCor components are then calculated from the top 2% variable voxels within the brain mask. For aCompCor, three probabilistic masks (CSF, WM and combined CSF+WM) are generated in anatomical space. The implementation differs from that of Behzadi et al. in that instead of eroding the masks by 2 pixels on BOLD space, a mask of pixels that likely contain a volume fraction of GM is subtracted from the aCompCor masks. This mask is obtained by dilating a GM mask extracted from the FreeSurfer's *aseg* segmentation, and it ensures components are not extracted from voxels containing a minimal fraction of GM. Finally, these masks are resampled into BOLD space and binarized by thresholding at 0.99 (as in the original implementation). Components are also calculated separately within the WM and CSF masks. For each CompCor decomposition, the *k* components with the largest singular values are retained, such that the retained components' time series are sufficient to explain 50 percent of variance across the nuisance mask (CSF, WM, combined, or temporal). The remaining components are dropped from consideration. The head-motion estimates calculated in the correction step were also placed within the corresponding confounds file. The confound time series derived from head motion estimates and global signals were expanded with the inclusion of temporal derivatives and quadratic terms for each [@confounds_satterthwaite_2013]. Frames that exceeded a threshold of 0.5 mm FD or 1.5 standardized DVARS were annotated as motion outliers. Additional nuisance time series are calculated by means of principal components analysis of the signal found within a thin band (*crown*) of voxels around the edge of the brain, as proposed by [@patriat_improved_2017]. The BOLD time-series were resampled into standard space, generating a *preprocessed BOLD run in MNI152NLin2009cAsym space*. First, a reference volume and its skull-stripped version were generated using a custom methodology of *fMRIPrep*. Automatic removal of motion artifacts using independent component analysis [ICA-AROMA, @aroma] was performed on the *preprocessed BOLD on MNI space* time-series after removal of non-steady state volumes and spatial smoothing with an isotropic, Gaussian kernel of 6mm FWHM (full-width half-maximum). Corresponding "non-aggressively" denoised runs were produced after such smoothing. Additionally, the "aggressive" noise-regressors were collected and placed in the corresponding confounds file. All resamplings can be performed with *a single interpolation step* by composing all the pertinent transformations (i.e. head-motion transform matrices, susceptibility distortion correction when available, and co-registrations to anatomical and output spaces). Gridded (volumetric) resamplings were performed using antsApplyTransforms (ANTs), configured with Lanczos interpolation to minimize the smoothing effects of other kernels [@lanczos]. Non-gridded (surface) resamplings were performed using mri_vol2surf (FreeSurfer).

Many internal operations of *fMRIPrep* use *Nilearn* 0.9.1 [@nilearn, RRID:SCR_001362], mostly within the functional processing workflow. For more details of the pipeline, see [the section corresponding to workflows in *fMRIPrep*'s documentation](https://fmriprep.readthedocs.io/en/latest/workflows.html).

Copyright Waiver

The above boilerplate text was automatically generated by fMRIPrep with the express intention that users should copy and paste this text into their manuscripts *unchanged*. It is released under the [CC0](https://creativecommons.org/publicdomain/zero/1.0/) license.
